# Ancestral diversity improves discovery and fine-mapping of genetic loci for anthropometric traits - the Hispanic/Latino Anthropometry Consortium

**DOI:** 10.1101/2021.05.27.445969

**Authors:** Lindsay Fernández-Rhodes, Mariaelisa Graff, Victoria L. Buchanan, Anne E. Justice, Heather M. Highland, Xiuqing Guo, Wanying Zhu, Hung-Hsin Chen, Kristin L. Young, Kaustubh Adhikari, Nicholette (Palmer) Allred, Jennifer E. Below, Jonathan Bradfield, Alexandre C. Pereira, LáShauntá Glover, Daeeun Kim, Adam G. Lilly, Poojan Shrestha, Alvin G. Thomas, Xinruo Zhang, Minhui Chen, Charleston W. K. Chiang, Sara Pulit, Andrea Horimoto, Jose E. Krieger, Marta Guindo-Martinez, Michael Preuss, Claudia Schumann, Roelof A.J. Smit, Gabriela Torres-Mejía, Victor Acuña-Alonzo, Gabriel Bedoya, Maria-Cátira Bortolini, Samuel Canizales-Quinteros, Carla Gallo, Rolando González-José, Giovanni Poletti, Francisco Rothhammer, Hakon Hakonarson, Robert Igo, Sharon G Adler, Sudha K. Iyengar, Susanne B. Nicholas, Stephanie M. Gogarten, Carmen R. Isasi, George Papnicolaou, Adrienne M. Stilp, Qibin Qi, Minjung Kho, Jennifer A. Smith, Carl Langfeld, Lynne Wagenknecht, Roberta Mckean-Cowdin, Xiaoyi Raymond Gao, Darryl Nousome, David V. Conti, Ye Feng, Matthew A. Allison, Zorayr Arzumanyan, Thomas A. Buchanan, Yii-Der Ida Chen, Pauline M. Genter, Mark O. Goodarzi, Yang Hai, Willa Hsueh, Eli Ipp, Fouad R. Kandeel, Kelvin Lam, Xiaohui Li, Jerry L. Nadler, Leslie J. Raffel, Kaye Roll, Kevin Sandow, Jingyi Tan, Kent D. Taylor, Anny H. Xiang, Jie Yao, Astride Audirac-Chalifour, Jose de Jesus Peralta Romero, Fernando Hartwig, Bernando Horta, John Blangero, Joanne E. Curran, Ravindranath Duggirala, Donna E. Lehman, Sobha Puppala, Laura Fejerman, Esther John, Carlos Aguilar-Salinas, Noël P. Burtt, Jose C. Florez, Humberto García-Ortíz, Clicerio González-Villalpando, Josep Mercader, Lorena Orozco, Teresa Tusié, Estela Blanco, Sheila Gahagan, Nancy J. Cox, Craig Hanis, Nancy F. Butte, Shelley A. Cole, Anthony G. Commuzzie, V. Saroja Voruganti, Rebecca Rohde, Yujie Wang, Tamar Sofer, Elad Ziv, Struan F.A. Grant, Andres Ruiz-Linares, Jerome I. Rotter, Christopher A. Haiman, Esteban J. Parra, Miguel Cruz, Ruth J.F. Loos, Kari E. North

**Affiliations:** Department of Biobehavioral Health, Pennsylvania State University, University Park, PA, USA 16802; Department of Epidemiology, Gillings School of Global Public Health, University of North Carolina at Chapel Hill, Chapel Hill, NC, USA 27599; Department of Biomedical and Translational Informatics, Geisinger Health System, Danville, PA, USA 17822; The Institute for Translational Genomics and Population Sciences, The Lundquist Institute for Biomedical Innovation at Harbor-University of California Los Angeles Medical Center, Torrance, CA, USA 90502; Vanderbilt Genetics Institute, Division of Genetic Medicine, Department of Medicine, Vanderbilt University Medical Center, Nashville, TN, USA 37232; School of Mathematics and Statistics, Faculty of Science, Technology, Engineering and Mathematics, The Open University, Milton Keynes, United Kingdom MK7 6AA; Center for Diabetes Research, Center for Genomics and Personalized Medicine Research, Wake Forest School of Medicine, Winston-Salem, NC, USA 27101; Center for Applied Genomics, Division of Human Genetics, The Children’s Hospital of Philadelphia, Philadelphia, PA, USA 19104; Laboratory of Genetics and Molecular Cardiology, Heart Institute, University of São Paulo, Brazil 05508-220; Department of Sociology, University of North Carolina at Chapel Hill, Chapel Hill, NC, USA 27599; Carolina Population Center, University of North Carolina at Chapel Hill, Chapel Hill, NC, USA 27599; Division of Pediatric and Public Health, Adams School of Dentistry, University of North Carolina at Chapel Hill, Chapel Hill, NC, USA 27599; Center for Genetic Epidemiology, Department of Preventive Medicine, Keck School of Medicine, University of Southern California, Los Angeles, CA, USA 90033; Department of Quantitative and Computational Biology, University of Southern California, Los Angeles, CA, USA 90007; Vertex Pharmaceuticals, Oxford, United Kingdom W2 6BD; The Charles Bronfman Institutes for Personalized Medicine, Icahn School of Medicine at Mount Sinai, New York, NY, USA 10029; The Novo Nordisk Center for Basic Metabolic Research, University of Copenhagen, Copenhagen, Denmark 2200; Hasso Plattner Institute, University of Potsdam, Digital Health Center, Potsdam, Germany 14482; Department of Research in Cardiovascular Diseases, Diabetes Mellitus, and Cancer, Population Health Research Center, National Institute of Public Health, Cuernavaca, Morelos, Mexico 62100; National Institute of Anthropology and History, Mexico City, Mexico 06600; Molecular Genetics Investigation Group, University of Antioquia, Medellín, Colombia 1226; Department of Genetics, Federal University of Rio Grande do Sul, Porto Alegre, Brazil 90040- 060; Population Genomics Applied to Health Unit, the National Institute of Genomic Medicine and the Faculty of Chemistry at the National Autonomous University of Mexico, Mexico City, Mexico 04510; Research and Development Laboratories, Faculty of the Sciences and Philosophy, Peruvian University Cayetano Heredia, Lima, Peru 15102; Patagonian Institute of the Social and Human Sciences, Patagonian National Center, Puerto Madryn, Argentina U9120; Institute of High Studies, University of Tarapacá, Arica, Chile 1000000; Department of Population and Quantitative Health Sciences, Case Western Reserve University, Cleveland, OH, USA 44106; Division of Nephrology and Hypertension, Harbor-University of California Los Angeles Medical Center, Torrance, CA, USA 90502; Department of Medicine, David Geffen School of Medicine at University of California, Los Angeles, CA, USA 90095; Department of Biostatistics, University of Washington, Seattle, WA, USA 98195; Department of Epidemiology and Population Health, Albert Einstein College of Medicine, Bronx, NY, USA 10461; National Heart, Lung and Blood Institute, Bethesda, MD, USA 20892; Department of Epidemiology, School of Public Health, University of Michigan, Ann Arbor, MI, USA 48109; Department of Biostatistical Sciences, Wake Forest School of Medicine, Winston-Salem, NC USA 27101; Division of Public Health Sciences, Wake Forest School of Medicine, Winston-Salem, NC USA 27101; Department of Preventive Medicine, Keck School of Medicine, University of Southern California, Los Angeles, CA, USA 90032; Department of Opthalmology and Visual Sciences, Department of Biomedical Informatics, Division of Human Genetics, The Ohio State University, Columbus, OH, USA 43210; Department of Family Medicine and Public Health, University of California, San Diego, CA, USA 92161; Department of Physiology and Biophysics, Keck School of Medicine of USC, Los Angeles, CA USA 90033; Department of Medicine, Division of Endocrinology, The Lundquist Institute for Biomedical Innovation at Harbor-UCLA Medical Center, Torrance, CA, USA 90502; Division of Endocrinology, Diabetes, and Metabolism, Department of Medicine, Cedars-Sinai Medical Center, Los Angeles, CA, USA 90048; Department of Internal Medicine, The Ohio State University Wexner Medical Center, Columbus, OH, USA 43210; Department of Translational Research & Cellular Therapeutics, Beckman Research Institute of City of Hope, Duarte, CA USA 91010; New York Medical College, School of Medicine, Valhalla, NY, USA 10595; Division of Genetic and Genomic Medicine, Department of Pediatrics, University of California, Irvine, CA, USA 92697; Research and Evaluation Branch, Kaiser Permanente of Southern California, Pasadena, CA, USA 91101; Medical Research Unit in Biochemistry, Specialty Hospital, National Medical Center of the Twenty-First Century, Mexican Institute of Social Security, Mexico City, Mexico 06725; Postgraduate Program in Epidemiology, Federal University of Pelotas, Pelotas, Brazil 96010- 610; Department of Human Genetics and South Texas Diabetes and Obesity Institute, School of Medicine, University of Texas Rio Grande Valley, Brownsville and Edinburg, TX, USA 78520 and 78539; Department of Medicine, School of Medicine, University of Texas Health San Antonio, San Antonio, TX, USA 78229; Department of Internal Medicine, Wake Forest University, Winston-Salem, NC, USA 27109; Department of Public Health Sciences, School of Medicine, and the Comprehensive Cancer Center, University of California Davis, Davis, CA, USA 95616; Departments of Epidemiology & Population Health and Medicine-Oncology, Stanford University School of Medicine, Stanford, CA, USA 94305; Division of Nutrition, Salvador Zubirán National Institute of Health Sciences and Nutrition, Mexico City, Mexico 14080; Programs in Metabolism and Medical and Population Genetics, Broad Institute of the Massachusetts Institute of Technology and Harvard, Cambridge, MA, USA 02142; Department of Medicine, Harvard Medical School, Boston, MA, USA 02115; Diabetes Unit and Center for Genomic Medicine, Massachusetts General Hospital, Boston, MA, USA 02114; Laboratory of Immunogenomics and Metabolic Diseases, National Institute of Genomic Medicine, Mexico City, Mexico 14610; Center for Diabetes Studies, Research Unit for Diabetes and Cardiovascular Risk, Center for Population Health Studies, National Institute of Public Health, Mexico City, Mexico 14080; Molecular Biology and Medical Genomics Unity, Institute of Biomedical Research, the National Autonomous University of Mexico and the Salvador Zubirán National Institute of Health Sciences and Nutrition, Mexico City, Mexico 14080; Center for Community Health, Division of Academic General Pediatrics, University of California at San Diego, San Diego, CA, USA 92093; University of Texas Health Science Center at Houston, Houston, TX, USA 77030; United States Department of Agriculture, Agricultural Research Service, The Children’s Nutrition Research Center, and the Department Pediatrics, Baylor College of Medicine, Houston, TX, USA 77030; Population Health Program, Texas Biomedical Research Institute, San Antonio, TX, USA 78227; The Obesity Society, Silver Spring, MD, USA 20910; Department of Nutrition, University of North Carolina Nutrition Research Institute, University of North Carolina, Kannapolis, NC, USA 28081; Division of Sleep and Circadian Disorders, Brigham and Women’s hospital, Boston MA, USA 02115; Division of General Internal Medicine, Department of Medicine, Helen Diller Family Comprehensive Cancer Center, Institute for Human Genetics, University of California, San Francisco, San Francisco, CA, USA 94115; Ministry of Education Key Laboratory of Contemporary Anthropology and Collaborative Innovation Center of Genetics and Development, School of Life Sciences and Human Phenome Institute, Fudan University, Shanghai, China 200438; Department of Genetics, Evolution and Environment, and Genetics Institute of the University College London, London, UK WC1E 6BT; Laboratory of Biocultural Anthropology, Law, Ethics, and Health, Aix-Marseille University, Marseille, France 13385; Department of Anthropology, University of Toronto- Mississauga, Mississauga, ON, Canada L5L 1C6; Carolina Center for Genome Sciences, University of North Carolina at Chapel Hill, Chapel Hill, NC, USA 27514

## Abstract

Hispanic/Latinos have been underrepresented in genome-wide association studies (GWAS) for anthropometric traits despite notable anthropometric variability with ancestry proportions, and a high burden of growth stunting and overweight/obesity in Hispanic/Latino populations. This address this knowledge gap, we analyzed densely-imputed genetic data in a sample of Hispanic/Latino adults, to identify and fine-map common genetic variants associated with body mass index (BMI), height, and BMI-adjusted waist-to-hip ratio (WHRadjBMI). We conducted a GWAS of 18 studies/consortia as part of the Hispanic/Latino Anthropometry (HISLA) Consortium (Stage 1, n=59,769) and validated our findings in 9 additional studies (HISLA Stage 2, n=9,336). We conducted a trans-ethnic GWAS with summary statistics from HISLA Stage 1 and existing consortia of European and African ancestries. In our HISLA Stage 1+2 analyses, we discovered one novel BMI locus, as well two novel BMI signals and another novel height signal, each within established anthropometric loci. In our trans-ethnic meta-analysis, we identified three additional novel BMI loci, one novel height locus, and one novel WHRadjBMI locus. We also identified three secondary signals for BMI, 28 for height, and two for WHRadjBMI. We replicated >60 established anthropometric loci in Hispanic/Latino populations at genome-wide significance—representing up to 30% of previously-reported index SNP anthropometric associations. Trans-ethnic meta-analysis of the three ancestries showed a small-to-moderate impact of uncorrected population stratification on the resulting effect size estimates. Our novel findings demonstrate that future studies may also benefit from leveraging differences in linkage disequilibrium patterns to discover novel loci and additional signals with less residual population stratification.

## INTRODUCTION

A complex interplay between political, social, and economic factors has led to an increasing obesogenic global environment. In this modern context, many low- to middle- income nations have experienced a rapid transition from under-nutrition and growth stunting to over- nutrition and obesity.^1^ Moreover, population-based surveys from 1975-2002 show that there is an inverse ecologic relationship between the prevalence of growth stunting and the prevalence of overweight seen among preschool children (0-5 years of age) in Latin America.^2^ Growth stunting of preschool children ranges from relatively rare (7%) in the Caribbean to notably common (20%) in Central America. Moreover, it is a risk factor for overweight/obesity independent of a child’s socioeconomic status.

In Latin America, by 2016 35% of the total population was overweight [body mass index (BMI) 25 to <30 kg/m^2^] and another 23% was living with obesity.^3^ In Mexico, more than 71% of adults are currently overweight;^4^ it is projected that by 2050 only 12% of men and 9% of women will have a healthy weight (BMI <25 kg/m^2^). In a recent study in Argentina, Chile, and Uruguay, the prevalence of obesity was 36%, but when using waist circumference as a measure of central obesity, it was far higher (53%).^5^ Within each of these populations, there are also disparities in obesity by sex and education.

Race, ethnicity, and ancestry may play a role in anthropometric-related health disparities in Latin American. Previous studies have described the historical contexts leading to admixture in Latin American populations^6; 7^ as characterized by highly diverse (variable) ancestral proportions^8–10^ from any of the following regions: the Americas, Europe, Africa and East Asia.^11–, 16^ In fact, proportion of Native American ancestry is associated with numerous biomedical traits, like obesity-related traits, and is most strongly associated with height.^17; 18^ Height is inversely associated with proportion of Native American ancestry, even after taking into account the fact that globally over time populations have become taller due to mainly non-genetic nutritional factors.^16^ The ultimate drivers of this association remain to be elucidated; it is possible that genetic factors and/or socio-economic factors strongly associated with Native American ancestry could be responsible for these findings. Recent studies are starting to provide relevant insights on this topic. As an example, a recent genome-wide association study (GWAS) in Peru^19^ identified a missense variant in the *FBN1* gene (rs200342067) that has the largest effect size so far described for common height-associated variants in human populations (each copy of the minor allele reduces height by 2.2 cm). In the 1000 Genomes Project samples, rs200342067 is only present in two American samples (MXL: 0.78% and PEL: 4.12%), and yet the authors reported that this missense variant shows subtle evidence of positive selection in the Peruvian population.^19^

Obesity in Latin America has quickly surpassed the levels previously seen only among adults of high-income nations, like Canada and the United States (US). In Canada the number of people reporting Latin American origins grew by 83% from the 2001 census^20^ relative to the 2016 census,^21^ representing 1.3% of the total Canadian population. In the US, both the population size and diversity in national origins (backgrounds) of US Hispanic/Latinos have been increasing over the past several decades.^22^ If past demographic trends continue, 24% of the US adult population will identify as Hispanic/Latino by 2065.^22^ Obesity-related financial costs in the US are projected to double every decade to ∼$900 billion by 2030.^23; 24^ US Hispanic/Latino adults and their children/adolescents face a greater burden of obesity than their non-Hispanic white counterparts.^25–28^ There is a need to study Hispanic/Latino populations in order to address these disparities.^28; 29^

Given the unique historical and recent demographic shifts occurring across the Americas, there is a clear need to also understand the role that Native American or other under- studied components of admixture have on the genetic architecture of anthropometric traits in Hispanic/Latinos, and its relationship with risk of downstream poor health outcomes. Yet, to date no large-scale GWAS of anthropometric traits have been conducted among Hispanic/Latino populations. Here, we perform the largest genomic study to date of anthropometric traits, including BMI, height, and waist-to-hip ratio adjusted for BMI (WHRadjBMI) in Hispanic/Latino populations to describe what might be novel loci or signals in established loci in this population by sex and life stage.

## MATERIALS AND METHODS

### Hispanic/Latino Study Samples

The Hispanic/Latino Anthropometry (HISLA) Consortium is comprised of 27 studies/ consortia of adult participants. First, HISLA Stage 1 includes 17 studies and one consortium (Consortium for the Analysis of the Diversity and Evolution of Latin America, CANDELA^18^) collectively representing up to 59,771 adults, depending on the trait, from Brazil, Chile, Colombia, Mexico, Peru, or the US with self-reported heritage from across Spanish-speaking Latin America, or Native American heritage, primarily Pima and Zuni^30^ (**Table S1**). HISLA Stage 2 includes nine studies with up to 10,538 adults from across Spanish-speaking Latin America or with related heritage and living in the US (**Table S1**).

This study was approved by the Institutional Review Boards of the University of North Carolina at Chapel Hill, and all contributing studies had received prior Institutional Review Boards approval for each study’s activities.

### Anthropometric Traits

BMI is a commonly derived index of obesity risk and is calculated as the ratio of body weight to height squared (kg/m^2^). Adult height was measured/self-reported using either metric units, or US units and then converted to meters. Waist-to-hip ratio (WHR) is used to capture central fat deposition, and it is derived from the circumference of the waist at the umbilicus compared to the circumference of the hip at the maximum protrusion of the gluteal muscles.

Residuals were calculated by sex and/or case status, adjusting for age, age^2^, and study-specific covariates [e.g., center, principal components of ancestry (PCA)]. For WHR, BMI was also adjusted for when creating the residuals to isolate the central deposition of fat from overall body mass. Residuals were then used to create inverse normalizations of BMI and WHRadjBMI, and z-scores of height (=residual/standard deviation for all residuals). In family-based studies the residuals were calculated in women and men together, adjusting for age and sex and other study covariates including PCs. Descriptive statistics on the covariates and anthropometric measures of are provided for each study’s analytic sample in **Table S2**. Only one family-based study in Stage 1 and two non-family based studies in Stage 2 (GOLDR 0.3% <18 years, and HTN-IRS 3.9%) included a small subset of adolescents aged 15-17 years, each less than 5% of the total sample. All other study samples included individuals 18-98 years of age.

### Childhood/Adolescence Study Samples, Anthropometric Traits, and Obesity

We assembled an independent sample of children/adolescents with anthropometrics, from three studies from the US, Mexico and Chile (**Table S3**). The distribution of covariates and anthropometrics of the samples of children/adolescents in each analysis are described in **Table S4.** First, childhood/adolescent obesity was defined as having a ≥95^th^ BMI-for-age percentile versus ≤50^th^ BMI-for-age percentiles, based on the Centers for Disease Control and Prevention growth curves,^31^ as done in previous analyses of childhood obesity.^32^ We used these two analyses to look up novel BMI and height findings from our adult HISLA meta-analysis and our trans-ethnic analyses. This resulted in 1,814 children/adolescents aged 2-18 years for this case-control analysis (**Tables S3-4**). Second, BMI and height-for-age z-scores were calculated in children/adolescents aged 5-18 years from the US and Chile (**Table S4**) based on the more international reference growth curves from the World Health Organization.^33^ In Viva la Familia, a family-based study,^34^ these residuals were calculated adjusting for sex in the combined sample. The resulting BMI and height-for-age z-scores were available for 1,914 and 1,945 children/adolescents, respectively.

### SNP Imputation and Statistical Analyses

We generated autosomal genome-wide imputed data based on 1000 Genomes Phase 1 and 3 references, with the exception of two studies that contributed Exomechip and MetaboChip (Illumina, Inc.; San Diego, CA) genotypes and one study that blended genotypes from multiple platforms (**Tables S5-6**). PCA analyses were conducted in each study to capture the main components of genetic ancestry from the Americas, Europe, Africa, and Asia. Studies with samples from related individuals accommodated this non-independence by projecting their principal component analysis from the reference to the study sample, and by accounting for relatedness using either generalized estimating equations^35^ or mixed linear models.^10; 36^ Assuming an additive genetic model, we tested the association of over 20 million autosomal variants on our traits, accounting for all trait or study-specific covariates (e.g., center, PCA).

### Meta-Analyses of HISLA Stage 1+2

The studies of the HISLA Consortium were meta-analyzed in two stages, including discovery (Stage 1) and validation (Stage 2). Stage 1 included a total sample of 59,771 individuals with data on BMI, 56,161 with height, and 42,455 with WHRadjBMI. All Stage 1 studies/consortia provided full genome-wide analysis results. All SNPs that met our significance criteria were brought forward for validation in Stage 2, which included 10,538 individuals with data on BMI, 8,110 with height, and 4,393 with WHRadjBMI. All reported association results passed our quality control criteria; i.e., variants with low quality (info score <0.4 or Rsq<0.3), minor allele count (MAC) <5, or sample size <100 were removed. We meta-analyzed effects across all studies using a fixed-effect inverse variance weighted meta-analysis with genomic control in METAL.^37^ Given the unique patterns of admixture and ancestry represented by the Brazilian or Native American samples, we conducted sensitivity analyses in Stage 1 studies (i.e., comparing the inclusion and exclusion of the Baependi Heart Study, 1982 Pelotas Birth Cohort Study, and Family Investigation of Nephropathy and Diabetes substudy of individuals of Pima and Zuni heritage) to assess the influence of the three studies on the meta-analysis results. CANDELA was retained in all analyses as <10% of the consortium’s samples came from Brazil, primarily originating from the South of Brazil with wide-spread European heritage with a lesser extent Native American or African admixture.^18^

Regional plots of all GWAS-significant HISLA Stage 1 findings were plotted using LocusZoom (https://locuszoom.org). From Stage 1, we selected lead variants that met genome-wide significance (P<5×10^-8^) that were independent of each other for replication. In cases where Stage 2 studies did not have the lead variant, we selected two proxies per lead variant with an r^2^≥0.9 using 1000 Genomes AMR linkage disequilibrium (LD). Stage 2 studies provided a list of the requested lead variants and/or their proxies from Stage 1 for validation. Stage 2 studies were meta-analyzed and subsequently combined with Stage 1 using METAL^25^. Effect heterogeneity was assessed through I^2^ across all 27 HISLA adult studies/consortia by entering each study separately into the meta-analysis, irrespective of stage. The characteristics of the final SNP array data used in the HISLA adult studies and the children/adolescent Hispanic/Latino studies are summarized separately in **Tables S5-6**.

### Meta-Analyses of HISLA Stage 1 with Other Ancestral Consortia

In addition to a Hispanic/Latino only meta-analysis, we combined the HISLA Stage 1 meta-analysis with data from previous large-scale GWAS meta-analyses from European (the Genetic Investigation of Anthropometric Traits, GIANT, Consortium^38–40^, N ∼ 300,000) and/or African (the African Ancestry Anthropometry Genetics Consortium, AAAGC^41^, N ∼ 50,000) descent populations. We used fixed-effect inverse variance weighted meta-analytic techniques in METAL to generate our trans-ethnic meta-analysis.^37^ We validated our potentially novel BMI, height^42^ and WHRadjBMI^43^ findings from this trans-ethnic meta-analysis in either our independent sample of Hispanic/Latino children/adolescents or the British subsample GWAS of the United Kingdom Biobank (UKBB). Regional plots of these analyses of all potentially novel trans-ethnic findings are shown in the supplement (**Figures S7-52**).

### Thresholds for Conditional Signals, Discovery, Validation and Transferability

We conducted approximate conditional analyses using the Genome-wide Complex Trait Analysis (GCTA, version 1.93.1) software. For the HISLA analyses, we used our Stage 1 discovery results with the Hispanic Community Health Study/Study of Latinos (HCHS/SOL) as the LD reference dataset. For the approximate conditional trans-ethnic analyses, we used our trans-ethnic results from HISLA Stage 1, AAAGC, or GIANT and a trans-ethnic LD reference dataset of Europeans and Africans from the Atherosclerosis Risk in Communities (ARIC) cohort, and Hispanic/Latinos from HCHS/SOL, which was representative of the ancestry distribution of our meta-analysis. In both conditional analyses (HISLA only and trans-ethnic results), we first identified all independent SNPs using the --cojo-slct command. Then, we conditioned each of these independent SNPs on all known SNPs up through December 2019 (BMI^38; 41; 42; 44–59^, Height^39; 42; 51; 54; 59; 60^, WHRadjBMI^40; 41; 43; 48; 50; 58; 59; 61–65^) within 10Mb of the index SNP. The trans-ethnic meta-analysis results with a *P-value*<5×10^-8^ after conditioning on known SNPs were taken forward for validation in the British subsample of the UKBB.

SNP associations were then defined as either newly discovered or established, depending on their location. An established locus was defined as a SNP association within ±500 kb of at least one previously identified index SNP, otherwise the association was considered a newly-discovered locus.

We designated our Hispanic/Latino SNP-associations within either newly-discovered or established loci as novel if they met the following criteria: 1) were associated at *P-value*<5×10^-8^ in HISLA Stage 1 and directionally consistent in Stage 2, and 2) the addition of Stage 2 samples improved the estimated Stage 1+2 meta-analysis. For the trans-ethnic analyses these criteria were as follows: 1) were associated at *P-value*<5×10^-8^ in the combined HISLA, AAAGC and GIANT meta-analysis, and 2) were both directionally consistent and associated at *P-value*<5×10^-2^ in the subsample of Hispanic/Latino children/adolescents or in the British subsample GWAS from the UKBB.

Novel Hispanic/Latino SNP effects were considered to transfer to Hispanic/Latino children/adolescents, or to African or European ancestry adults, if they were 1) directionally consistent, 2) associated at *P-value*<5×10^-2^, and 3) had a heterogeneity of I^2^<75% in either the Hispanic/Latino children/adolescent lookups, or either 1) the AAAGC or 2) the GIANT adult GWAS results. Conversely, SNP effects of variants previously associated with anthropometric traits in non-Hispanic/Latino populations (i.e., index published SNPs) were considered to be transferable (generalizable) to Hispanic/Latinos if they were 1) directionally consistent, 2) displayed a *P-value*<5×10^-2^, and 3) had little to moderate effect heterogeneity (I^2^<75%) in Stage 1.

### Fine-Mapping Methods

We used FINEMAP^66^ for fine-mapping analyses of the newly-discovered loci identified as part of the HISLA Stage 1 meta-analysis or trans-ethnic meta-analysis, and in established loci. For the established loci, we included index SNP-associations published as of April 2018 (BMI^38; 41; 44; 46–48; 50; 52; 55–58^, Height^39; 54; 60^, WHRadjBMI^40; 41; 48; 50; 67^) prior to the publications with the UKBB results.^42; 43^ We used a 1Mb region subset of the summary statistics from the Stage 1 meta-analyses and HCHS/SOL^10^ unrelated sample set (N ∼ 7,670) to calculate the LD for each locus.

For trans-ethnic fine-mapping of the novel loci and signals identified in the trans-ethnic meta-analysis of HISLA, AAAGC, and GIANT, we used a 1Mb region defining each locus using the summary statistics of the given meta-analysis. We calculated the LD for Hispanic/Latino samples using the HCHS/SOL^10^ unrelated sample (N ∼ 7,670). For African and European ancestry samples, we calculated the LD using the ARIC unrelated sample that included self-reported African ancestry (N ∼ 2,800) and European ancestry (N ∼ 9,700). We weighted the LD matrices by the GWAS sample sizes for each trait (HISLA range: ∼42,400-56,100; AAAGC: 20,300-42,700; GIANT: 210,000-330,000).

All regions allowed up to a maximum of 10 causal variants. The cumulative 95^th^% credible set was calculated from the estimated posterior probabilities. Convergence failed for three regions (lead SNPs: rs2902635, rs6900530, and rs4425978, all in known height loci) using the stochastic approach. For these three regions, we used the conditional approach to determine number of causal variants.

### Gene Expression & Other Bioinformatic Analyses

To assess for potential validation of our potentially novel or validated HISLA hits, we performed association analyses of measured whole blood gene expression in 606 individuals from Cameron County Hispanic Cohort.^68^ RNA sequencing was conducted using 150bp paired-end reads on the Illumina NovaSeq 6000 by Vanderbilt Technologies for Advanced Genomics. Initial sequencing quality was checked by FastQC.^69^ STAR-2.7.8a was applied to align sequencing reads alignment to the human genome reference (UCSC, hg38),^70^ and the aligned reads were assigned to genes using featureCounts.^71^ We excluded either samples with less than 15M total aligned reads, a rate of successful alignment of less than 20%, or less than 15M total assigned reads. The sequencing library size was normalized using DESeq2^72^ and read counts were transformed using variance stabilizing transformations (vst in DESeq2 package). We performed expression quantitative trait loci (eQTL) analysis with our top HISLA SNP findings, by modeling SNP dosages (exposure) in a linear regression of gene expression levels (outcomes), for each gene within the 1 MB interval around each lead SNP. We inverse normalized the gene expression levels and adjusted for age, sex, and three principal components to capture population substructure. Bonferroni correction for each region varied according to the number of SNPs tested.

To gain further insight into the possible functional role of the identified variants and to assess their relevance to other phenotypes, we conducted bioinformatics queries of our potentially novel loci and novel signals within known loci in multiple publicly available databases, including PhenoScanner,^73^ RegulomeDB,^74^ Haploreg,^75^ UCSC GenomeBrowser,^76^ and GTEx.^77^

### Trans-Ethnic Findings to Account for Population Structure in Previous GWAS

To quantify the impact of population stratification, we computed the correlation between PC loadings and beta effects estimated from GWAS. We first conducted PCA analysis on the four European populations (CEU, GBR, IBS, and TSI) from 1000 Genomes. We excluded the FIN (Finnish in Finland) population because of its known unique demographic history.^38^ We only used biallelic SNPs with minor allele frequency (MAF) > 5% in the four European populations, and then pruned them by both distance and LD using PLINK 1.9.^78^ Specifically, we pruned the dataset such that no two SNPs were closer than 2 kb, and then pruned using a 50 SNP LD window (moving in steps of 5 SNPs), such that no SNPs had r^2^>0.2. We further removed SNPs in regions of long-range LD.^79^ PCA was performed on the remaining SNPs using Eigensoft version 7.2.1(https://github.com/DReichLab/EIG/archive/v7.2.1.tar.gz).

We performed linear regressions of individual PC values on the allelic genotype count for each polymorphic variant in the four European populations from 1000 Genomes and used the resulting regression coefficients as the estimate of the variant’s PC loading. For each PC, we then computed Pearson correlation coefficients of PC loadings and effect sizes (of variants with MAF>1%) from each GWAS summary statistics. We estimated *P-values* based on Jackknife standard errors, by splitting the genome into 1,000 blocks with an equal number of variants. If there was significant correlation in either the GIANT dataset or the HISLA Stage 1, AAAGC and GIANT trans-ethnic meta-analysis, we then further evaluated the improvement of bias due to stratification in trans-ethnic meta-analysis by comparing the correlation coefficients in the trans-ethnic meta-analysis with those in GIANT. Restricting to variants shared between GIANT and the trans-ethnic meta-analysis, we computed their difference in correlation coefficients of PC loadings and effect sizes, and estimated *P-values* again based on Jackknife standard errors from 1,000 equal sized blocks.

## RESULTS

### One Novel BMI Locus Discovered and Validated in Hispanic/Latino Adults

The first goal of this study was to conduct a genome-wide meta-analysis of anthropometric traits in Hispanic/Latino adults to identify novel loci in an under-studied population (**Figure 1**). All regional plots of all GWAS-significant HISLA Stage 1 findings are shown in the supplement **(Figures S1-6).** No novel loci were identified in all samples combined. Yet, when excluding the Brazilian or Native American samples from Stage 1, we discovered one locus for adult BMI at *PAX3* on chromosome 2 in the HISLA Stage 1 sample (**Table S7**), and we validated this locus in HISLA Stage 2 (**Table 1**). The lead SNP, rs994108, is in moderate LD (rs7559271, r^2^=0.46, D’=1.0 in 1000 Genomes phase 3 AMR) (**Figure 2**) and lies on the same haplotype as a SNP reported to influence facial morphology, including position of the nasion (the deepest point on the nasal bridge where the nose meets the forehead) in Europeans^80^ and Hispanic/Latino^81^ descent individuals. Other *PAX3* variants in lower LD with the lead SNP have also been associated with nasion position,^82^ monobrow, and male-pattern baldness.^83; 84^ *PAX3* is a well-known transcription factor in normal embryonic neural crest development and differentiation.^85^ Neural crest cells can give rise to mesenchymal stem cells,^86^ which can in turn give rise to adipocytes;^86–88^ thus, the possible role of *PAX3* in adipogenesis may at least partially explain the association signal with BMI near this gene. Another BMI SNP (rs1505851) near *ARRDC3* on chromosome 5 found at GWAS significance in HISLA Stage 1 (**Table S7, Figure S1**) did not validate in Stage 2 (**Table 1**).

**Figure 1.**
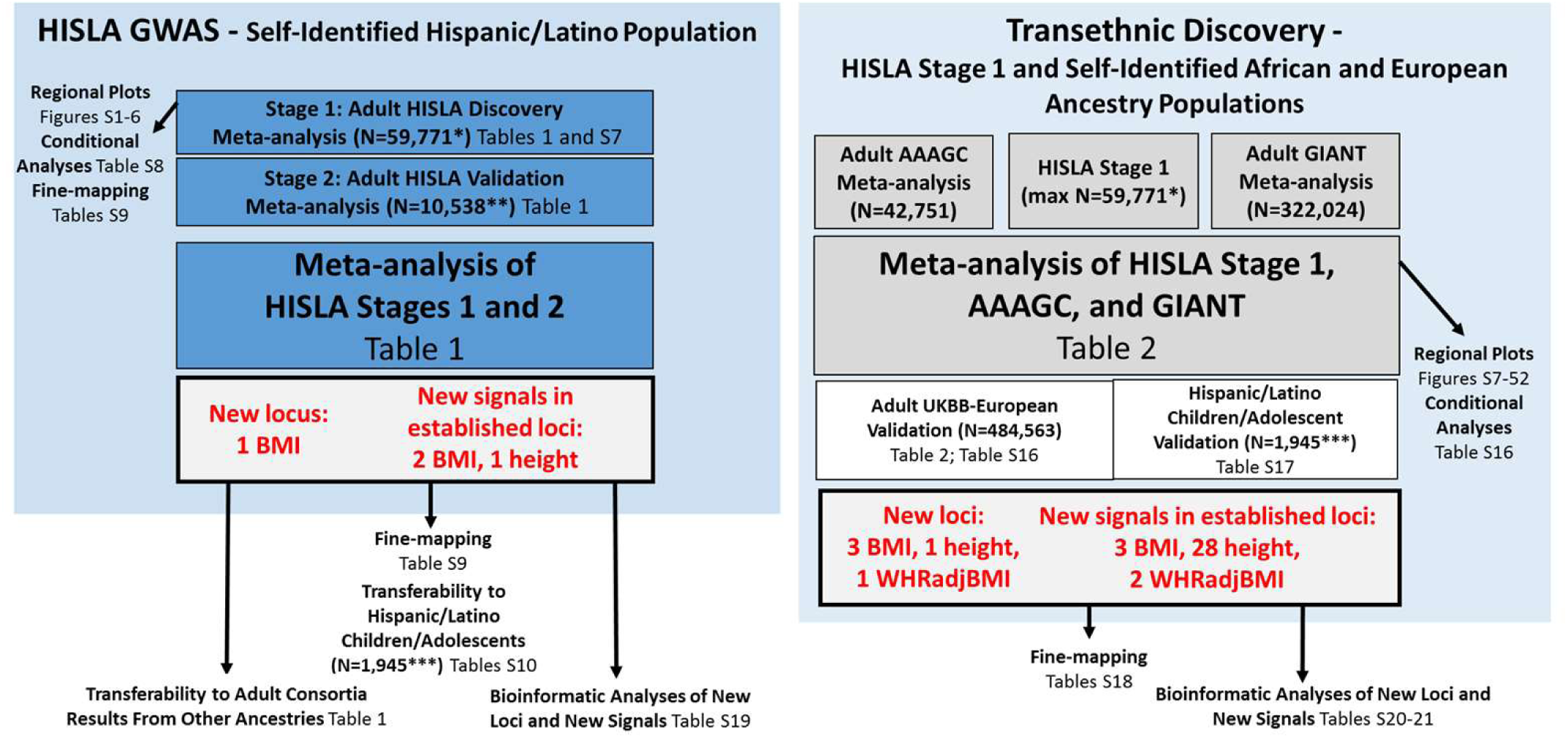
Flowchart of the design and discovery of 6 novel loci and 36 novel signals in known loci in the Hispanic/Latino Anthropometry Consortium (HISLA) Meta-Analysis and the Trans-Ethnic Meta-Analysis of HISLA and Consortia of Other Ancestral Heritages. *Stage 1 maximum sample sizes varied between and 59,771 for BMI, 56,161 for height, and 42,455 for WHRadjBMI (sex combined). **Stage 2 sample sizes varied between 10,538 for BMI, 8,110 for height, and 4,393 for WHRadjBMI (sex combined). Actual sample sizes may vary by SNP. ***The BMI and height-for-age z-score models were conducted using up to 1,914 and 1,945 of children/adolescents, respectively. In contrast, the obesity case-control study compared up to 1,814 children/adolescents who were either ≥95th versus ≤50th BMI-for-age percentiles

**Figure 2.**
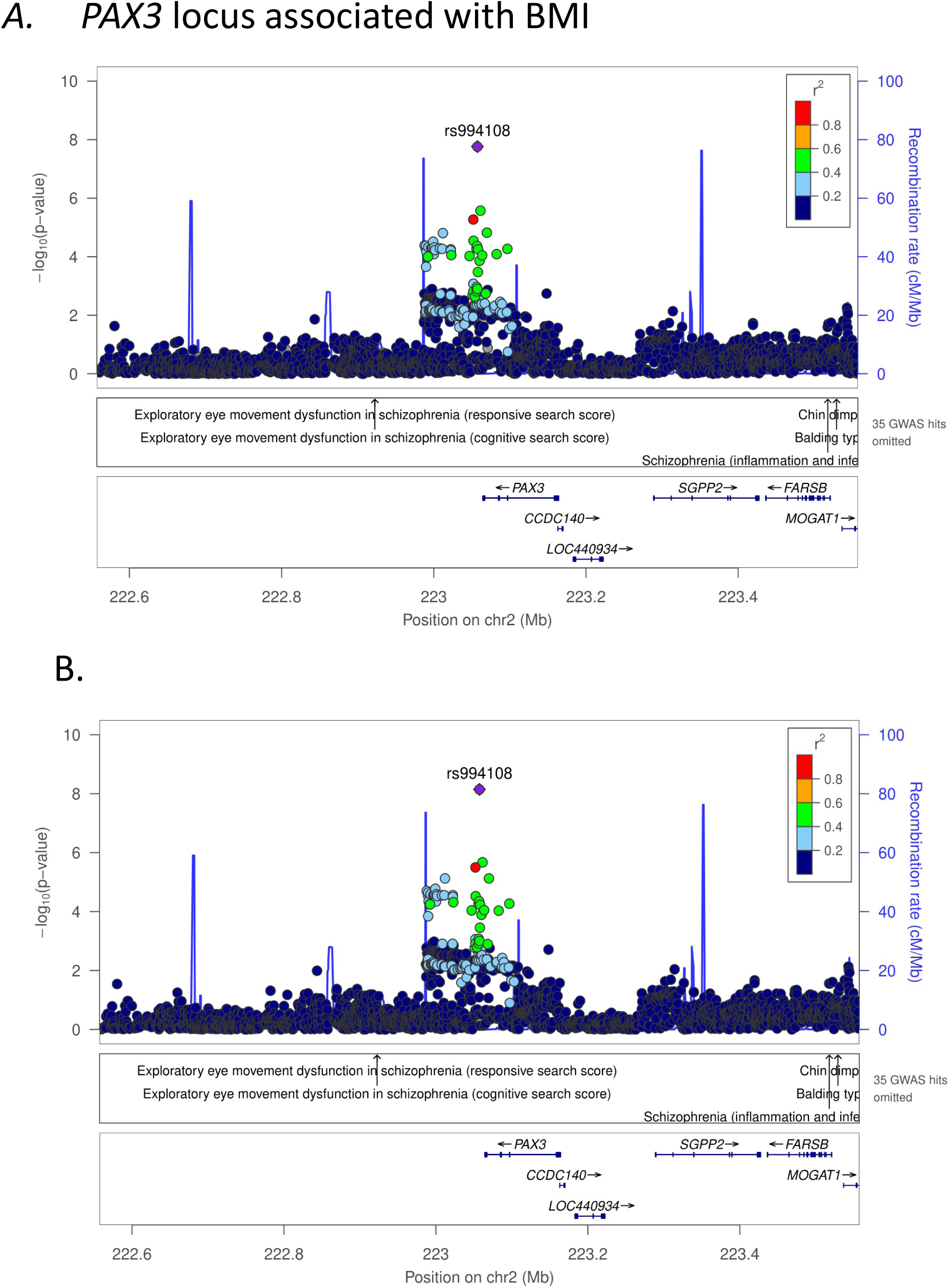
Regional plot, unconditioned (A) and conditioned (B), of the novel PAX3 locus associated with body mass index (BMI) in the Hispanic/Latino Anthropometry Consortium (HISLA), excluding Brazilian and Native American samples. Linkage disequilibrium patterns are based on rs994108 (shown by the purple triangle) from the Hispanic Communities in Health Study/Study of Latinos.

**Table 1.** Potential novel loci and new signals in known loci from the Stage 1: Adult HISLA Discovery combined with results from the Stage 2: Adult HISLA Validation.^1^ In addition, lookup of results of each locus from the AAAGC and GIANT. Abbreviations: Chr - chromosome; EAF - effect allele frequency; HetIsq - heterogeneity I- square; N - sample size; WHRadjBMI - waist to hip ratio adjusted for BMI; AAAGC- African American Anthropometry Genetics Consotrium; GIANT- Genetic Investigation of ANthropometric Traits ^1^ All studies were meta-analyzed using METAL (PMID 20616382), with each study entered individuals into Stage 1+2 analyses.^2^ These BMI and WHRadjBMI analyses did not include Brazilian and/or Native American samples. ^3^ New loci or signals are those that were validated by HISLA stage 2 results that are directionally consistent with Stage 1 and remaining genome-wide significant after meta-analysis with Stage 1. ^4^ Proxy GIANT, rs1573905 (r2= 0.96 AMR)

| Each locus from the AAAGC and GIANT. |  |  |  |  |  |  |  |  |  |  |  |  |  |  |
| --- | --- | --- | --- | --- | --- | --- | --- | --- | --- | --- | --- | --- | --- | --- |
| Trait | Locus Name | SNP rsid | Genomic region | Chr | Position (hg19) | Effect/ Other Alleles | Stage | EAF | Beta | SE | P-value | HetISq | N | Yes/ No |
| <u>Novel loci</u> |  |  |  |  |  |  |  |  |  |  |  |  |  |  |
| BMI<sup>2</sup> | PAX3 | rs994108 | intergenic | 2 | 223057288 | C/A | Stage 1: Discovery | 0.390 | 0.041 | 0.007 | **1.62E-08** | 0 | 43048 | Yes |
|  |  |  |  |  |  |  | Stage 2: Validation | 0.394 | 0.030 | 0.016 | 5.65E-02 | 8 | 9336 |  |
|  |  |  |  |  |  |  | Stage 1 + 2 | 0.389 | 0.038 | 0.006 | **2.19E-09** | 0 | 52384 |  |
|  |  |  |  |  |  |  | AAAGC | 0.526 | 0.007 | 0.007 | 3.26E-01 | 0 | 42751 |  |
|  |  |  |  |  |  |  | GIANT | 0.342 | 0.0001 | 0.004 | 9.81E-01 | - | 233955 |  |
| BMI | ARRDC3 | rs1505851 | intronic | 5 | 90893954 | T/C | Stage 1: Discovery | 0.741 | 0.041 | 0.007 | **2.287E-08** | 0 | 52365 | No |
|  |  |  |  |  |  |  | Stage 2: Validation | 0.709 | 0.005 | 0.017 | 7.62E-01 | 33.3 | 9336 |  |
|  |  |  |  |  |  |  | Stage 1 + 2 | 0.735 | 0.035 | 0.007 | 1.16E-07 | 14.1 | 61701 |  |
|  |  |  |  |  |  |  | AAAGC | 0.307 | 0.027 | 0.008 | 7.00E-04 | 46.6 | 42752 |  |
|  |  |  |  |  |  |  | GIANT | 0.680 | 0 | 0.004 | 7.90E-01 | - | 233999 |  |
| WHRadjBMI - Women only<sup>2</sup> | DOCK2 | rs6879439 | intronic | 5 | 169314869 | C/T | Stage 1: Discovery | 0.520 | 0.060 | 0.010 | **1.02E-08** | 0 | 18591 | No |
|  |  |  |  |  |  |  | Stage 2: Validation | 0.526 | 0.013 | 0.028 | 6.54E-01 | 28.7 | 2747 |  |
|  |  |  |  |  |  |  | Stage 1 + 2 | 0.515 | 0.049 | 0.0093 | 1.57E-07 | 1.9 | 23382 |  |
|  |  |  |  |  |  |  | AAAGC | 0.440 | 0.012 | 0.012 | 3.09E-01 | 0 | 15600 |  |
|  |  |  |  |  |  |  | GIANT | 0.610 | 0.0025 | 0.005 | 6.30E-01 | - | 86317 |  |
| WHRadjBMI - Sex combined | TAOK3 | rs115981023 | intronic | 12 | 118751105 | A/G | Stage 1: Discovery | 0.009 | 0.328 | 0.057 | **1.08E-08** | 44.8 | 19640 | No |
|  |  |  |  |  |  |  | Stage 2: Validation | 0.004 | -0.339 | 0.687 | 6.22E-01 | 0 | 1340 |  |
|  |  |  |  |  |  |  | Stage 1 + 2 | 0.009 | 0.308 | 0.057 | 5.18E-08 | 52 | 20980 |  |
|  |  |  |  |  |  |  | AAAGC | 0.050 | 0.027 | 0.027 | 3.07E-01 | 0 | 15601 |  |
|  |  |  |  |  |  |  | GIANT | 0.002 | no proxy |  |  |  |  |  |
| <u>New signals in known loci</u> |  |  |  |  |  |  |  |  |  |  |  |  |  |  |
| BMI | ADCY5 | rs17361324 | intronic | 3 | 123131254 | T/C | Stage 1: Discovery | 0.280 | 0.042 | 0.008 | **2.60E-08** | 0 | 43333 | Yes |
|  |  |  |  |  |  |  | Stage 2: Validation | 0.269 | 0.035 | 0.018 | 4.70E-02 | 0 | 9035 |  |
|  |  |  |  |  |  |  | Stage 1 + 2 | 0.278 | 0.041 | 0.007 | **2.84E-09** | 0 | 52368 |  |
|  |  |  |  |  |  |  | AAAGC | 0.119 | 0.023 | 0.011 | 3.85E-02 | 0 | 42682 |  |
|  |  |  |  |  |  |  | GIANT | 0.253 | 0.013 | 0.004 | 9.90E-04 | - | 320704 |  |
|  | C6orf106 | rs148899910 | intergenic | 6 | 34232259 | C/G | Stage 1: Discovery | 0.275 | 0.040 | 0.007 | **9.03E-09** | 0 | 54105 | Yes |
|  |  |  |  |  |  |  | Stage 2: Validation | 0.282 | 0.049 | 0.017 | 4.43E-03 | 0 | 9035 |  |
|  |  |  |  |  |  |  | Stage 1 + 2 | 0.276 | 0.041 | 0.006 | **1.24E-10** | 0 | 63140 |  |
|  |  |  |  |  |  |  | AAAGC | 0.316 | -0.016 | 0.008 | 5.02E-02 | 30 | 42750 |  |
|  |  |  |  |  |  |  | GIANT<sup>4</sup> | 0.017 | 0.036 | 0.012 | 3.99E-03 | - | 216522 |  |
| Height | B4GALNT3 | rs215226 | intronic | 12 | 591300 | A/G | Stage 1: Discovery | **0.550** | **-0.032** | **0.005** | **5.53E-09** | 22.1 | 52156 | Yes |
|  |  |  |  |  |  |  | Stage 2: Validation | 0.565 | -0.020 | 0.017 | 2.37E-01 | 19.3 | 6906 |  |
|  |  |  |  |  |  |  | Stage 1 + 2 | 0.551 | -0.031 | 0.005 | **1.98E-09** | 21.8 | 59062 |  |
|  |  |  |  |  |  |  | AAAGC | 0.772 | -0.031 | 0.009 | 8.99E-04 | 24 | 41327 |  |
|  |  |  |  |  |  |  | GIANT | 0.633 | 0.006 | 0.004 | 1.10E-01 | - | 220370 |  |

Abbreviations: Chr - chromosome; EAF - effect allele frequency; HetISq - heterogeneity I-square; N - sample size; WHRadjBMI - waist to hip ratio adjusted for BMI; African American Anthropometry Genetics Consortium AAAGC; Genetic Investigation of Anthropometric Traits (GIANT)
<sup>1</sup> All studies were meta-analyzed using METAL (Willer et al 2010 PMID 20616382), with each study entered individuals into Stage 1+2 analyses.
<sup>4</sup> Proxy GIANT, rs1573905 (r<sup>2</sup> = 0.96 AMR)

We identified two WHRadjBMI loci at *DOCK2* and *TAOK3* at GWAS significance in HISLA Stage 1 after excluding the Brazilian and Native American samples (**Table S7, Figures S2-S3**), and neither met the p-value threshold for replication and in HISLA Stage 2. The *DOCK2* association for WHRadjBMI was observed among women in Stage 1was, however, directionally consistent among women in Stage 2. The *TAOK3* association was led by a low frequency variant (rs115981023) that was not directionally consistent across Stages. rs115981023 exhibited moderate heterogeneity across Stage 1 samples after excluding Brazilian and Native American samples (I^2^=45%), and this heterogeneity remained (I^2^=52%) in the combined meta-analysis of HISLA Stage 1 and 2 samples (**Table 1**).

No potentially novel loci were identified for height in HISLA Stage 1, and the exclusion of the Brazilian and Native American samples did not reveal additional novel height or WHRadjBMI loci.

### Three Novel Signals in Established Loci for BMI and Height Discovered and Validated in Hispanic/Latino Adults

At two established loci for BMI, we identified new signals at *ADCY5* and near *C6orf106,* which has recently been renamed *ILRUN* (**Table S7**). These signals were both independent of any previously published anthropometric findings (**Table S8, Figures S4-5**). We validated these signals in Stage 2 with directional consistency and the combined Stage 1+2 meta-analysis at GWAS significance (**Table 1**). We also identified one new signal for height in an established height locus, *B4GALNT3,* which was independent of the previously reported SNPs for height (**Tables S7-8**, **Figure S6**). We validated this signal in Stage 2 with directional consistency and a Stage 1+2 meta-analysis that was GWAS significant (**Table 1**). In additional gene expression and bioinformatics analyses (**Table S18-20**), we found that each of the three novel signals in an established anthropometric loci is supported by either an eQTL in whole blood in Hispanic/Latino populations (**Table S18**), and/or an in eQTL other tissues from publicly available (non-Hispanic/Latino) datasets, e.g., thyroid, esophagus, artery (**Table S19-20**).

### Fine-Mapping of Novel Adult Hispanic/Latino Anthropometric Findings

We fine-mapped using 1MB regions, the novel *PAX3* locus for BMI and three new signals in known loci discovered and replicated in Stages 1+2 (BMI: *ADCY5* and *C6orf106*; height: *B4GALNT3*; **Table S9**). For the three BMI loci, FINEMAP revealed one potential causal set for each locus at *PAX3, ADCY5,* and *C6orf106* locus. For the *PAX3* locus, only one causal set was proposed and the 95^th^% credible contained only nine plausibly causal SNPs, with lead SNP rs994108 having a very high posterior probability of being causal (0.89, **Table S21**). However, functional annotation of this SNP was unremarkable (**Tables S22-23**). In contrast, for *ADCY5* and *C6orf106,* FINEMAP revealed one causal configuration for each locus but with much greater uncertainty with respect to the likely functional variant given the size of the credible sets, 14 and 22 SNPs in the credible region for *ADCY5* and *C6orf106*, respectively. The posterior probability of the best lead SNP at these loci had relatively low posterior probabilities of being the causal SNP, with the best posterior probabilities of 0.23 for rs17361324 (*ADCY5*), and 0.11 for rs73420913 (*C6orf106*), respectively. Interestingly, however, the best candidate for causality at *PAX3* and *ADCY5* loci were the lead SNPs from the HISLA meta-analysis and for *C6orf106*, the FINEMAP and HISLA SNPs were in tight LD (rs73420913 had an r^2^=0.96 with the lead HISLA SNP rs148899910), providing greater support for the prioritization of these SNPs for functional interrogation. For the *B4GALNT3* locus for height, FINEMAP revealed six causal configurations. Four of the variants (rs11063185, rs215230, rs7303572, and rs11063184 with each configuration each had a posterior probability >0.99 and contained only itself in the 95% credible set. One variant (rs215223) had a posterior probability of 0.93 and thus included two variants in the 95% credible set. The sixth 95% credible set had a lead variant with a posterior probability of 45%, but contained a total of 1621 additional variants all of which had very small posterior probabilities (i.e., ≤0.05).

### Transferability of Adult Novel Loci/Signals from Hispanic/Latinos to Consortia of Other Ancestral Backgrounds

To assess how well the effect estimates are transferable (generalizable) to other populations, we looked up the novel BMI and height findings from Hispanic/Latinos in the AAAGC and GIANT meta-analysis results (**Table 1**). Keeping limitations with respect to sample size, LD, allele frequency, and effect size heterogeneity in mind, we did observe directionally consistent BMI effects at the *PAX3* locus in the other consortia, although without observing nominal significance. The new BMI signals at the *ADCY5* locus (rs17361324) transferred to both AAAGC and GIANT with directional consistency (betas=0.13-0.23) and nominal significance (*P-values*<5×10^-2^). The BMI lead SNP (rs148899910) representing a novel signal near *C6orf106* was available in AAAGC, the signal only appeared to be transferable to GIANT at a proxy SNP (rs1573905, r^2^=0.96-1 in 1000 Genomes AMR and EUR; **Table 1**).

The new signal for height in *B4GALNT3* (rs215226) was directionally consistent and nominally significant in AAAGC. In all cases the effect sizes observed in GIANT and AAAGC were attenuated compared to the effect sizes from HISLA Stage 1.

### Relevance of Novel Hispanic/Latino Anthropometric Loci/Signals from Adults to Childhood/Adolescence

We looked up our novel HISLA findings in Hispanic/Latino children/adolescents using BMI-for-age and height-for-age z-scores, as well as a case-control study of childhood obesity. Two of the three novel BMI signals were directionally consistent with the anticipated effect on the odds of obesity during childhood/adolescence, one of which was nominally significant (rs17361324 at *ADCY5*; *P-value*=2.2×10^-2^). None of the novel HISLA findings generalized at nominal significance with the BMI/height-for-age z-score, but were directionally consistent with the corresponding effect in adulthood (**Table S10**). This may have been due to the small available sample size of Hispanic/Latino children/adolescents.

### Transferability of Established Anthropometric Loci to Hispanic/Latino Adults

Using HISLA Stage 1 results, we assessed how many established anthropometric loci, discovered in predominantly non-Hispanic/Latino samples could be transferred to Hispanic/Latino adults, given the current sample size. As shown in **Table S11**, the index SNPs at 332 of 1280 (25.9%) previously reported BMI loci were transferable to Hispanic/Latinos. Of these BMI loci, 13 SNPs in the HISLA Stage 1 displayed genome-wide significant associations with the SNP reported in the literature (**Table S7**). **Table S12** shows that a slightly higher percentage of known height loci (1177 of 3925, or 30.0%) were transferable to Hispanic/Latinos. Forty-nine height loci displayed a genome wide significant association with height in the surrounding 1 MB interval in HISLA Stage 1 (**Table S7**), with 44 of 49 SNPs being the exact index SNP from the literature (**Table S11**). Lastly, **Tables S13-15** show that 143 of 754 (19.0%) known WHRadjBMI in both sexes combined, 103 of 504 (20.4%) in women only, and 28 of 186 (15.1%) in men only loci were transferable to Hispanic/Latinos. However, none of the index SNPs from the previous literature for WHRadjBMI reached genome-wide significance. We did observe genome-wide significant evidence for association of a SNP with WHRadjBMI in the 1 MB interval of one known region (*HOXC13*), although not replicating the exact previously reported index SNP (**Table S7**).

### Five Novel Loci and Thirty-three New Signals in Established Loci for Adult Anthropometric Traits Discovered and Replicated in a Trans-Ethnic Meta-Analytic Context

As shown in **Figure 1**, we pursued a secondary goal of assembling a trans-ethnic meta-analysis of HISLA Stage 1 with the AAAGC and GIANT consortia results to attempt to further leverage differences in allele frequencies across populations to identify additional novel loci and fine-map established loci. As anticipated, this trans-ethnic meta-analysis revealed eight new loci and 35 new signals in established loci that were associated at GWAS significance in the combined HISLA, AAAGC and GIANT meta-analysis (**Table S16, Figures S7-S52**), and independent of established SNPs within a 10Mb region (**Table 2**). Of this set, five new loci (3 BMI, 1 height, and 1 WHRadjBMI) and 33 new signals in established loci (3 BMI, 28 Height, and 2 WHRadjBMI) were validated using the adult British subsample of the UKBB. In some cases, the significance in the trans-ethnic results had additional signal driven more by the AAAGC and/or HISLA consortia, which could explain the lack of association in the UKBB British subsample (**Table S16, Figure S53**). We looked up the potentially novel findings from our trans-ethnic meta-analyses in the sample of Hispanic/Latino children/adolescents (**Table S17)**. Four trans-ethnic SNPs were associated at nominal significance in the child/adolescent sample, each having been already replicated in UKBB (**Table S16**). Three of these four loci were directionally consistent in the childhood/adolescence results with the trans-ethnic adult findings (**Table S17**). In summary, we found that two of the seven novel BMI/height trans-ethnic loci and 17 of the 33 new trans-ethnic BMI/height signals in established loci were directionally consistent between their adult directions of association and the BMI/height-for-age z-scores in children/adolescents. However, this directional consistency was not more than what would have been expected by chance alone (*P-values*_binomial_>0.10).

**Table 2.**
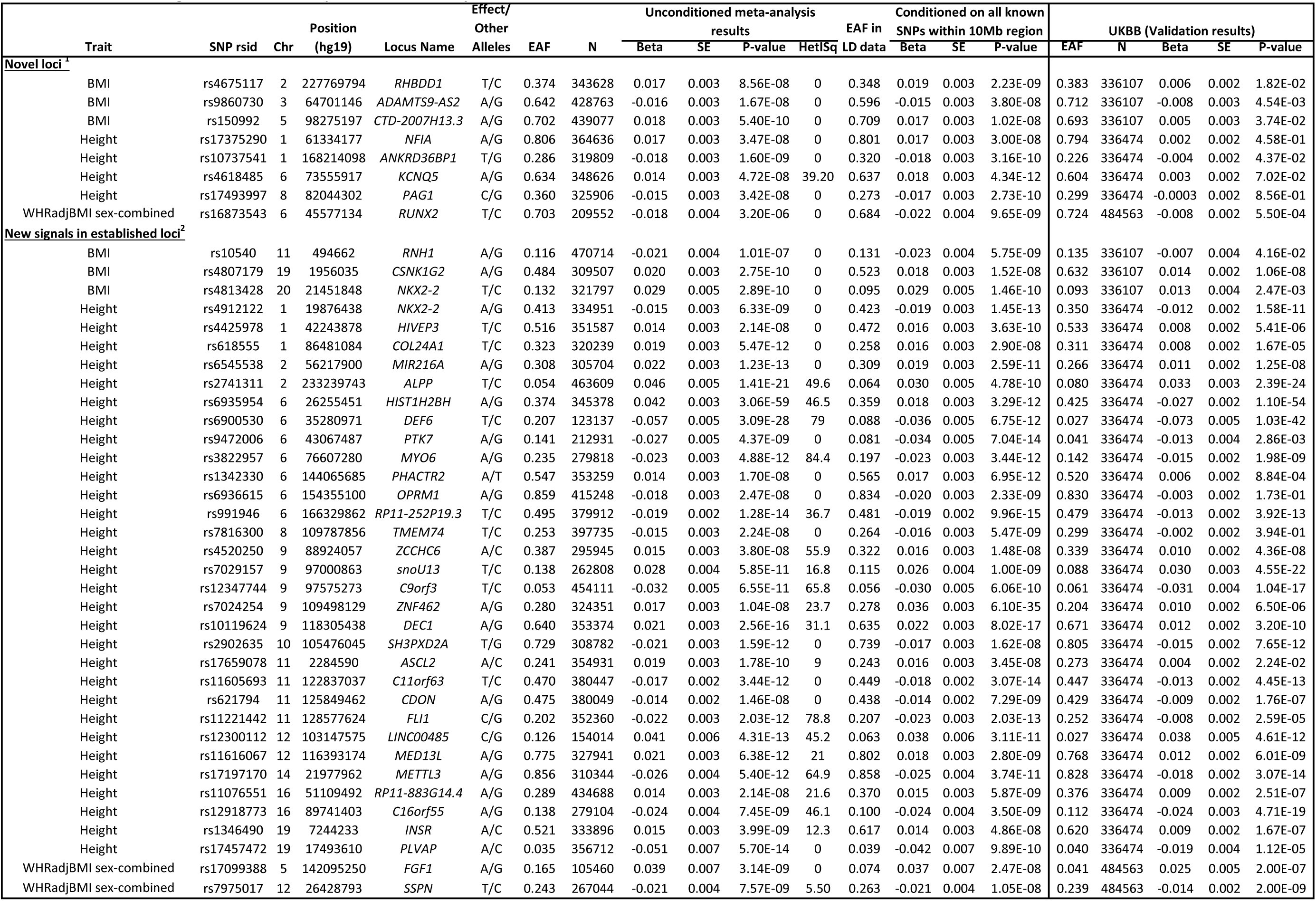
Novel loci and new signals in established loci by trait from a meta-analyses of HISLA, AAAGC, and GIANT. Abbreviations: Chr - chromosome; EAF - effect allele frequency; HetIsq - heterogeneity I- square; N - sample size; WHRadjBMI - waist to hip ratio adjusted for BMI; AAAGC- African American Anthropometry Genetics Consotrium; GIANT- Genetic Investigation of ANthropometric Traits ^1^ Each novel locus was defined by the absence of known (previously published) SNPs within 1Mb (+/-500 Kb) of the lead SNP. ^2^ Each known locus was defined by a 1Mb region around previously identified SNP(s) for the indicated trait; the known SNP(s), P<5e-8, at each established locus can be found in Table S16.

### Fine-Mapping of Trans-Ethnic Anthropometric Findings

We also fine-mapped the novel trans-ethnic findings (**Table S21**) using FINEMAP^66^ to pinpoint individual variants and genes within each locus region that have a direct effect on the trait. FINEMAP uses a shotgun stochastic search algorithm^89^ that iterates through causal configurations of SNPs by concentrating efforts on the configurations with non-negligible probability. Within a 1MB region which was a novel locus for the given trait, or included a new signal within a known locus, we report the causal configuration of SNPs with highest posterior probability and the posterior probability that each of these SNPs is causal.

For four of the five novel loci (three for BMI and one for WHRadjBMI) there was one SNP within the configuration with the highest posterior probability. For the novel height locus near *ANKRD36BP1*, there were two SNPs in the configuration with the highest posterior probability. In all five novel loci, the SNP with the highest posterior probability from each of these credible sets was either the exact SNP with the strongest GWAS evidence or in high LD (r^2^ between 0.70 and 0.99 in each ancestry) with the lead GWAS SNP. Two of these five regions had strong prioritization given high posterior probabilities (≥0.8) and small 95^th^% credible sets: 1) for BMI, the *CTD-2007H13.3* region had a posterior probability of 0.88 for rs150992 with three SNPs in the credible set, and 2) for height, the *ANKRS36BP1* region had a posterior probability of 0.93 for rs10737541 with five SNPs in the credible set. From the functional annotations (**Table S22 and S23**), we find that all three of the BMI loci, the height and WHRadjBMI loci have enhancer marks and eQTLs, most of which are in relevant tissues, e.g., adipose, muscle, thyroid, or brain.

For the other novel trans-ethnic loci, the posterior probabilities were lower, between 0.09 and 0.42; yet, four loci (rs9860730, rs17375290, rs4324883, and rs9463108) still had relatively few SNPs (<10) in the 95^th^% credible sets suggesting a narrow window (combination of variants) around the causal variant. For example, functional annotations of rs17375290 lead GWAS SNP in the NFIA locus associated with height, show it to have promoter markers in muscle, CADD score of 13.29 (CADD > 10 ranks variants among the top 10% potentially deleterious), and an eQTL with *FGGY* in Osteoclast tissue (**Table S22 and S23**). Three of the other SNPs in the credible set (rs599989, rs1762881, and rs17121184) have nominally significant (p-value 0.01 to 0.005) eQTLs with *FGGY* in osteoclast tissue but are not in high LD with rs17375290 (r^2^ range from 0.03 to 0.1). Diseases associated with *FGGY* include Lateral Sclerosis and Spastic Paraplegia 7, Autosomal Recessive, which is known to affect height.

Within the 33 novel trans-ethnic signals in known loci, 31 had configurations with more than one putative causal SNP (e.g. more than one credible set). This made sense given these are loci with multiple independent signals, as described by our earlier conditional analyses. Among the putative causal SNPs within each locus, there were a number of SNPs that represented previously-known signals (either the exact SNP or something in high LD among all ancestries). We found that for many of these the credible sets contained <10 SNPs. Among the 33 novel signals in known loci, 26 included a putative causal SNP that is the lead GWAS SNP reported here or a SNP in high LD (r^2^ > 0.75) with the lead GWAS SNP, suggesting causality for this signal in general, though perhaps maybe not initially discovered at the most- putatively-causal SNP(s). For these 24 putatively-causal SNPs, the posterior probabilities ranged from 0.09 to 1. Twenty-two of these SNPs had 95^th^% credible sets that contained <10 SNPs and 15 also had posterior probability ≥0.8.

Many have functional annotations that help support the fine-mapping results (**Table S22-S23**). For example, we find eQTLs for the three BMI signals (and enhancer marks for rs4807179) in relevant tissues including adipose, brain, muscle, and/or thyroid. The lead SNPs of these credible sets had posterior probabilities >0.75 and the credible sets included <10 SNPs. Of the 28 newly identified height signals, we find 13 putatively-causal SNPs that are the lead GWAS SNP or are in high LD (r^2^ > 0.75) with it, have <10 SNPs in the credible set and have eQTLs in relevant tissues including muscle, thyroid, adipose, lung and osteoclasts. Some also have promoter or enhancer marks in some of the same tissues. For the two WHRadjBMI signals, both have three SNPs in the most probably causal configurations. One of these causal SNPs for each region is either the lead GWAS SNP (rs7975017) or a SNP in high LD (rs17099388 and rs6895040 LD: AFR R2=1.0; AMR R2=1.0; EUR R2=1.0), has a posterior probability ≥0.95, and is the only SNP in the credible set. Furthermore, for rs7975017 we find eQTLs in thyroid for multiple genes (*BHLHE41, SSPN,* and *AC022509.3*) and enhancer marks in multiple tissues including those related to the WHRadjBMI trait, e.g., thyroid, muscle, fat, bone, and adrenal gland. Overall, across many of the novel loci and secondary signals, FINEMAP revealed SNPs with somewhat strong prioritization (posterior probability ≥0.8) and at some loci putatively-causal SNPs have small 95^th^% credible sets thus demonstrating the utility of trans-ethnic approaches to fine mapping GWAS loci.

### Trans-Ethnic Findings to Account for Population Structure in Previous GWAS

The first two PCs in the PCA (**Figure S54**) reflect geographical or population structure in Europe, corresponding to the North-South and Southeast-Southwest axes of variation, respectively. We found that the bias in effect size estimates due to stratification is most obvious for height as the phenotype is known to be differentiated across Europe.^90–92^ Effect sizes on height estimated from the GIANT and our trans-ethnic meta-analysis were both highly correlated with the loadings of the first PCA (rho = 0.125, *P-value*= 3.2×10^-94^ in GIANT; rho = 0.105, *P-value*= 3.4×10^-70^ in meta-analysis). The correlation was much lower in AAAGC and HISLA (rho = 0.012, *P-value*= 2.17×10^-4^ in AAAGC; rho = 0.007, *P-value*= 9.2×10^-2^ in HISLA; **Figure 4A**). Importantly, the magnitude of correlation was lessened in meta-analysis compared with GIANT (*P-value*= 6.6×10^-9^). Other traits were not *a priori* known to be as differentiated across Europe as height, and thus the degree of correlation between effect sizes and PC loadings are much lower in GIANT (e.g. rho = -0.025 for BMI; **Figure 4B-E**).

## DISCUSSION

Hispanic/Latinos are a unique population with continental admixture from the Americas, Africa and Europe^11–15^ and population of great interest for anthropometric studies. Here, we present results from a large-scale meta-analysis of anthropometric traits in Hispanics/Latinos. As the first of its kind, we have assembled a large sample of Hispanics/Latinos to map a total of six novel loci and 36 novel signals using both Hispanic/Latino population-specific and trans-ethnic discovery efforts (**Figure 1**). More than 1,600 anthropometric-SNP associations were transferable at nominal significance to Hispanics/Latinos—representing between 19-30% of all index sex-combined SNP-anthropometric associations (**Tables S11-13**). Sixty-seven previously reported loci reached GWAS significance at the same index or another lead SNP in our Hispanic/Latino adult sample (**Table S7**). Moreover, we established that four of seven of our novel HISLA findings were transferable to other ancestral populations at nominal significance. We note that even though these findings provide additional evidence for transferability of common loci for anthropometrics,^93^ still a number of previously-reported anthropometric loci may not be transferable to this population in part due to variability in allele frequencies or effect sizes across ancestral populations.^59^

Our conditional and fine-mapping analyses revealed 36 novel signals in established anthropometric loci, which independently replicated in HISLA Stage 2 or the UKBB British subsample. In addition, our lead SNPs for the novel BMI signals discovered at *ADCY5* (from the HISLA meta-analysis) and *ADAMTS9-AS2* (from the trans-ethnic meta-analysis) are both nominally associated with obesity status between 2-18 years of age. Three of our new trans-ethnic signals in established height loci also displayed association with height-for-age z-scores in children/adolescents between 5-18 years of age. These observations support our premise that diverse and trans-ethnic studies represent a valuable tool for identifying multiple signals and fine-mapping in established association regions. This was done with the overarching goal of identifying putative variants that will account for some of the missing heritability of complex diseases and reveal candidate genes and SNPs for functional follow-up.

In light of the notable ancestral, geographical or environmental diversity of the studies analyzed in our meta-analyses, we observed evidence of allele frequency differences for many of our novel discoveries (**Figure 3** and **Figure S53**). Similar to reports from other diverse genome-wide analyses,^59^ in some cases this allele frequency heterogeneity may drive the apparent heterogeneity effect across consortia in our HISLA, AAAGC, and GIANT meta-analysis (e.g., *IGF2BP2* I^2^=78.7; *MY06* with I^2^=84.4, **Tables 2 and S16**). These observations reinforce how studies of one predominant ancestry group, such as Europeans, may fail to identify novel loci or, more likely, new signals in known loci (given how many known loci there are currently) with allele frequency differences across ancestral populations.

**Figure 3.**
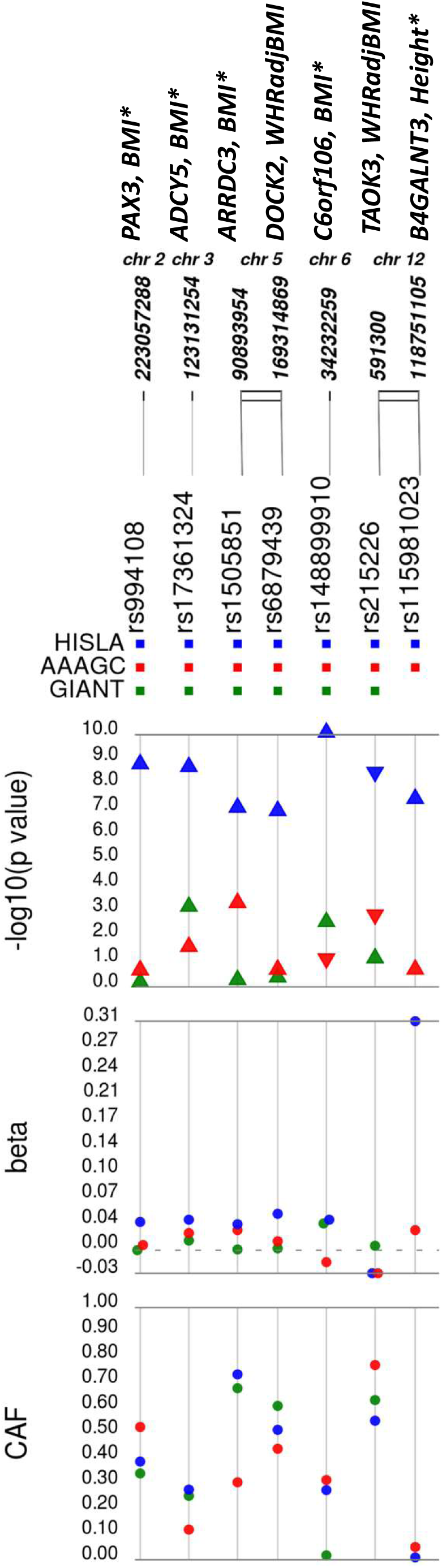
Variability in HISLA Stage 1+2, GIANT, and AAAGC P-values, Effect Sizes and Risk Allele Frequencies. Hispanic/Latino Anthropometry Consortium (HISLA); African American Anthropometry Genetics Consortium (AAAGC); Genetic Investigation of ANthropometric Traits (GIANT); WHRadjBMI - waist to hip ratio adjusted for BMI. *Asterisks indicating a SNPs that were significant either as a novel locus or new signals in a known locus.

**Figure 4.**
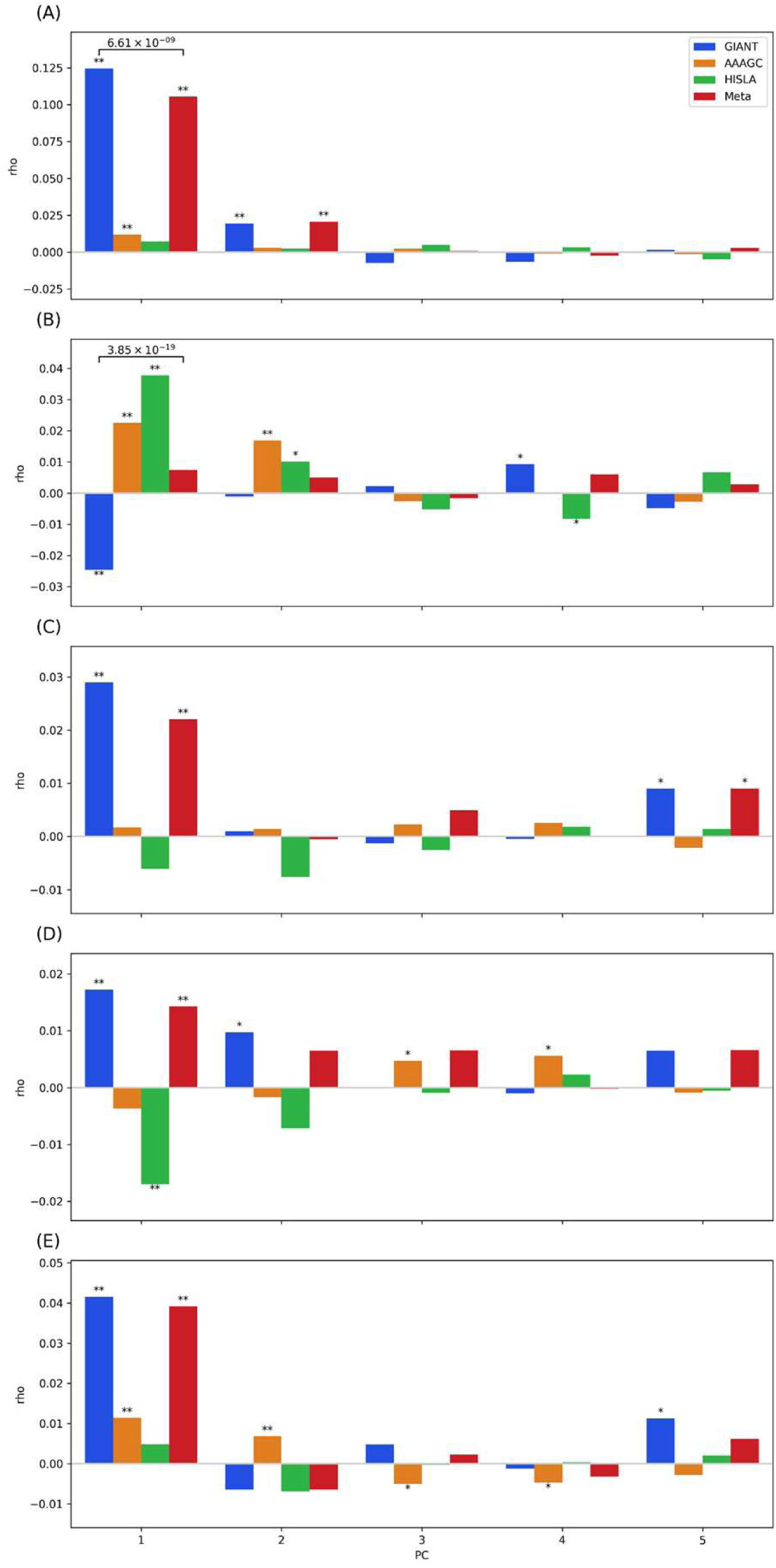
Correlations (rho) between effect estimates and the loadings of the principal components 1-5 in each consortia (HISLA, AAAGC, GIANT) and the meta-analysis of all 3 consortia (Meta) by trait. (A) height, (B) BMI, (C) Waist-to-hip ratio adjusted for BMI (WHRadjBMI) for men and women combined, (D) WHRadjBMI for women only, (E) WHRadjBMI for men only. Hispanic/Latino Anthropometry Consortium (HISLA); African American Anthropometry Genetics Consortium (AAAGC); Genetic Investigation of ANthropometric Traits (GIANT); WHRadjBMI - waist to hip ratio adjusted for BMI

Residual uncorrected stratification in GWAS could result in biased estimates of effect sizes.^39^ For example, effect sizes on height from GIANT were reported to be significantly correlated with North-South axis of variation in Europe suggesting residual uncorrected stratification,^92; 94; 95^ which we also observe here. Note that the residual stratification effect is subtle, and while the effect sizes may be biased, this does not imply the identified associations are spurious. For example, compared with effect sizes on height from UKBB, which is based on a single homogeneous population and results in better control of population stratification, the genetic correlation between GIANT and UKBB was 0.94.^92^

Of the three traits studied here, height is the most stratified in Europe. The correlation coefficient between effect sizes on height and PC loadings reached 0.125 in the GIANT only for PC1, while it was much smaller for other traits (e.g., the maximum |*rho*| = 0.042 in GIANT on WHR using only males on PC1). The decrease in bias in trans-ethnic meta-analysis was also obvious in height. The correlation with PC1 was non-significant in HISLA (*rho* = 0.007) and statistically significant but weak in AAAGC (*rho =* 0.012), consistent with a decreased impact of European population stratification on the estimate of effect size in AAAGC and HISLA. This decreased correlation could be due to large non-European ancestries known in these populations (Africans and Native Americans, respectively) that are less affected by population stratification in Europe; it could also be that by using European ancestry based loadings we are less likely to detect non-European based population stratification patterns or that smaller sample sizes in these cohorts resulting in greater noise in effect size estimates. Regardless of the reason, compared to GIANT alone, trans-ethnic meta-analysis of the three cohorts showed less impact of uncorrected stratification in effect size estimates, even though the sample size in AAAGC and HISLA are comparably small. For other traits, the conclusions are qualitatively similar: that trans-ethnic meta-analysis lessened the bias due to stratification, even though the bias in GIANT was not as strong in the first place.

As described above, in this study we were able to 1) discover six novel loci with a notably smaller analytic size than other anthropometric consortia like GIANT, 2) describe 36 new signals in established loci in HISLA or our trans-ethnic meta-analysis, and 3) generate trans-ethnic effect estimates with better control for population structure. Taken together, these findings indicate the added value of building large, more diverse GWAS in the near future.

Gene expression and bioinformatic analyses of our population-specific (**Table S18-S20**) and trans-ethnic findings in newly discovered loci gave us important insights into the underlying biology of obesity, bone development and growth (**Tables S22-S23**). For example, the previously reported BMI locus *C6orf106* has also been associated with adult height^96; 97^ and height change during puberty.^98^ The first BMI signal described at *C6orf106* was at index SNP, rs205262, an eQTL for another gene within the region, *SNRPC,* in European ancestry samples.^38^ A second signal (rs75398113) has also been reported at *SNRPC* for extremes of the body mass index distribution.^99^ Yet, our novel signal led by rs148899910 is more than 300kb away and in low LD with these two index SNPs (r^2^=0.01-0.05 in AMR). More recently, rs148899910 has been associated with height in Korean women.^100^ Using whole blood gene expression data from 606 participants of the Cameron County Hispanic Cohort, we find evidence that our novel BMI signal at rs148899910 is an eQTL for increased gene expression of *C6orf1* (p-value=3×10^-7^) and not any other genes in the region (**Table S18**).

In general, the lead SNPs from our HISLA only meta-analyses appear relatively benign (not pathogenic) based on CADD and FATHMM-XF scores (**Table S20**). All SNPs potentially change motifs. Both rs17361324 (*ADCY5*) and rs215226 (*B4GALNT3*) have enhancer and promoter histone marks and eQTLs in the respective genes in relevant tissues. For BMI, there is an eQTL for rs17361324-ADCY5 in thyroid, and *ADCY5* has been previously associated with type 2 diabetes,^101^ BMI,^102^ central obesity traits,^43^ height,^51^ birth outcomes,^103–105^ and a number of other phenotypes. Additionally, rs17361324 is proximate to an *ADCY5* intronic variant (rs1093467, r²=0.3 in 1000 Genomes AMR) that is highly conserved across species (Haploreg v4.1). For height, there is an eQTL for rs215226-B4GALNT3 in aortic, and coronary arteries, and tibial nerve. The lead SNP for the height signal in *B4GALNT3*, rs215226, has enhancer histone marks in bone and muscle, and promoter marks in muscle tissue. In addition, the variant rs215226 (*B4GALNT3*) has a posterior probability of 1 for causality in FINEMAP analyses (see **Table S9**). Other interesting information about these regions is provided in **Table S19.**

The lead SNPs at our newly discovered trans-ethnic loci were mainly located in intronic and intergenic regions (**Table S22**) and were benign. One exception was the novel locus *C11orf63* associated with height led by rs11605693, which showed pathogenic scores for CADD and FATHMM-XF (CADD score=17.1 and FATHMM-XF score=0.87). This lead SNP has an eQTL in *C11orf63* for adipose, tibial nerve, and testis. *C11orf63*, junctional cadherin complex regulator, is responsible for ependymal cells that line the brain and spinal cord.

Among the trans-ethnic findings, a new signal at a known locus for BMI, rs10540 at *RNH1*, has a posterior probability of 0.82 as one of two causal variants in the locus, and is an eQTL for a wide range of tissues and genes (see **Tables S21 and S23**), potentially making it relevant to body mass. A new signal in a known locus for height, led by rs12918773 that has a posterior probability of 0.98 and is one of four casual variants suggested from fine-mapping in the locus (**Table S21**), has an eQTL (in lung, thyroid, tibial nerve and artery, breast, testis) with *CDK10*, a gene also associated with growth retardation.^106^ In addition, rs1342330, another new signal in a known height locus, has a low regulomeDB score at 2b and several enhancer and promoter histone marks in relevant tissues (Tables S22). As an intronic variant, it is an eQTL in the pancreas with *PHACTR2* (**Table S23**), a gene associated with body dysmorphic disorder.^107^ While many of the novel loci/signals appeared to be benign based on CADD and FATHMM-XF scores, they still show enhancer and promoter histone marks in trait relevant tissues such as adipose, bone, and muscle, thymus, brain, and adrenal gland.

Large-scale analyses of diverse populations hold great potential for advancing the field of genetic epidemiology.^59^ This study illustrates how studying admixed populations, like Hispanic/Latinos, and leveraging them in trans-ethnic epidemiologic investigations, can yield additional insights into the genetic architecture of anthropometric traits. Future discovery efforts in Hispanic/Latino populations and with other diverse populations will address the research gap between who is studied and who is affected by conditions like obesity, to the benefit of both public health and precision medicine.

## Supplemental Data

Supplemental Data include 23 tables and 54 figures.

## Acknowledgements

The Baependi Heart Study was supported through a collaborative effort by FAPESP and Brazil Health Ministry (PROADI). ACP was supported by NHLBI R01HL141881-01A1. The Hispanic Community Health Study/Study of Latinos was carried out as a collaborative study supported by contracts from the National Heart, Lung, and Blood Institute (NHLBI) to the University of North Carolina (N01-HC65233), University of Miami (N01-HC65234), Albert Einstein College of Medicine (N01-HC65235), Northwestern University (N01-HC65236), and San Diego State University (N01-HC65237). The following Institutes/Centers/Offices contribute to the HCHS/SOL through a transfer of funds to the NHLBI: National Institute on Minority Health and Health Disparities, National Institute on Deafness and Other Communication Disorders, National Institute of Dental and Craniofacial Research, National Institute of Diabetes and Digestive and Kidney Diseases, National Institute of Neurological Disorders and Stroke, NIH Institution-Office of Dietary Supplements. The Genetic Analysis Center including (SMS, CCL, AMS) at the University of Washington was supported by NHLBI and NIDCR contracts (HHSN268201300005C AM03 and MOD03). MESA and the MESA SHARe projects are conducted and supported by the National Heart, Lung, and Blood Institute (NHLBI) in collaboration with MESA investigators. Support for MESA is provided by contracts 75N92020D00001, HHSN268201500003I, N01-HC-95159, 75N92020D00005, N01-HC-95160, 75N92020D00002, N01-HC-95161, 75N92020D00003, N01-HC-95162, 75N92020D00006, N01-HC-95163, 75N92020D00004, N01-HC-95164, 75N92020D00007, N01-HC-95165, N01- HC-95166, N01-HC-95167, N01-HC-95168, N01-HC-95169, UL1-TR-000040, UL1-TR-001079, UL1-TR-001420, UL1-TR-001881, and DK063491. Funding for SHARe genotyping was provided by NHLBI Contract N02-HL-64278. Genotyping was performed at Affymetrix (Santa Clara, California, USA) and the Broad Institute of Harvard and MIT (Boston, Massachusetts, USA) using the Affymetrix Genome-Wide Human SNP Array 6.0.

The FIND study was supported by grants U01DK57292, U01DK57329, U01DK057300, U01DK057298, U01DK057249, U01DK57295, U01DK070657, U01DK057303, and U01DK57304 from the National Institute of Diabetes and Digestive and Kidney Diseases (NIDDK) and, in part, by the Intramural Research Program of the NIDDK. Support was also received from the National Heart, Lung and Blood Institute grants U01HL065520, U01HL041654, and U01HL041652. This project has been funded in whole or in part with federal funds from the National Cancer Institute, National Institutes of Health (NIH), under contract N01- CO-12400 and the Intramural Research Program of the NIH, National Cancer Institute, Center for Cancer Research. This work was also supported by the National Center for Research Resources for the General Clinical Research Center grants: Case Western Reserve University, M01-RR-000080; Wake Forest University, M01-RR-07122; Harbor-University of California, Los Angeles Medical Center, M01-RR-00425; College of Medicine, University of California, Irvine, M01-RR-00827–29; University of New Mexico, HSC M01-RR-00997; and Frederic C. Bartter, M01-RR-01346. Computing resources were provided, in part, by the Wake Forest School of Medicine Center for Public Health Genomics. The funders had no role in study design, data collection and analysis, decision to publish, or preparation of the manuscript.

The Northern California Breast Cancer Family Registry (NC-BCFR) is supported by grant UM1 CA164920 from the U.S. National Cancer Institute. The content of this manuscript does not necessarily reflect the views or policies of the National Cancer Institute or any of the collaborating centers in the Breast Cancer Family Registry (BCFR), nor does mention of trade names, commercial products, or organizations imply endorsement by the United States Government or the BCFR. The San Francisco Bay Area Breast Cancer Study (SFBCS) was supported by grants R01 CA63446 and R01 CA77305 from the National Cancer Institute, grant DAMD17-96-1-6071 from the U.S. Department of Defense, and grant 7PB-0068 from the California Breast Cancer Research Program.

Several studies were also supported by the National Institutes of Health, National Heart, Lung, and Blood Institute in collaboration with the Mexican-American Coronary Artery Disease Project (MACAD) HL-088457, the HTN-IR study HL-0697974, the Genetics of Latinos Diabetic Retinopathy (GOLDR) study grant EY14684, Hypertriglyceridemia (HyperTG) study contract R01 HL0767711, and DK-079888 (work related to insulin clearance in HTN-IR, MACAD, and NIDDM-Atherosclerosis Study (NIDDM-Athero) funded by the NHLBI. The Mexico City 1, Mexico City 2 and Mexico Kids Case control studies were supported in Mexico by the Fondo Sectorial de Investigación en Salud y Seguridad Social (SSA/IMSS/ISSSTECONACYT, project 150352), Temas Prioritarios de Salud Instituto Mexicano del Seguro Social (2014-FIS/IMSS/PROT/PRIO/14/34), and the Fundación IMSS. We thank Miguel Alexander Vazquez Moreno, Daniel Locia and Araceli Méndez Padrón for technical support in Mexico. In Canada, this research was enabled in part by two CIHR Operating grants to EJP, a CIHR New Investigator Award to EJP and by support provided by Compute Ontario (www.computeontario.ca), and Compute Canada (www.compute.canada.ca). The SAMAFS was supported by HL045522, DK053889, DK047482, and MH059490. The Santiago Longitudinal Study (SLS) was supported by the Eunice Kennedy Shriver National Institute of Child Health & Human Development (R01 HD033487-15), National Institute on Drug Abuse (R01 DA021181- 05), and National Heart Lung, and Blood Institute (T32 HL079891-11). The Viva La Familia Study was supported by R01DK59264 and R01DK080457.

AGL was supported by T32 HD091058, P2C HD050924, and P30 AG066615. AGT was supported by T32HL007055. ARL is supported by the Leverhulme Trust (F/07 134/DF), BBSRC (BB/I021213/1), the Excellence Initiative of Aix-Marseille University - A*MIDEX (a French “Investissements d’Avenir” programme), the National Natural Science Foundation of China (#31771393), Universidad de Antioquia (CODI sostenibilidad de grupos 2013- 2014 and MASO 2013-2014). The other PIs of the Consortium for the Analysis of the Diversity and Evolution of Latin America (CANDELA) were supported by multiple grants (VA-A: 0051 1 3190000, GB: 0054 (0)280 488-3184, M-CB: 0051 1 3190000, SC-Q: 0056 58 2205073). The investigators of BioME 1 and 2 were supported by the following grants (MG-M: 0052155282325, MP: 0057 320 7034343, CS: 0055 51 999523134, RAJS: 0052153501900). KA was supported by 0044 (0)20 3108 4003. LF was supported by R01CA204797. LF-R was supported by an American Heart Association (AHA) predoctoral grant (13PRE16100015). MG, KLY and KEN were supported by AHA grant 13GRNT16490017 and 15GRNT25880008, R01DK089256 and R01DK101855. In addition, LF-R, CAH, and RFJL were also supported by R01DK101855. XRG was supported by NIH grant R01EY022651. KEN was also supported by R01HD057194, R01DK122503, R01HG010297, R01HL142302, R01HL143885, and R01HG009974. LG was supported by T32 HL129982 and AGT was supported by T32HL007055. QQ was supported by R01HL060712, R01HL140976, and R01DK119268. SFAG was supported by the Daniel B. Burke Endowed Chair for Diabetes Research and R01 HD056465. In addition the above MESA and MESA SHARe funding, XG, MAA, Y-DIC, JY, and JIR were supported by EY14684, HL-0767711, HL- 0697974, HL-088457, NEI EY11753, UL1-TR-001881. XL and KR were supported by R01 HL0767711 and DK-079888. ZA, TAB, JT, and AHX were supported by HTN-IR funding (HL- 0697974); PMG, YH, EI, and KDT were supported by GOLDR funding (EY-14684); MOG, WH, KL, and KS were supported by MACAD funding (HL-088457). AEJ was supported in part by K99/R00 HL130580. EMJ was supported by R01 CA063446, R01 CA077305, DOD RP9590546, CBCRP 7PB-0068. JM was supported by American Diabetes Association Innovative and Clinical Translational Award 1-19-ICTS-068 and by the National Human Genome Research Institute (U01HG011723). Computation for part of this work was supported by the University of Southern California’s Center for Advanced Research Computing (https://carc.usc.edu).

