## Supplemental Figures 1-54 for "Ancestral diversity improves discovery and fine-mapping of genetic loci for anthropometric traits - the Hispanic/Latino Anthropometry Consortium"

### Slide 1
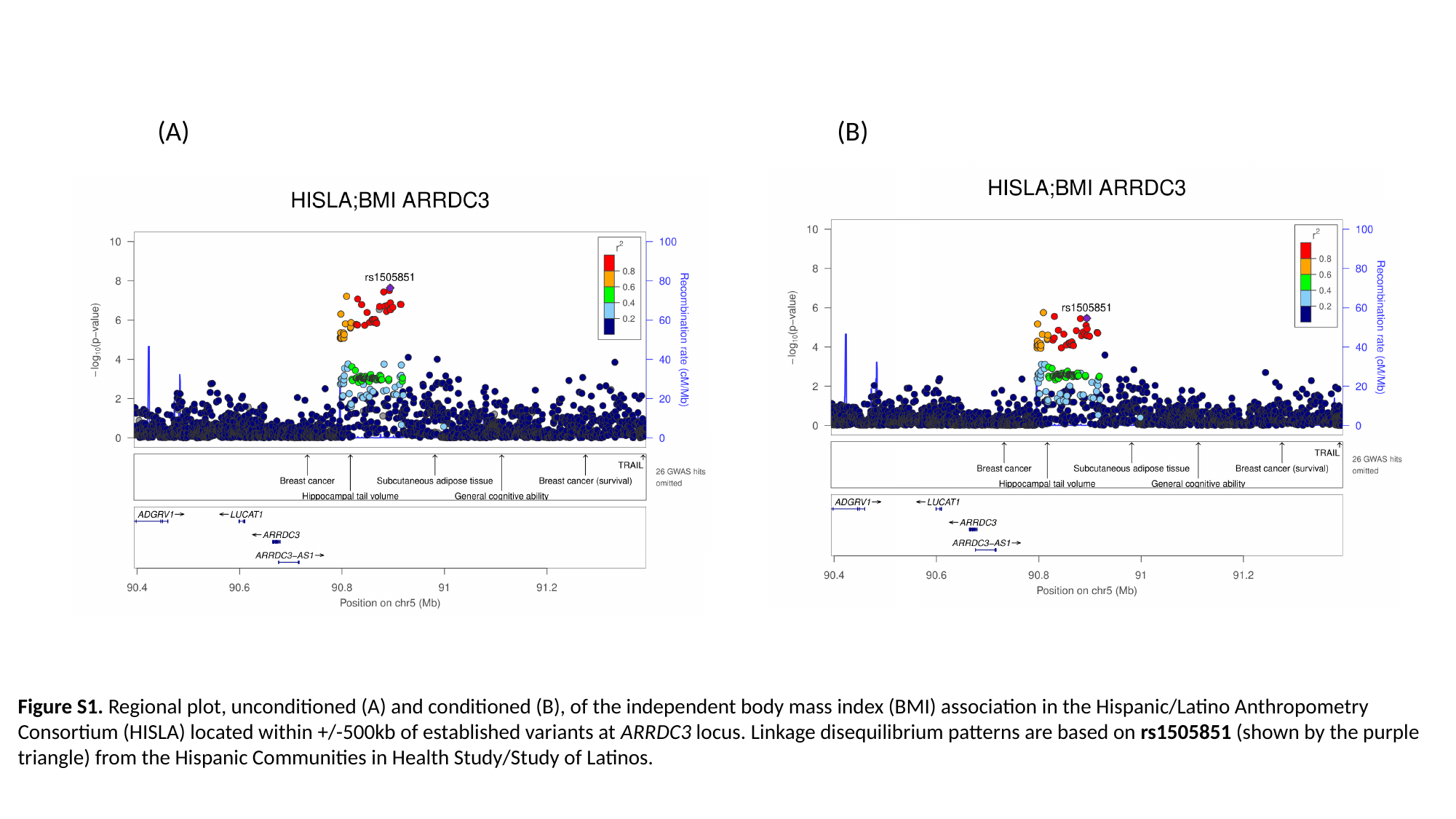

(A) (B)
Figure S1. Regional plot, unconditioned (A) and conditioned (B), of the independent body mass index (BMI) association in the Hispanic/Latino Anthropometry Consortium (HISLA) located within +/-500kb of established variants at ARRDC3 locus. Linkage disequilibrium patterns are based on rs1505851 (shown by the purple triangle) from the Hispanic Communities in Health Study/Study of Latinos.

### Slide 2
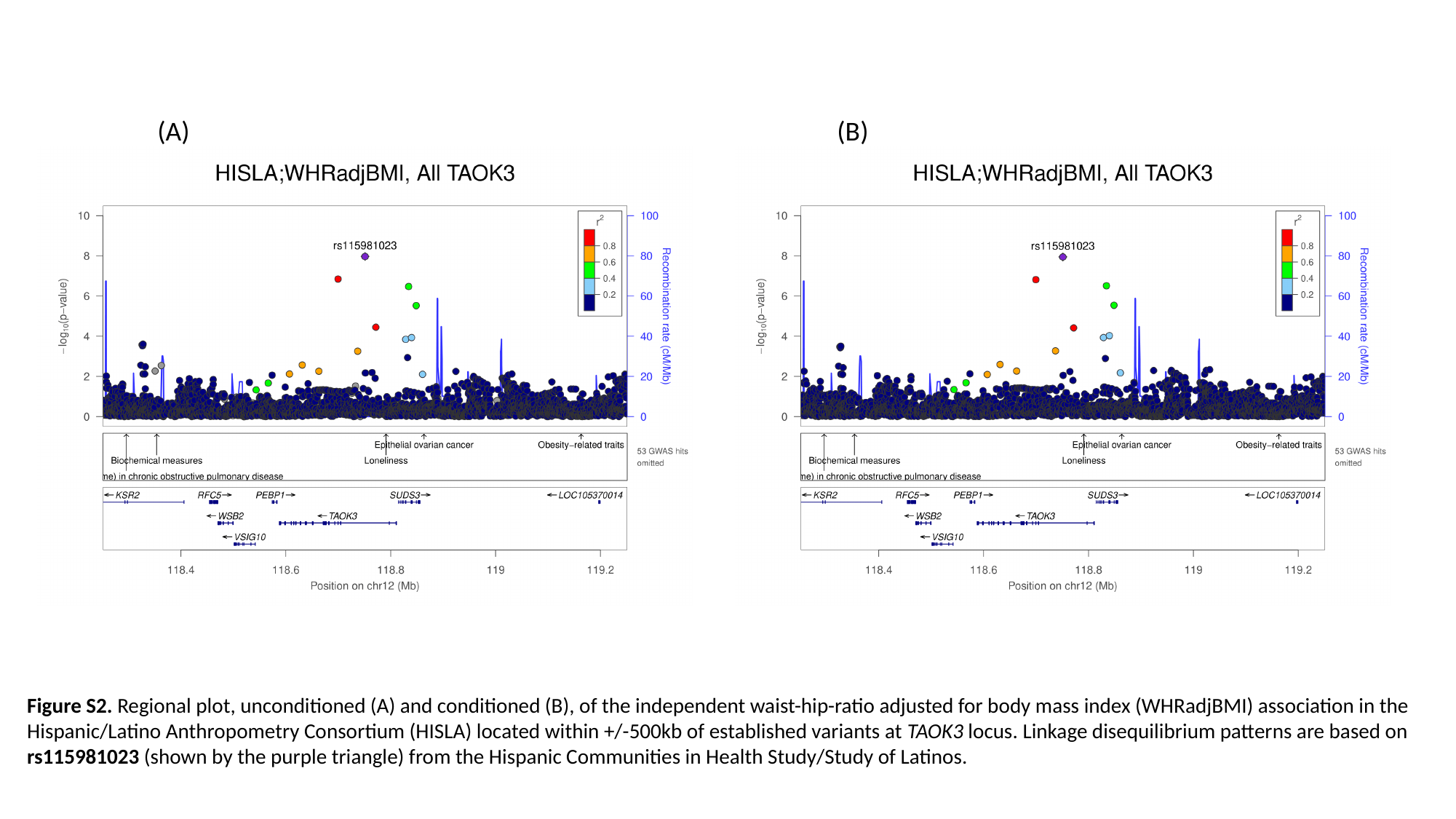

(A) (B)
Figure S2. Regional plot, unconditioned (A) and conditioned (B), of the independent waist-hip-ratio adjusted for body mass index (WHRadjBMI) association in the Hispanic/Latino Anthropometry Consortium (HISLA) located within +/-500kb of established variants at TAOK3 locus. Linkage disequilibrium patterns are based on rs115981023 (shown by the purple triangle) from the Hispanic Communities in Health Study/Study of Latinos.

### Slide 3
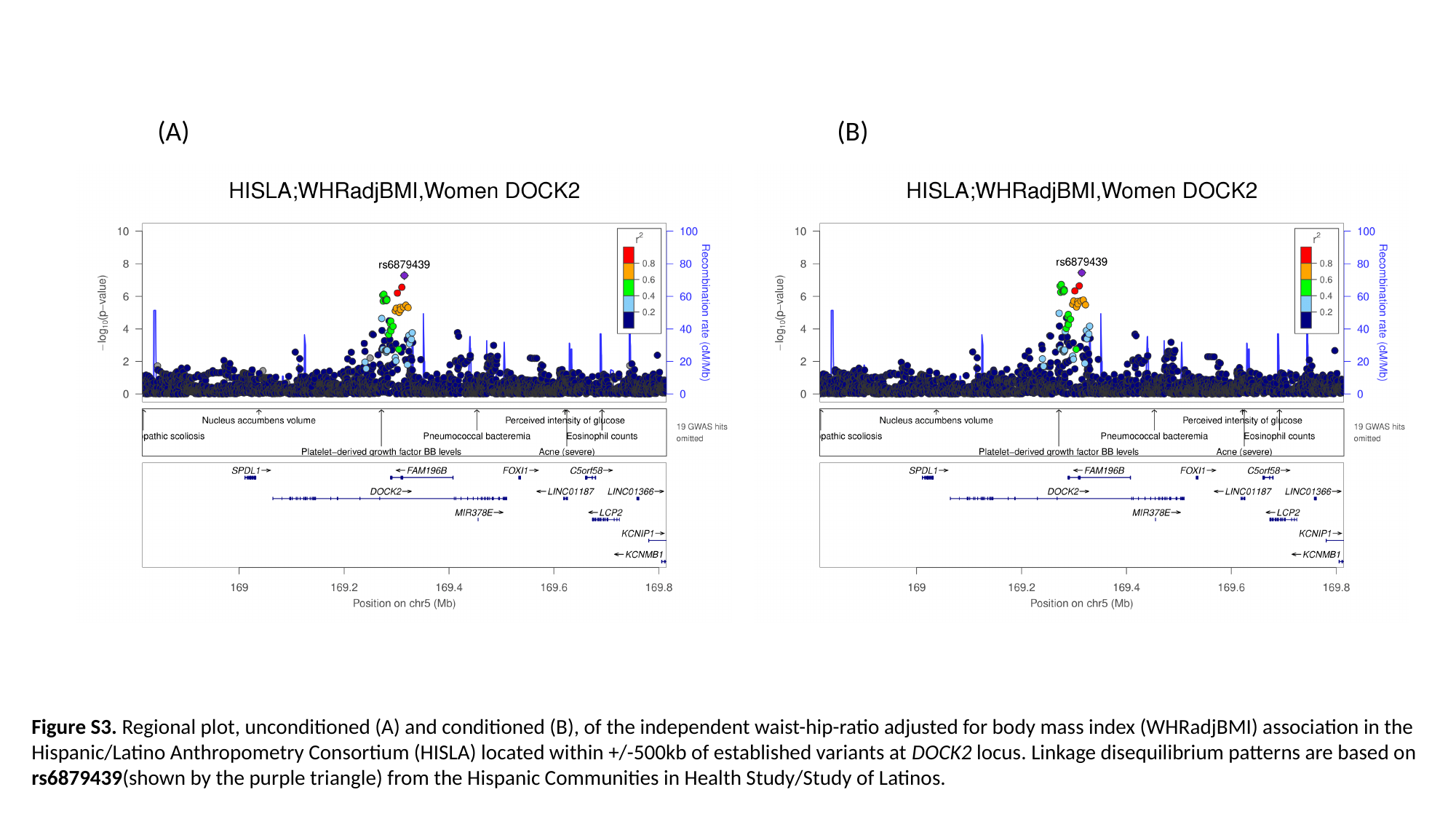

(A) (B)
Figure S3. Regional plot, unconditioned (A) and conditioned (B), of the independent waist-hip-ratio adjusted for body mass index (WHRadjBMI) association in the Hispanic/Latino Anthropometry Consortium (HISLA) located within +/-500kb of established variants at DOCK2 locus. Linkage disequilibrium patterns are based on rs6879439(shown by the purple triangle) from the Hispanic Communities in Health Study/Study of Latinos.

### Slide 4
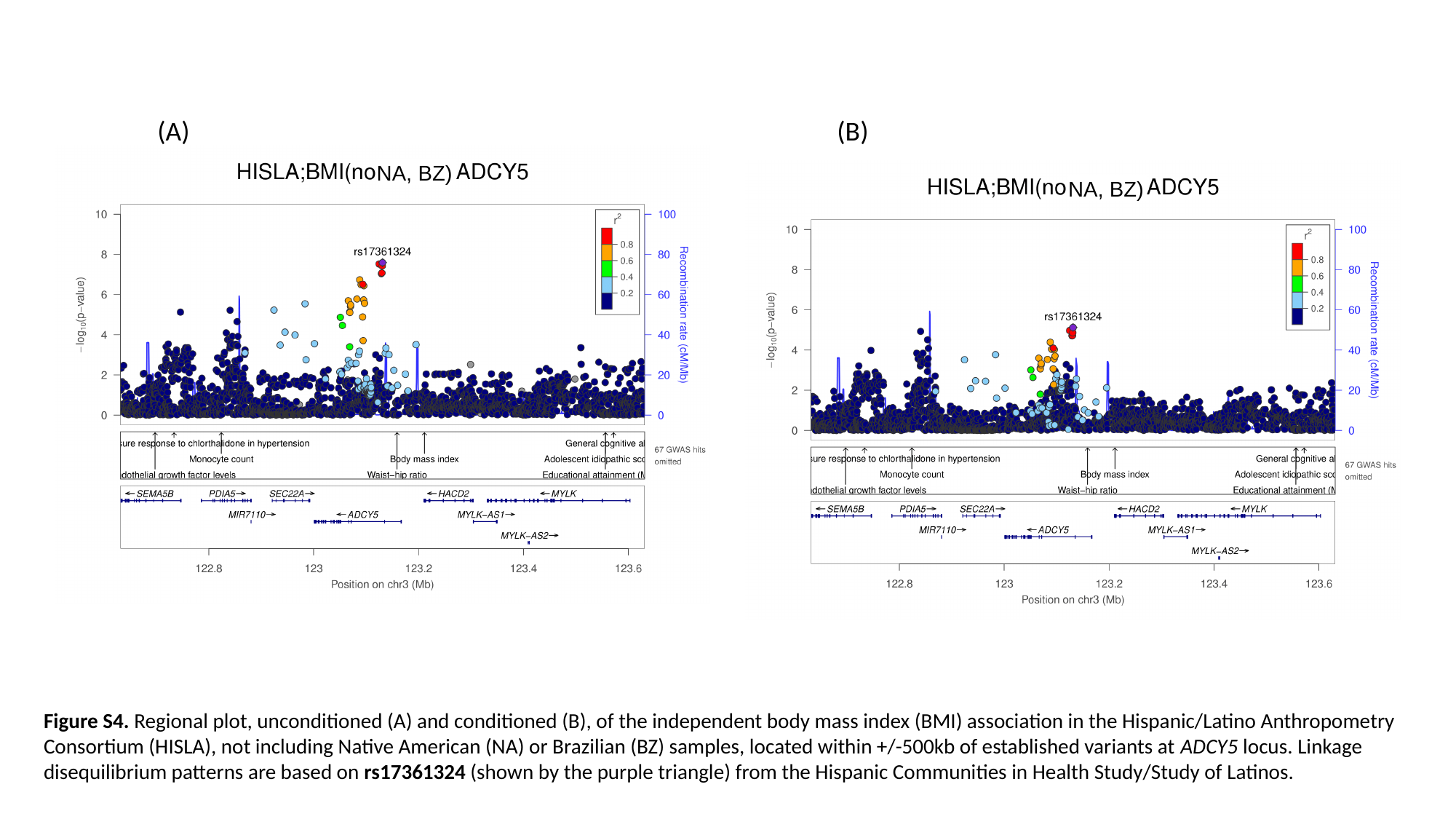

(A) (B)
NA, BZ)
NA, BZ)
Figure S4. Regional plot, unconditioned (A) and conditioned (B), of the independent body mass index (BMI) association in the Hispanic/Latino Anthropometry Consortium (HISLA), not including Native American (NA) or Brazilian (BZ) samples, located within +/-500kb of established variants at ADCY5 locus. Linkage disequilibrium patterns are based on rs17361324 (shown by the purple triangle) from the Hispanic Communities in Health Study/Study of Latinos.

### Slide 5
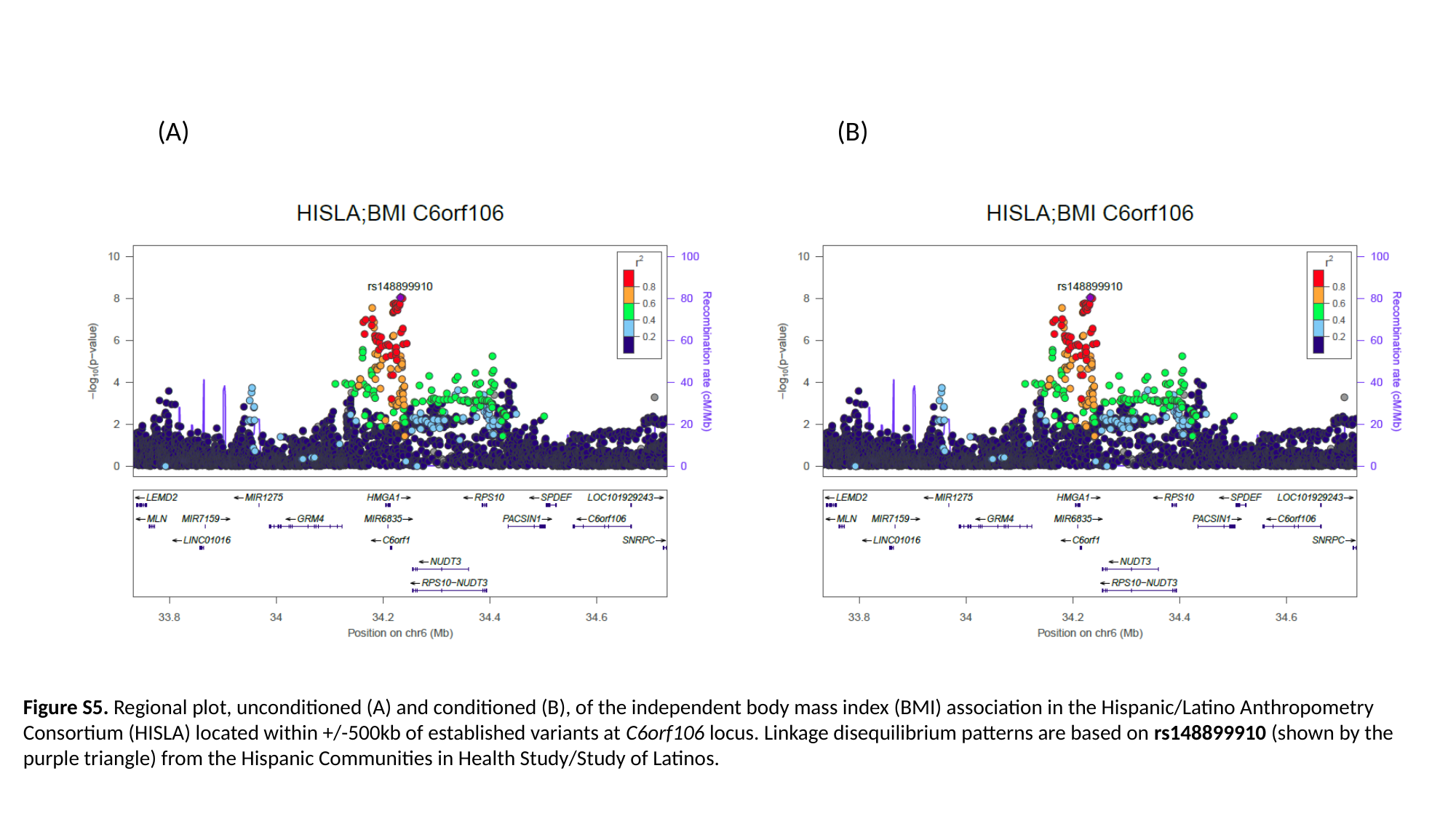

(A) (B)
Figure S5. Regional plot, unconditioned (A) and conditioned (B), of the independent body mass index (BMI) association in the Hispanic/Latino Anthropometry Consortium (HISLA) located within +/-500kb of established variants at C6orf106 locus. Linkage disequilibrium patterns are based on rs148899910 (shown by the purple triangle) from the Hispanic Communities in Health Study/Study of Latinos.

### Slide 6
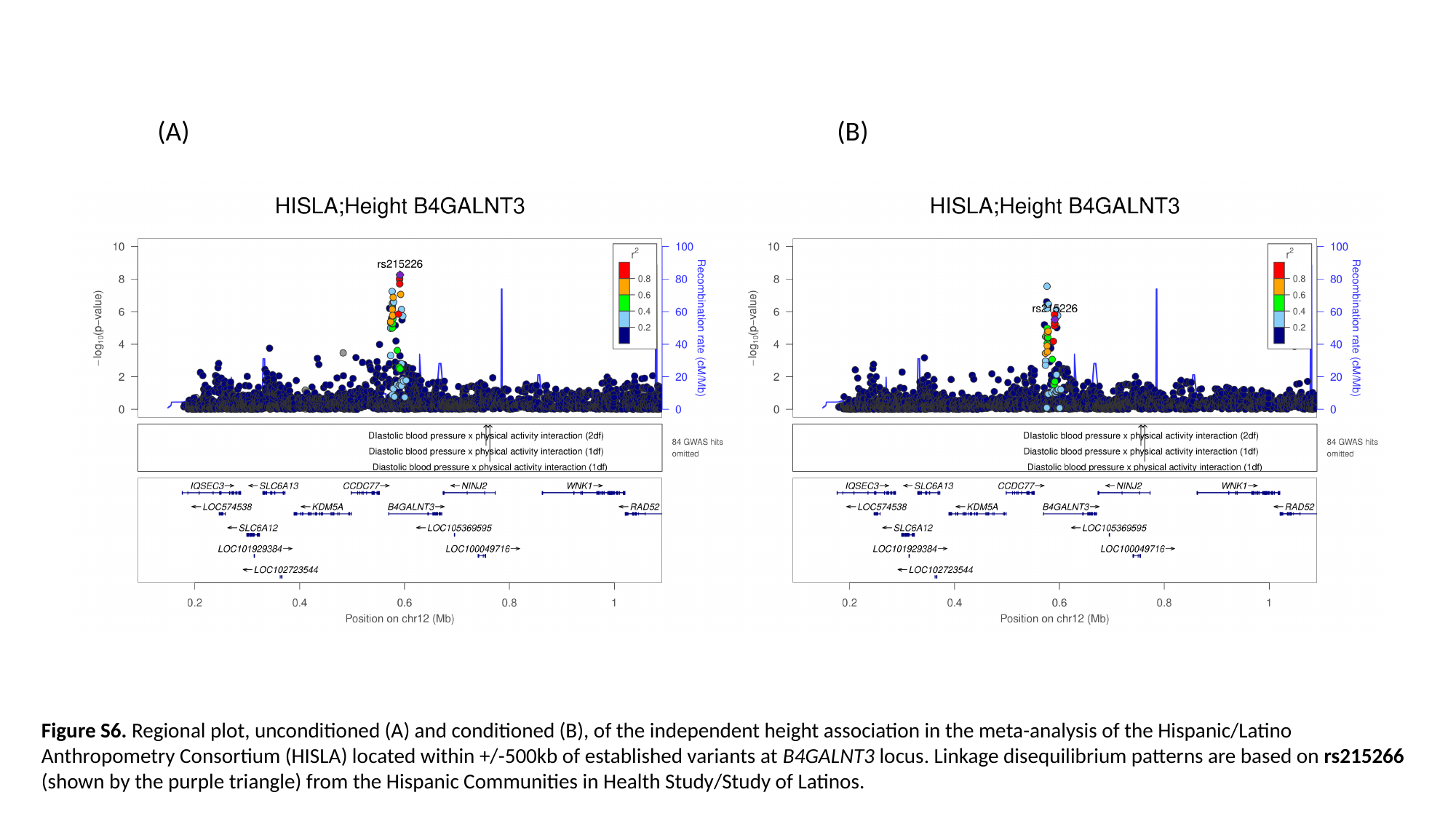

(A) (B)
Figure S6. Regional plot, unconditioned (A) and conditioned (B), of the independent height association in the meta-analysis of the Hispanic/Latino Anthropometry Consortium (HISLA) located within +/-500kb of established variants at B4GALNT3 locus. Linkage disequilibrium patterns are based on rs215266 (shown by the purple triangle) from the Hispanic Communities in Health Study/Study of Latinos.

### Slide 7
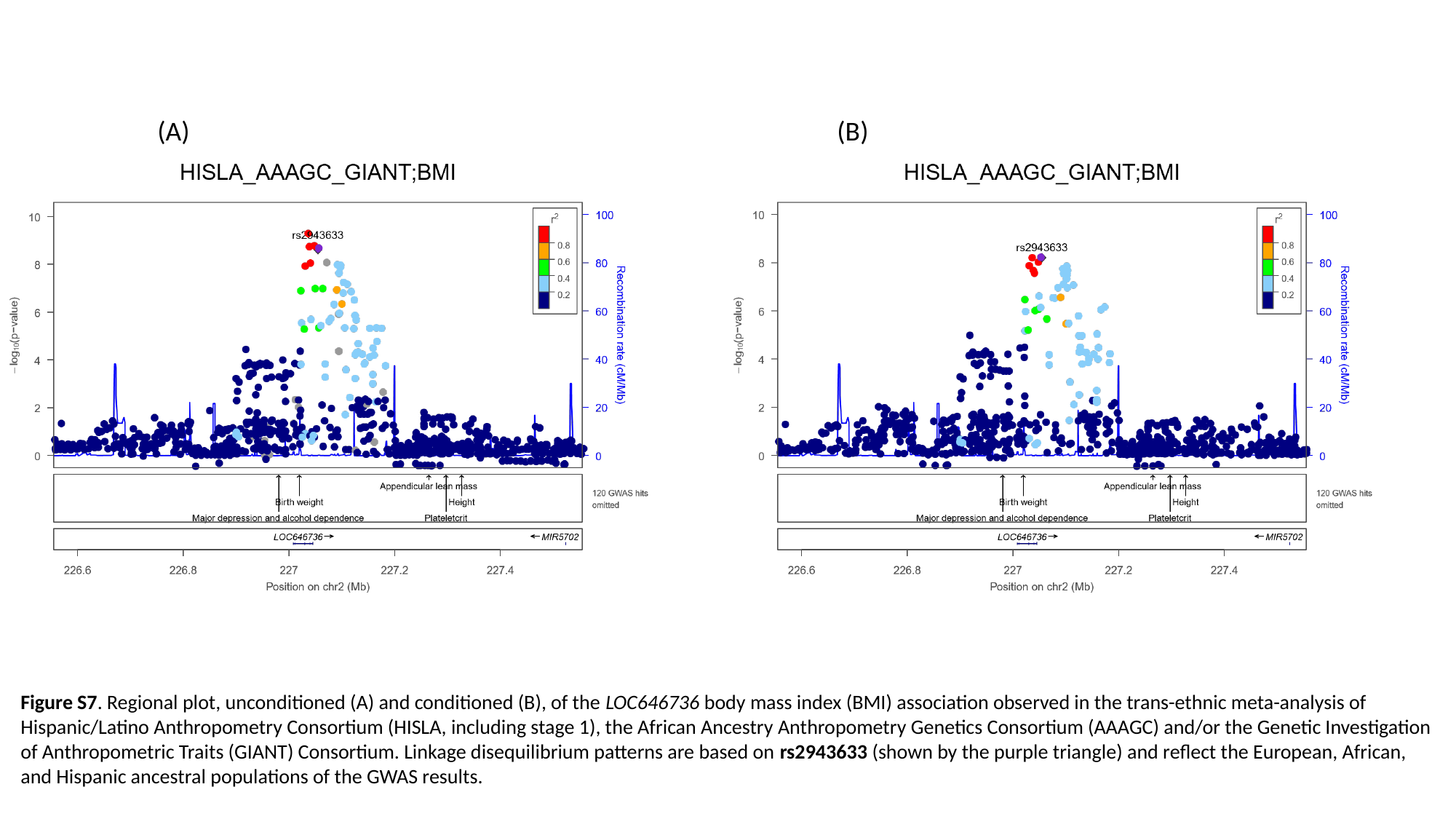

(A) (B)
Figure S7. Regional plot, unconditioned (A) and conditioned (B), of the LOC646736 body mass index (BMI) association observed in the trans-ethnic meta-analysis of Hispanic/Latino Anthropometry Consortium (HISLA, including stage 1), the African Ancestry Anthropometry Genetics Consortium (AAAGC) and/or the Genetic Investigation of Anthropometric Traits (GIANT) Consortium. Linkage disequilibrium patterns are based on rs2943633 (shown by the purple triangle) and reflect the European, African, and Hispanic ancestral populations of the GWAS results.

### Slide 8
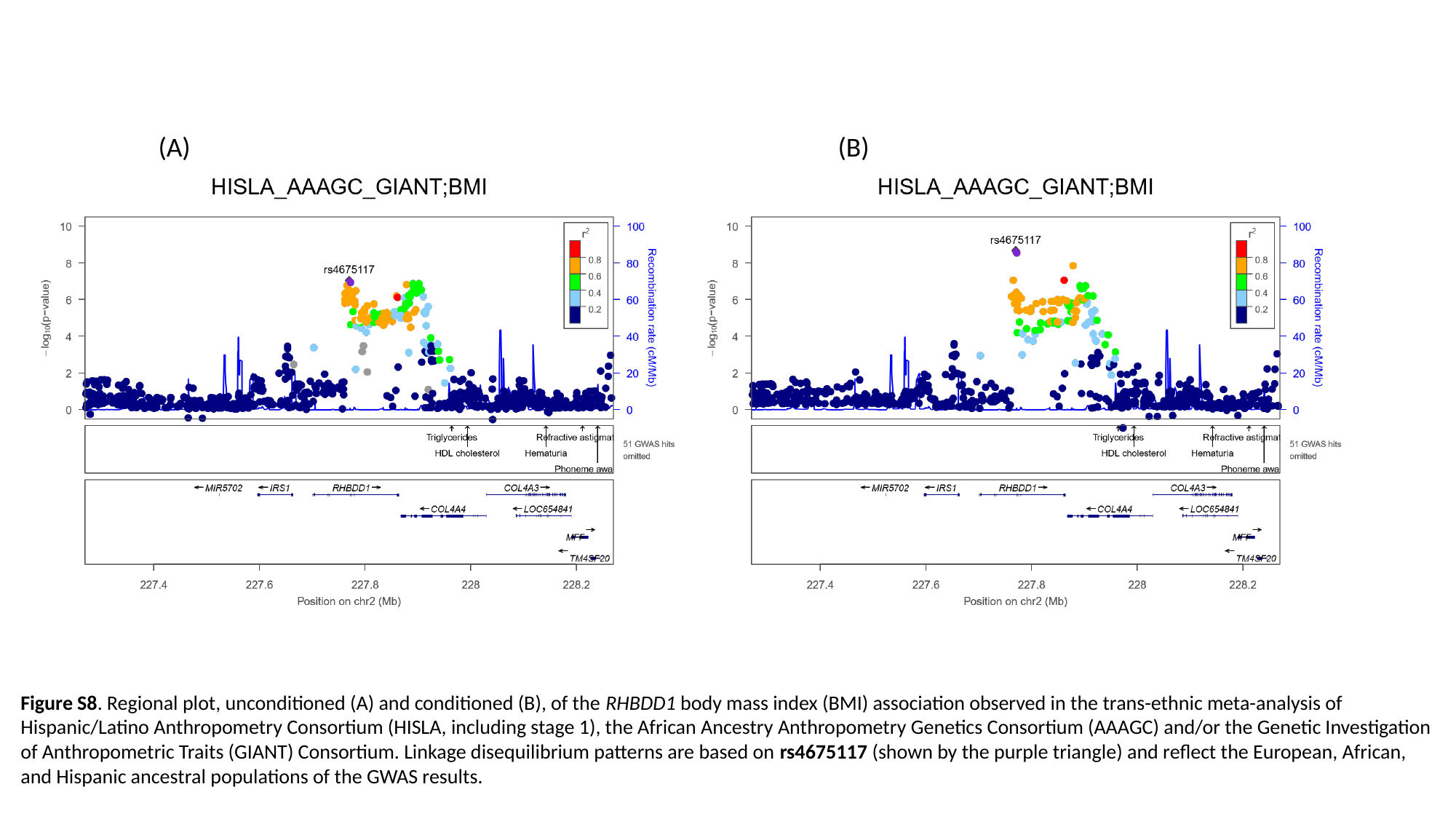

(A) (B)
Figure S8. Regional plot, unconditioned (A) and conditioned (B), of the RHBDD1 body mass index (BMI) association observed in the trans-ethnic meta-analysis of Hispanic/Latino Anthropometry Consortium (HISLA, including stage 1), the African Ancestry Anthropometry Genetics Consortium (AAAGC) and/or the Genetic Investigation of Anthropometric Traits (GIANT) Consortium. Linkage disequilibrium patterns are based on rs4675117 (shown by the purple triangle) and reflect the European, African, and Hispanic ancestral populations of the GWAS results.

### Slide 9
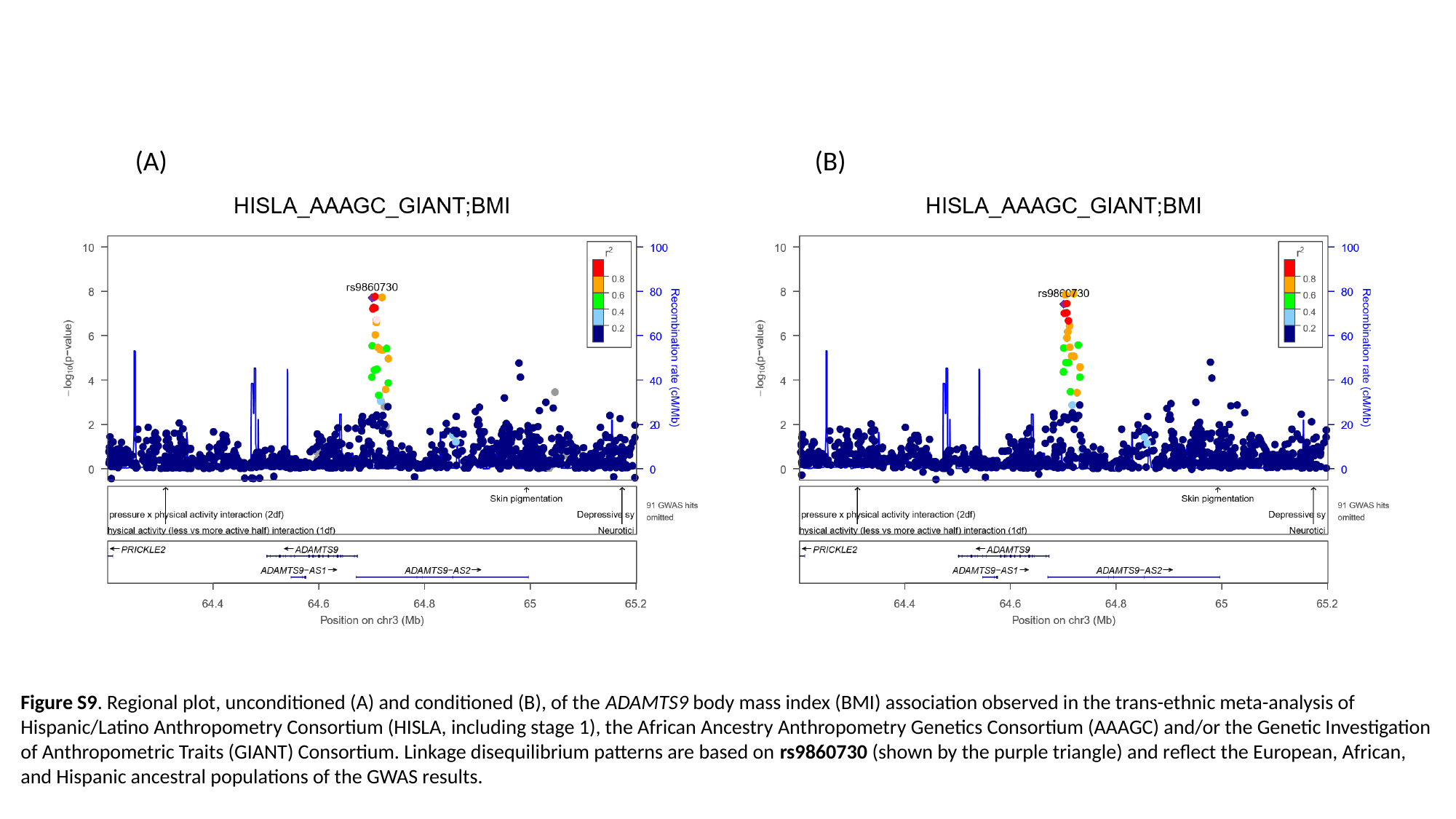

(A) (B)
Figure S9. Regional plot, unconditioned (A) and conditioned (B), of the ADAMTS9 body mass index (BMI) association observed in the trans-ethnic meta-analysis of Hispanic/Latino Anthropometry Consortium (HISLA, including stage 1), the African Ancestry Anthropometry Genetics Consortium (AAAGC) and/or the Genetic Investigation of Anthropometric Traits (GIANT) Consortium. Linkage disequilibrium patterns are based on rs9860730 (shown by the purple triangle) and reflect the European, African, and Hispanic ancestral populations of the GWAS results.

### Slide 10
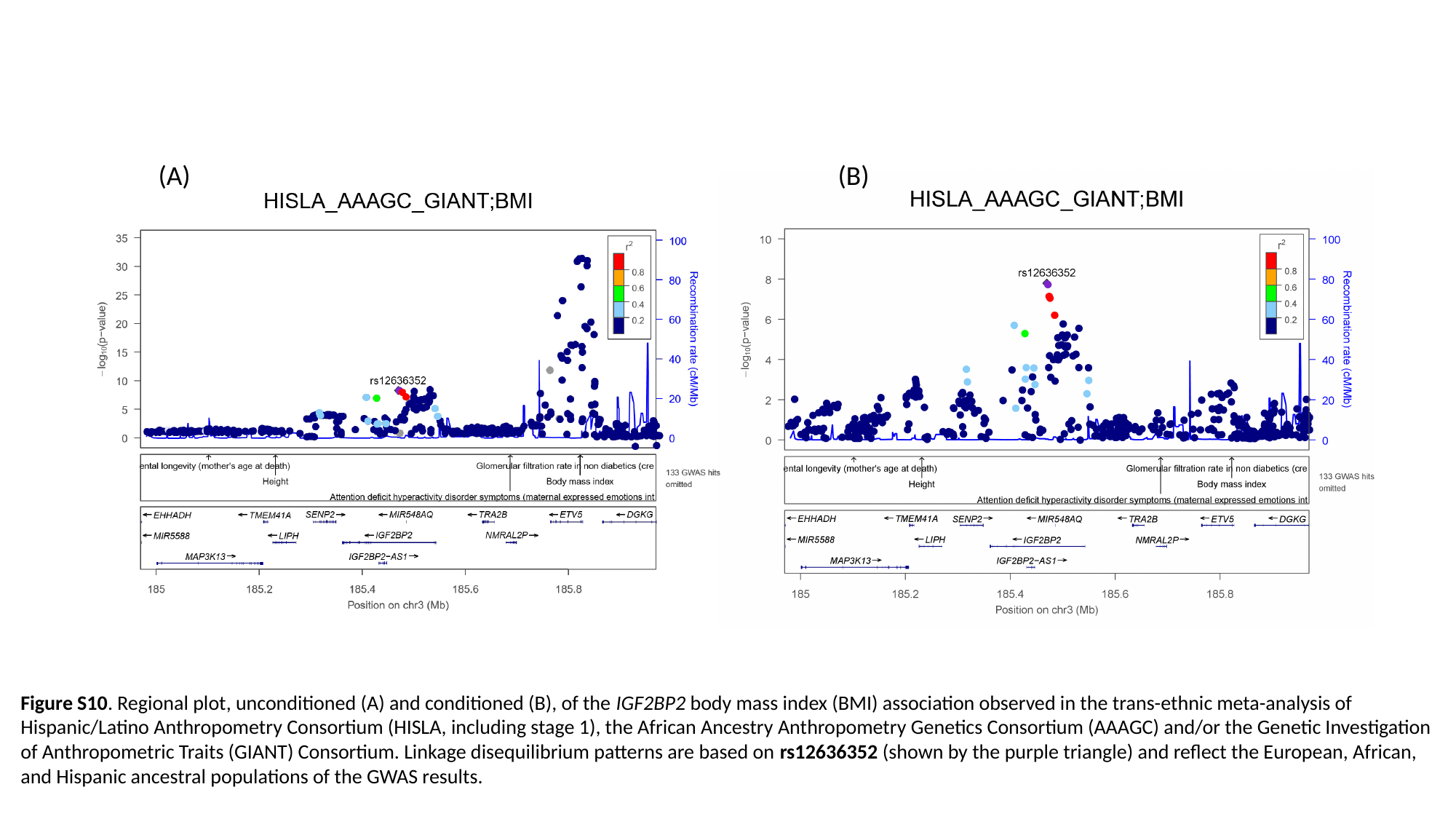

(A) (B)
Figure S10. Regional plot, unconditioned (A) and conditioned (B), of the IGF2BP2 body mass index (BMI) association observed in the trans-ethnic meta-analysis of Hispanic/Latino Anthropometry Consortium (HISLA, including stage 1), the African Ancestry Anthropometry Genetics Consortium (AAAGC) and/or the Genetic Investigation of Anthropometric Traits (GIANT) Consortium. Linkage disequilibrium patterns are based on rs12636352 (shown by the purple triangle) and reflect the European, African, and Hispanic ancestral populations of the GWAS results.

### Slide 11
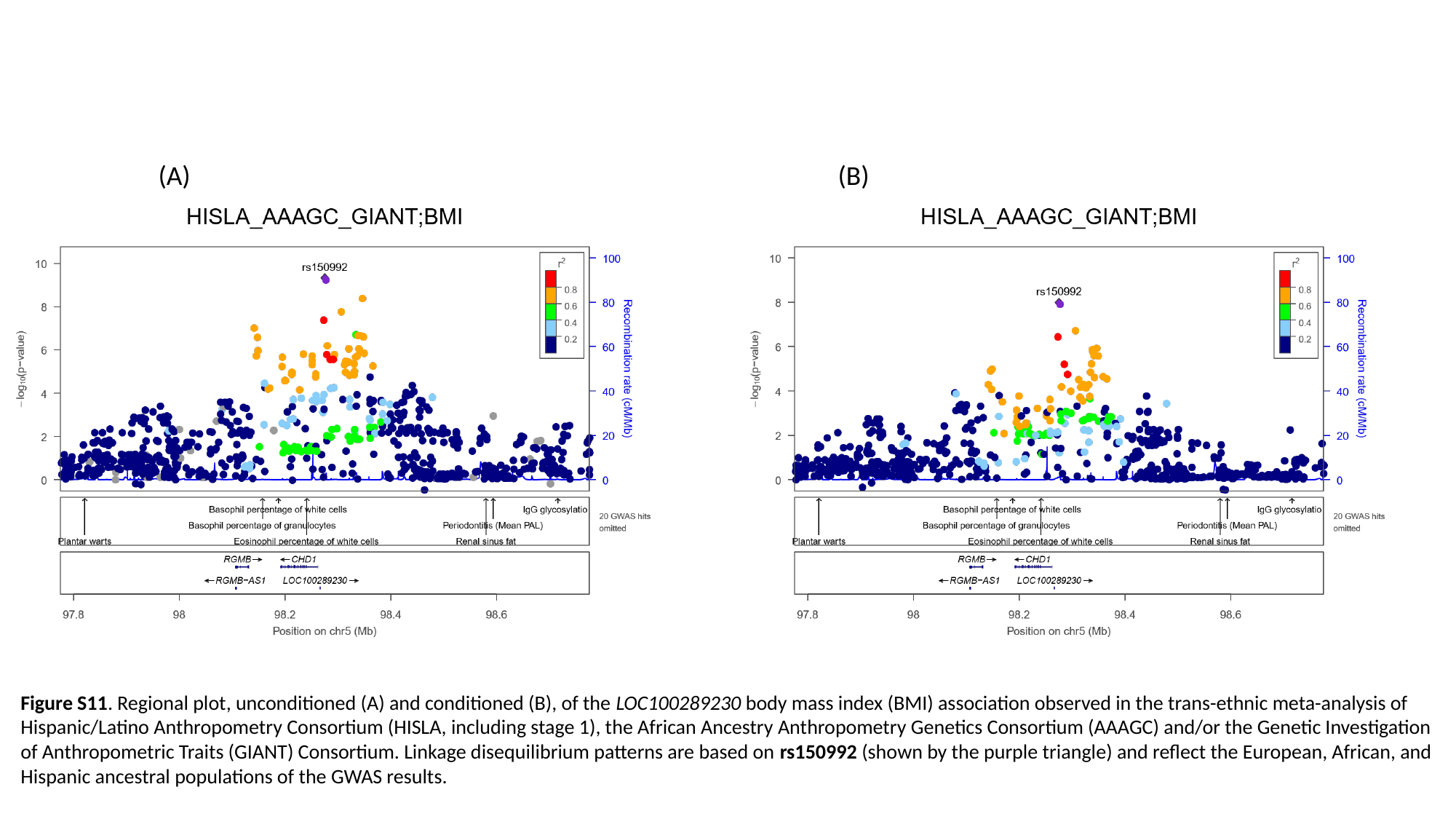

(A) (B)
Figure S11. Regional plot, unconditioned (A) and conditioned (B), of the LOC100289230 body mass index (BMI) association observed in the trans-ethnic meta-analysis of Hispanic/Latino Anthropometry Consortium (HISLA, including stage 1), the African Ancestry Anthropometry Genetics Consortium (AAAGC) and/or the Genetic Investigation of Anthropometric Traits (GIANT) Consortium. Linkage disequilibrium patterns are based on rs150992 (shown by the purple triangle) and reflect the European, African, and Hispanic ancestral populations of the GWAS results.

### Slide 12
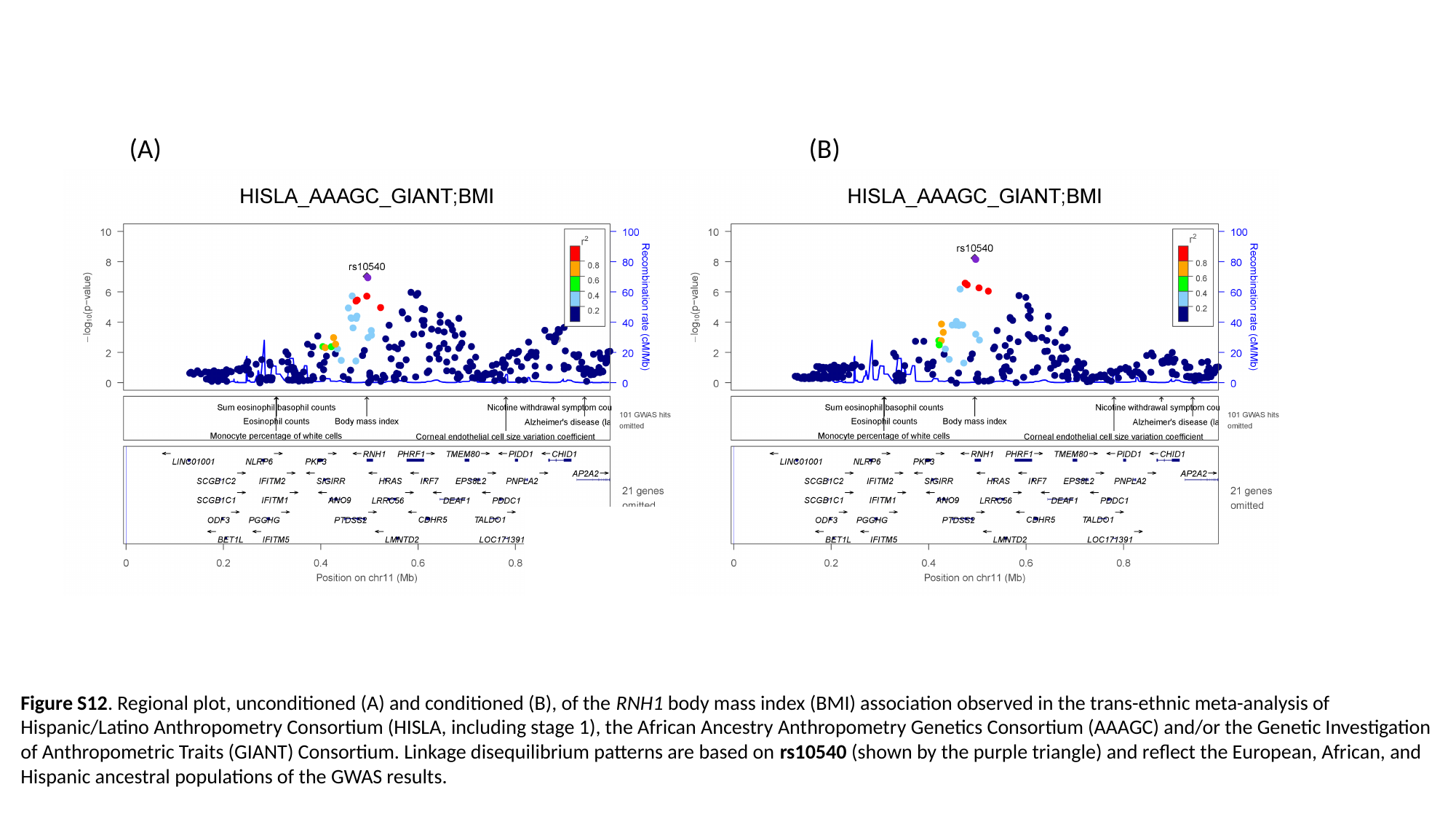

(A) (B)
Figure S12. Regional plot, unconditioned (A) and conditioned (B), of the RNH1 body mass index (BMI) association observed in the trans-ethnic meta-analysis of Hispanic/Latino Anthropometry Consortium (HISLA, including stage 1), the African Ancestry Anthropometry Genetics Consortium (AAAGC) and/or the Genetic Investigation of Anthropometric Traits (GIANT) Consortium. Linkage disequilibrium patterns are based on rs10540 (shown by the purple triangle) and reflect the European, African, and Hispanic ancestral populations of the GWAS results.

### Slide 13
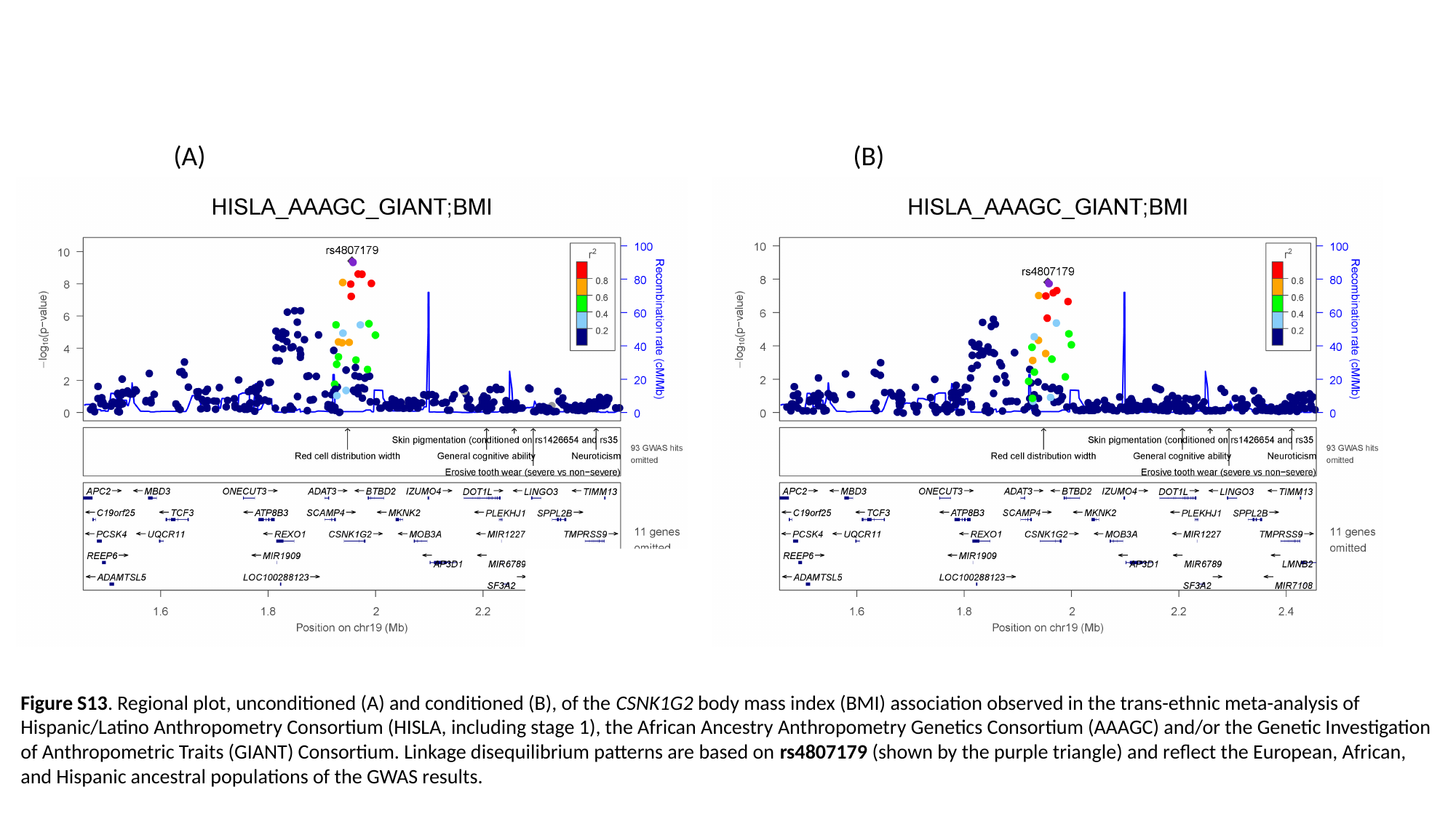

(A) (B)
Figure S13. Regional plot, unconditioned (A) and conditioned (B), of the CSNK1G2 body mass index (BMI) association observed in the trans-ethnic meta-analysis of Hispanic/Latino Anthropometry Consortium (HISLA, including stage 1), the African Ancestry Anthropometry Genetics Consortium (AAAGC) and/or the Genetic Investigation of Anthropometric Traits (GIANT) Consortium. Linkage disequilibrium patterns are based on rs4807179 (shown by the purple triangle) and reflect the European, African, and Hispanic ancestral populations of the GWAS results.

### Slide 14
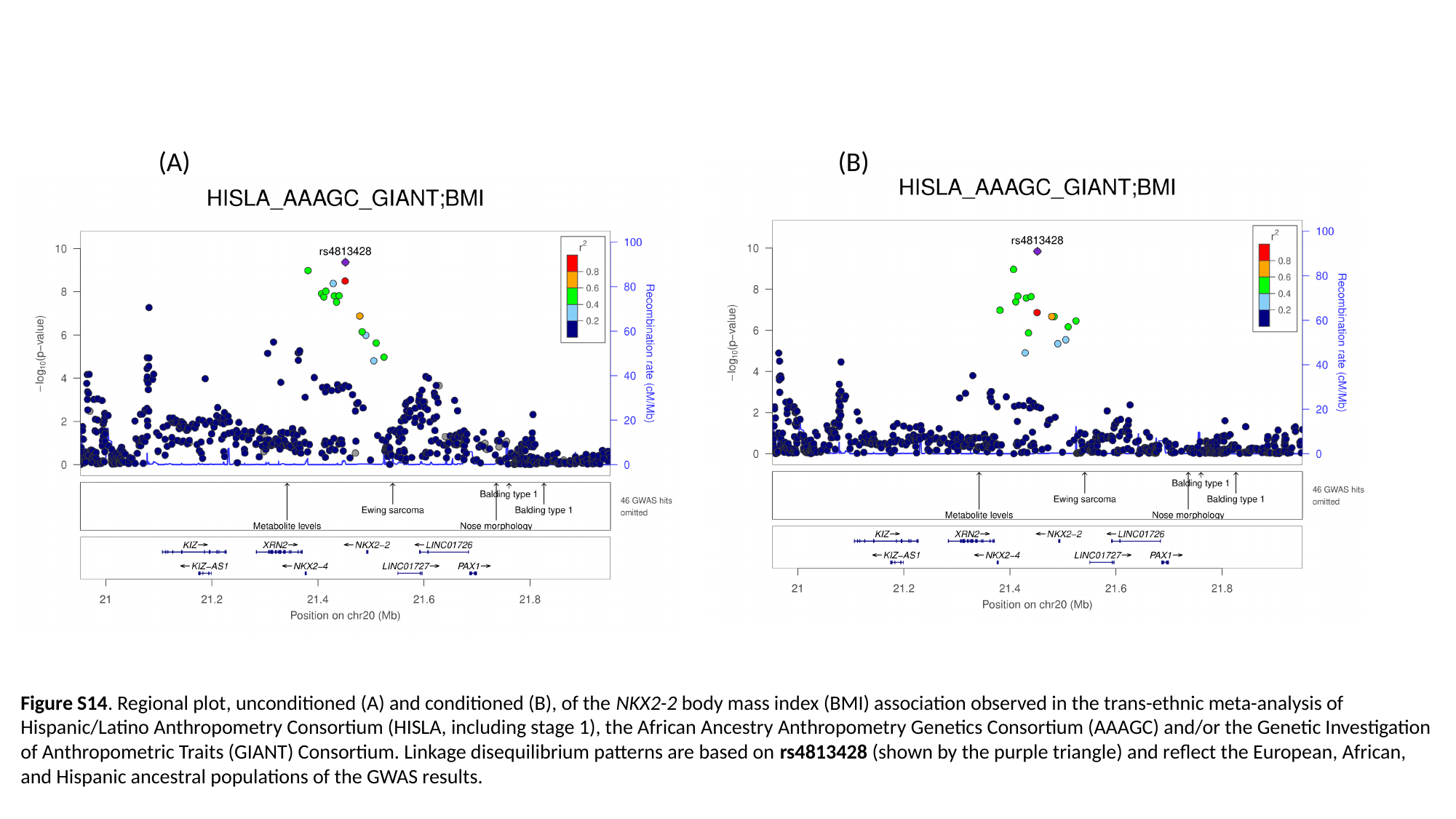

(A) (B)
Figure S14. Regional plot, unconditioned (A) and conditioned (B), of the NKX2-2 body mass index (BMI) association observed in the trans-ethnic meta-analysis of Hispanic/Latino Anthropometry Consortium (HISLA, including stage 1), the African Ancestry Anthropometry Genetics Consortium (AAAGC) and/or the Genetic Investigation of Anthropometric Traits (GIANT) Consortium. Linkage disequilibrium patterns are based on rs4813428 (shown by the purple triangle) and reflect the European, African, and Hispanic ancestral populations of the GWAS results.

### Slide 15
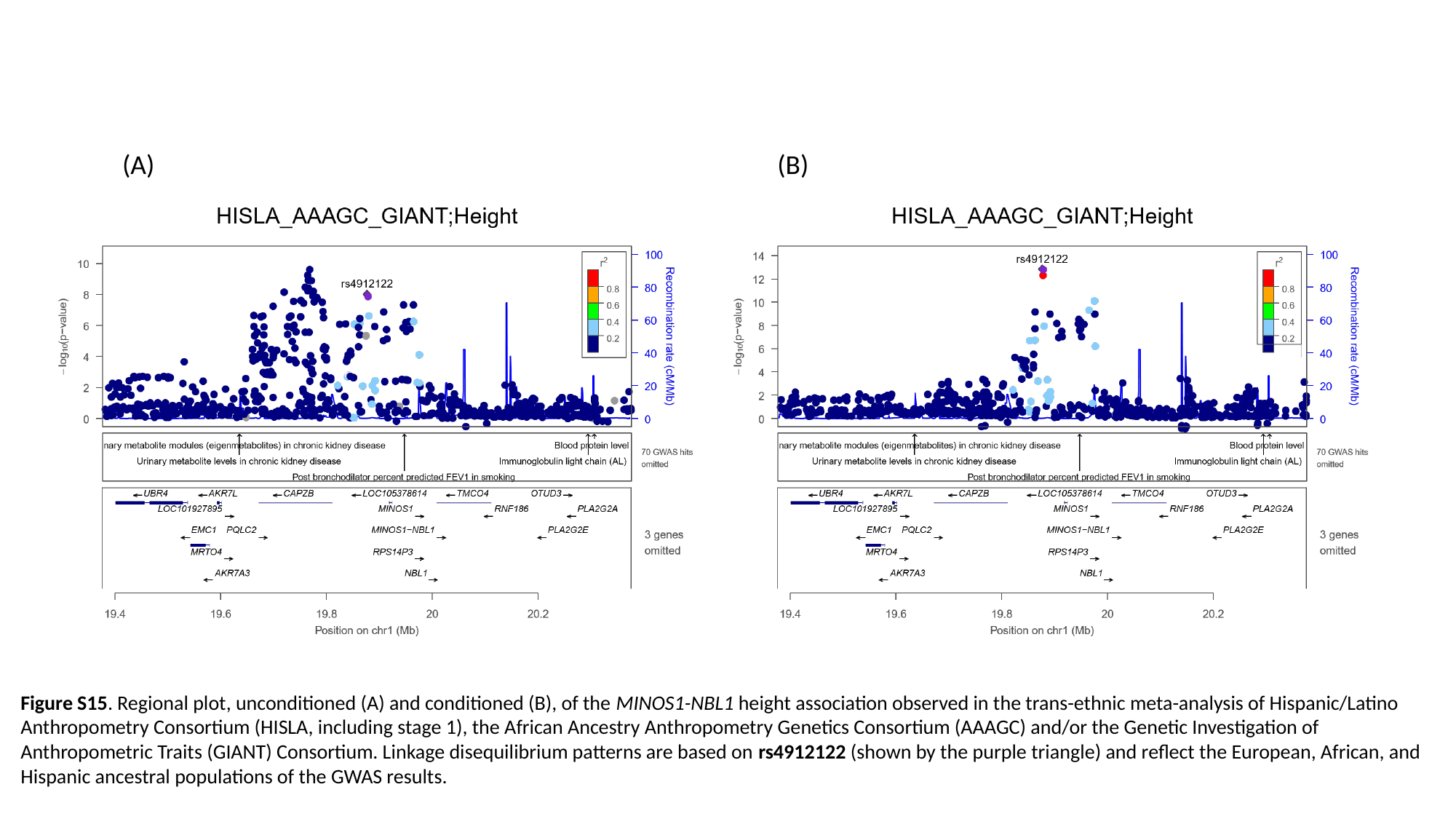

(A) (B)
Figure S15. Regional plot, unconditioned (A) and conditioned (B), of the MINOS1-NBL1 height association observed in the trans-ethnic meta-analysis of Hispanic/Latino Anthropometry Consortium (HISLA, including stage 1), the African Ancestry Anthropometry Genetics Consortium (AAAGC) and/or the Genetic Investigation of Anthropometric Traits (GIANT) Consortium. Linkage disequilibrium patterns are based on rs4912122 (shown by the purple triangle) and reflect the European, African, and Hispanic ancestral populations of the GWAS results.

### Slide 16
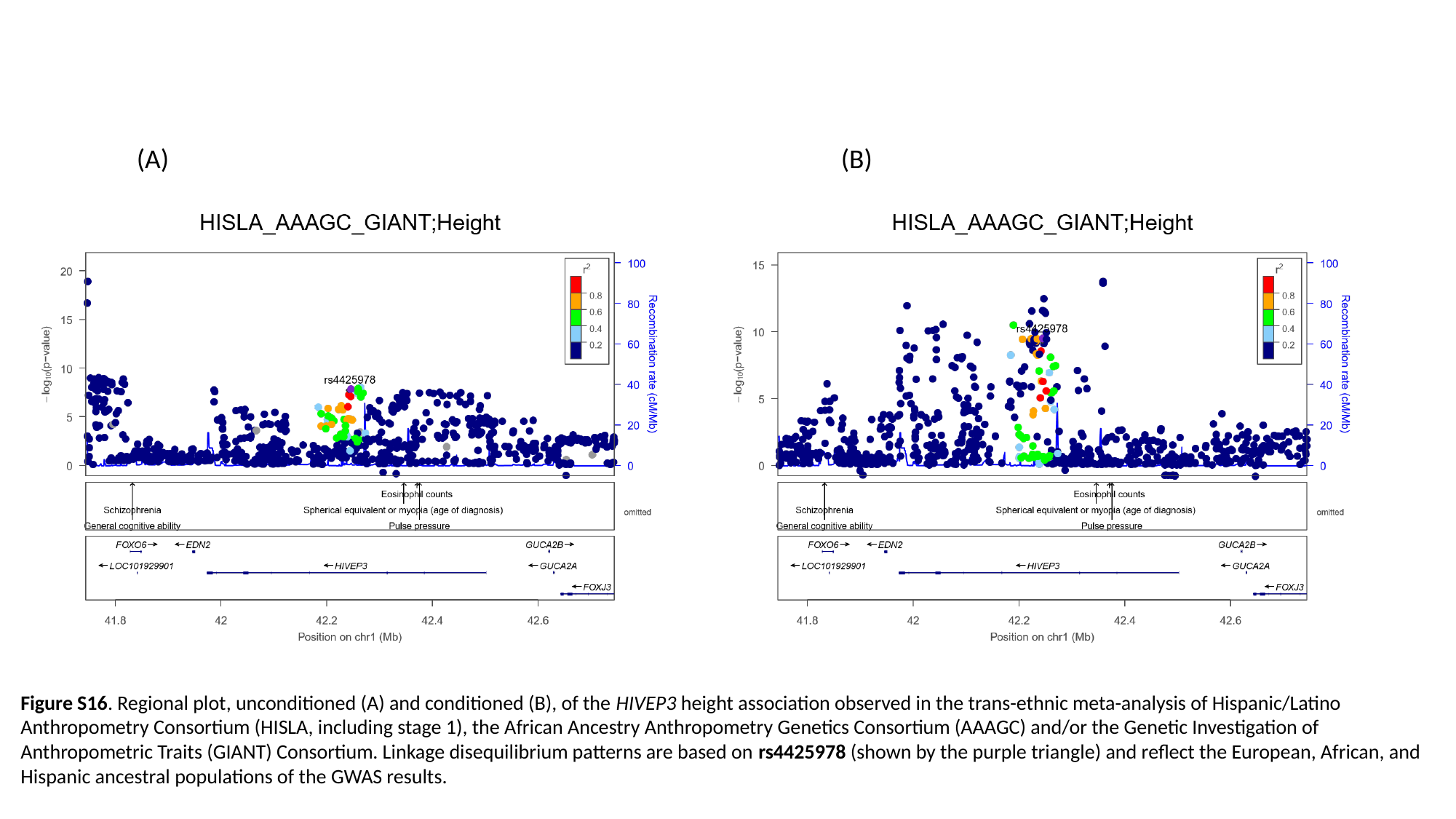

(A) (B)
Figure S16. Regional plot, unconditioned (A) and conditioned (B), of the HIVEP3 height association observed in the trans-ethnic meta-analysis of Hispanic/Latino Anthropometry Consortium (HISLA, including stage 1), the African Ancestry Anthropometry Genetics Consortium (AAAGC) and/or the Genetic Investigation of Anthropometric Traits (GIANT) Consortium. Linkage disequilibrium patterns are based on rs4425978 (shown by the purple triangle) and reflect the European, African, and Hispanic ancestral populations of the GWAS results.

### Slide 17
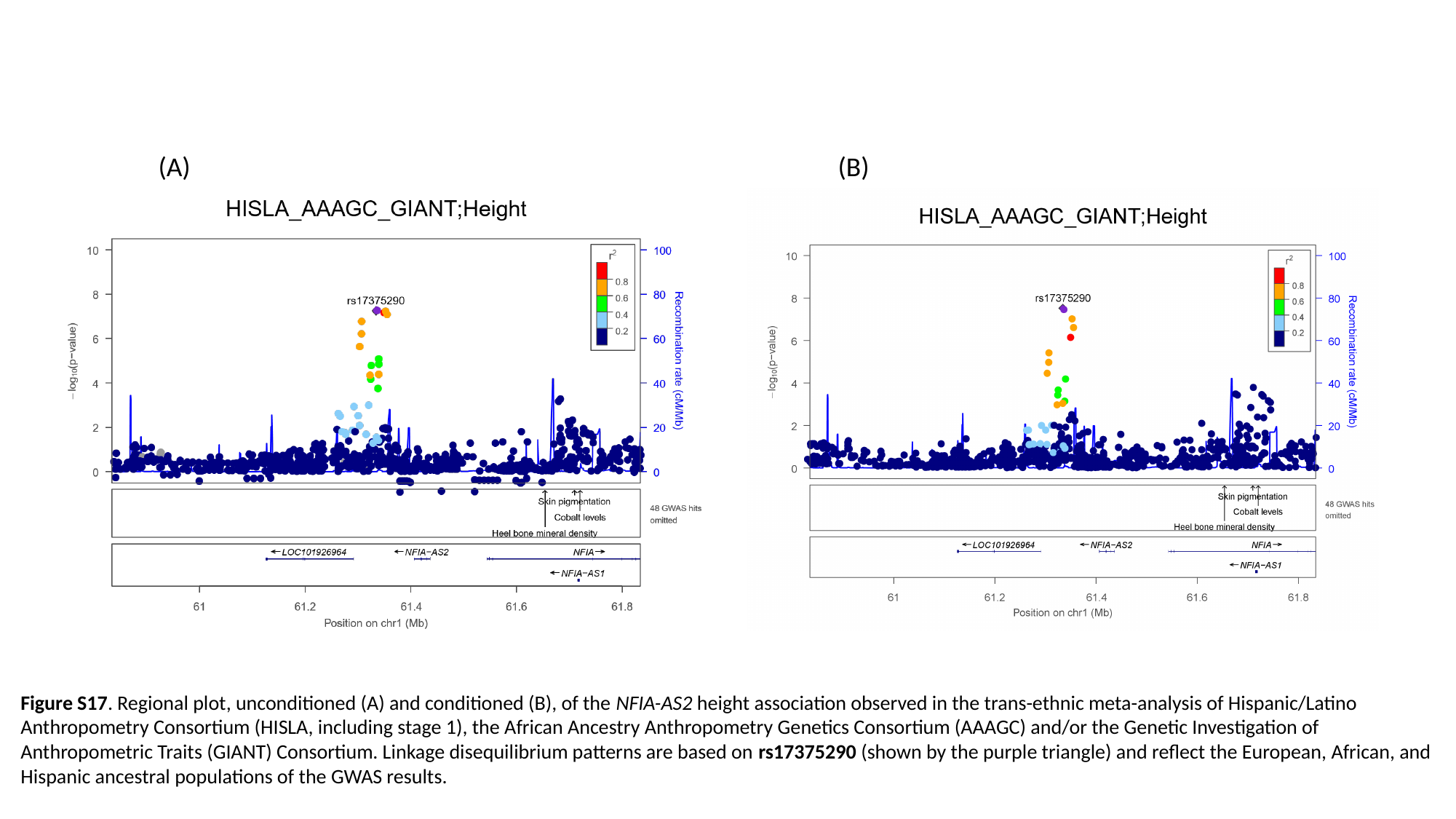

(A) (B)
Figure S17. Regional plot, unconditioned (A) and conditioned (B), of the NFIA-AS2 height association observed in the trans-ethnic meta-analysis of Hispanic/Latino Anthropometry Consortium (HISLA, including stage 1), the African Ancestry Anthropometry Genetics Consortium (AAAGC) and/or the Genetic Investigation of Anthropometric Traits (GIANT) Consortium. Linkage disequilibrium patterns are based on rs17375290 (shown by the purple triangle) and reflect the European, African, and Hispanic ancestral populations of the GWAS results.

### Slide 18
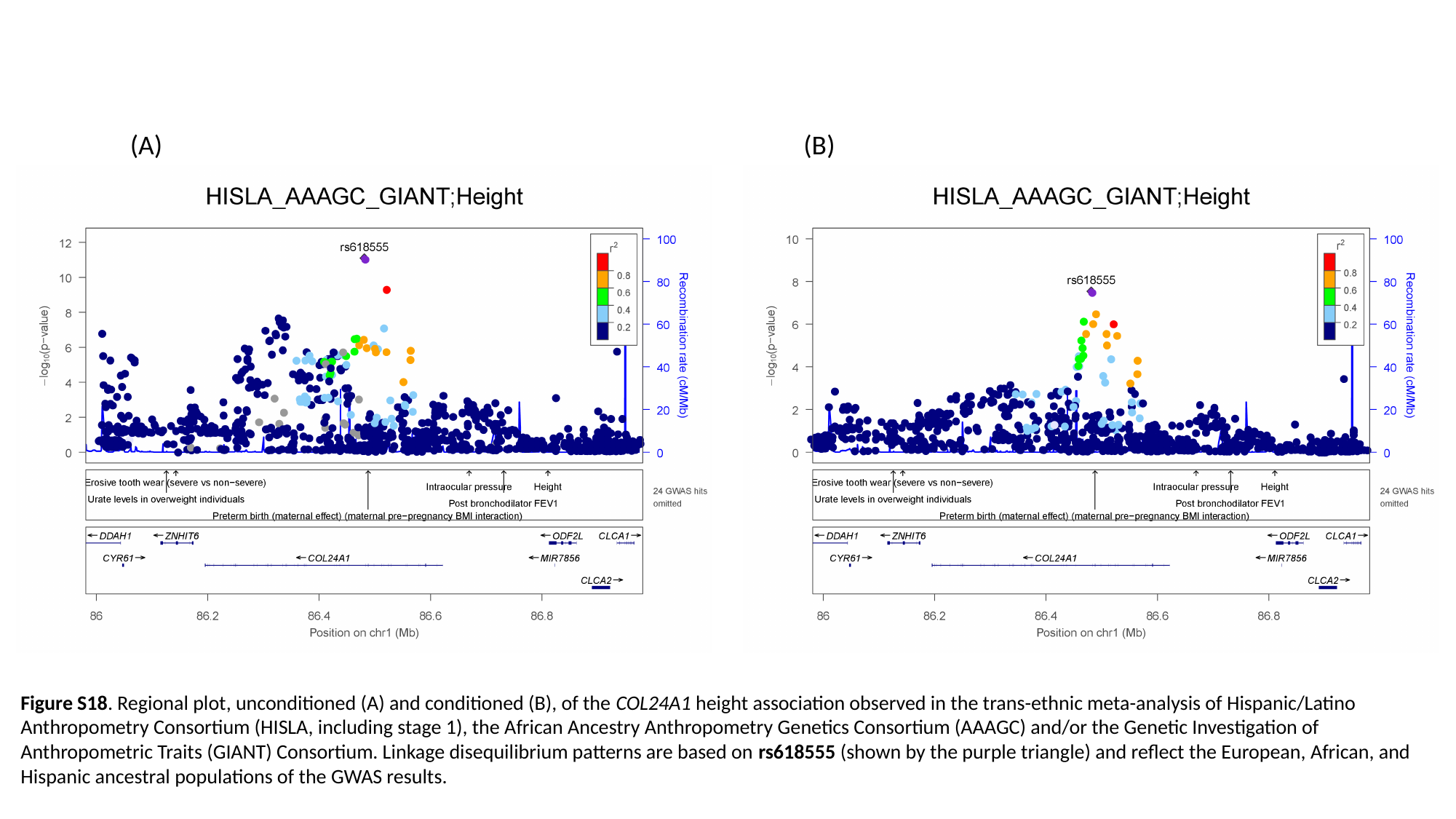

(A) (B)
Figure S18. Regional plot, unconditioned (A) and conditioned (B), of the COL24A1 height association observed in the trans-ethnic meta-analysis of Hispanic/Latino Anthropometry Consortium (HISLA, including stage 1), the African Ancestry Anthropometry Genetics Consortium (AAAGC) and/or the Genetic Investigation of Anthropometric Traits (GIANT) Consortium. Linkage disequilibrium patterns are based on rs618555 (shown by the purple triangle) and reflect the European, African, and Hispanic ancestral populations of the GWAS results.

### Slide 19
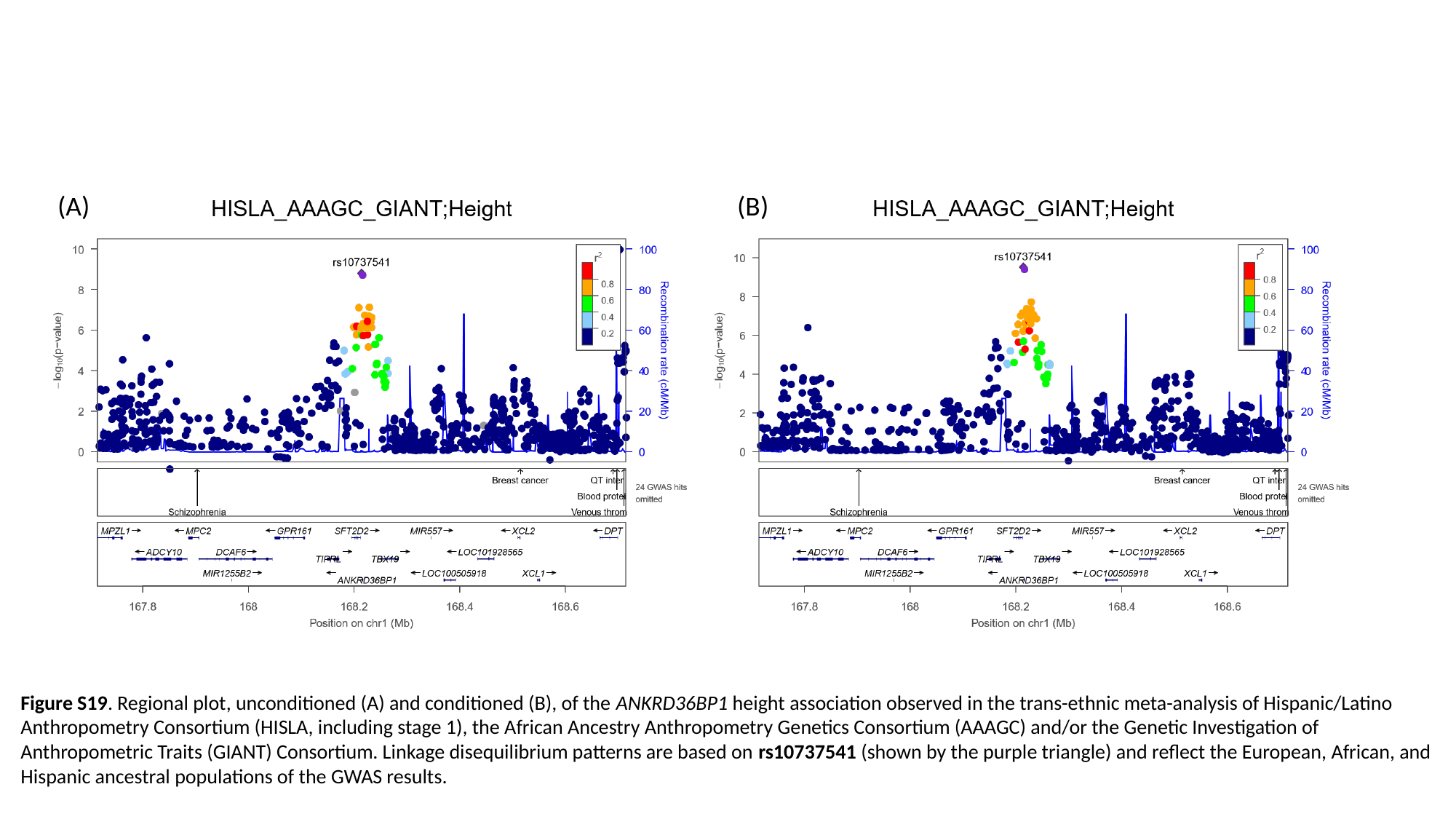

(A) (B)
Figure S19. Regional plot, unconditioned (A) and conditioned (B), of the ANKRD36BP1 height association observed in the trans-ethnic meta-analysis of Hispanic/Latino Anthropometry Consortium (HISLA, including stage 1), the African Ancestry Anthropometry Genetics Consortium (AAAGC) and/or the Genetic Investigation of Anthropometric Traits (GIANT) Consortium. Linkage disequilibrium patterns are based on rs10737541 (shown by the purple triangle) and reflect the European, African, and Hispanic ancestral populations of the GWAS results.

### Slide 20
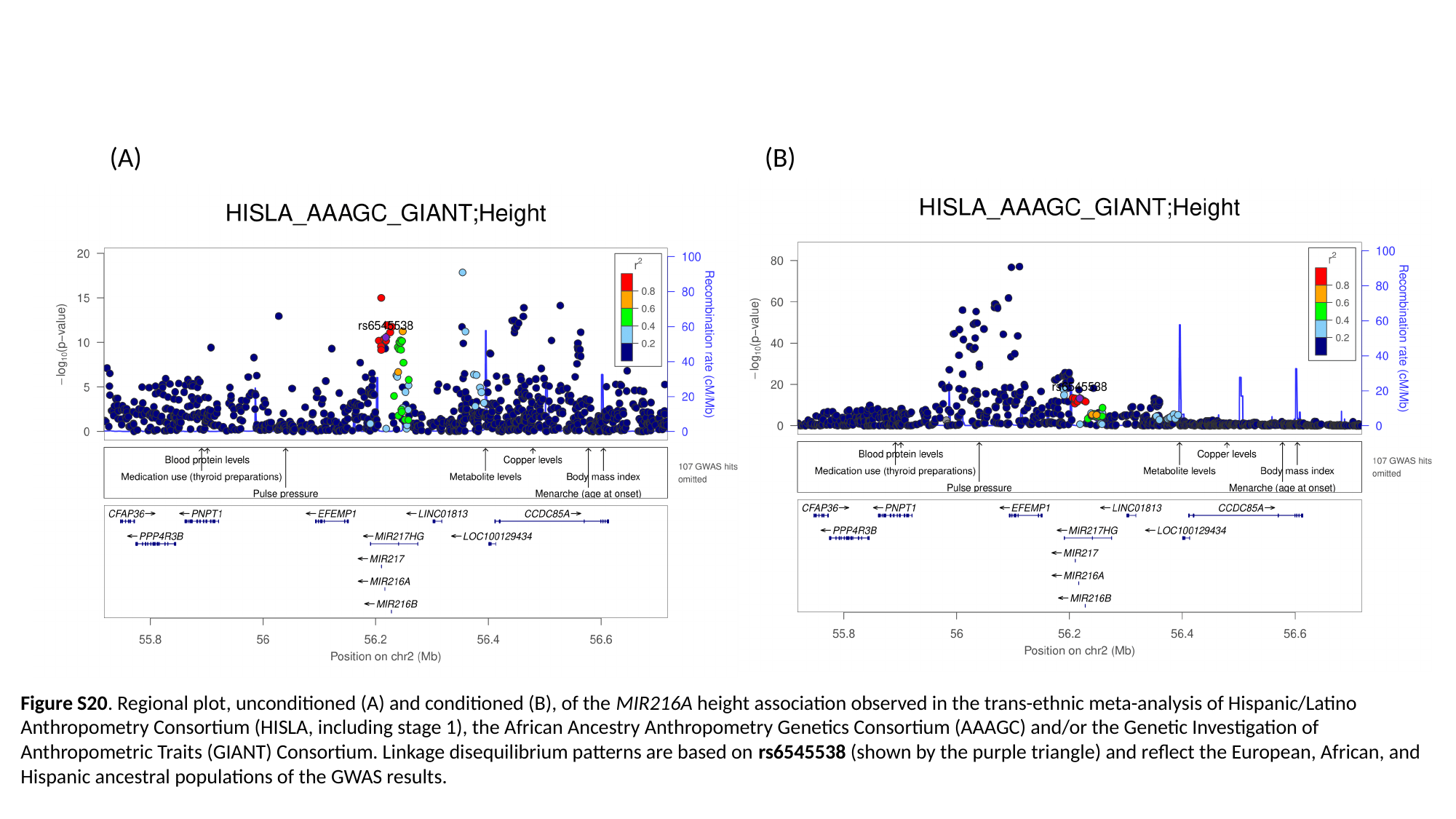

(A) (B)
Figure S20. Regional plot, unconditioned (A) and conditioned (B), of the MIR216A height association observed in the trans-ethnic meta-analysis of Hispanic/Latino Anthropometry Consortium (HISLA, including stage 1), the African Ancestry Anthropometry Genetics Consortium (AAAGC) and/or the Genetic Investigation of Anthropometric Traits (GIANT) Consortium. Linkage disequilibrium patterns are based on rs6545538 (shown by the purple triangle) and reflect the European, African, and Hispanic ancestral populations of the GWAS results.

### Slide 21
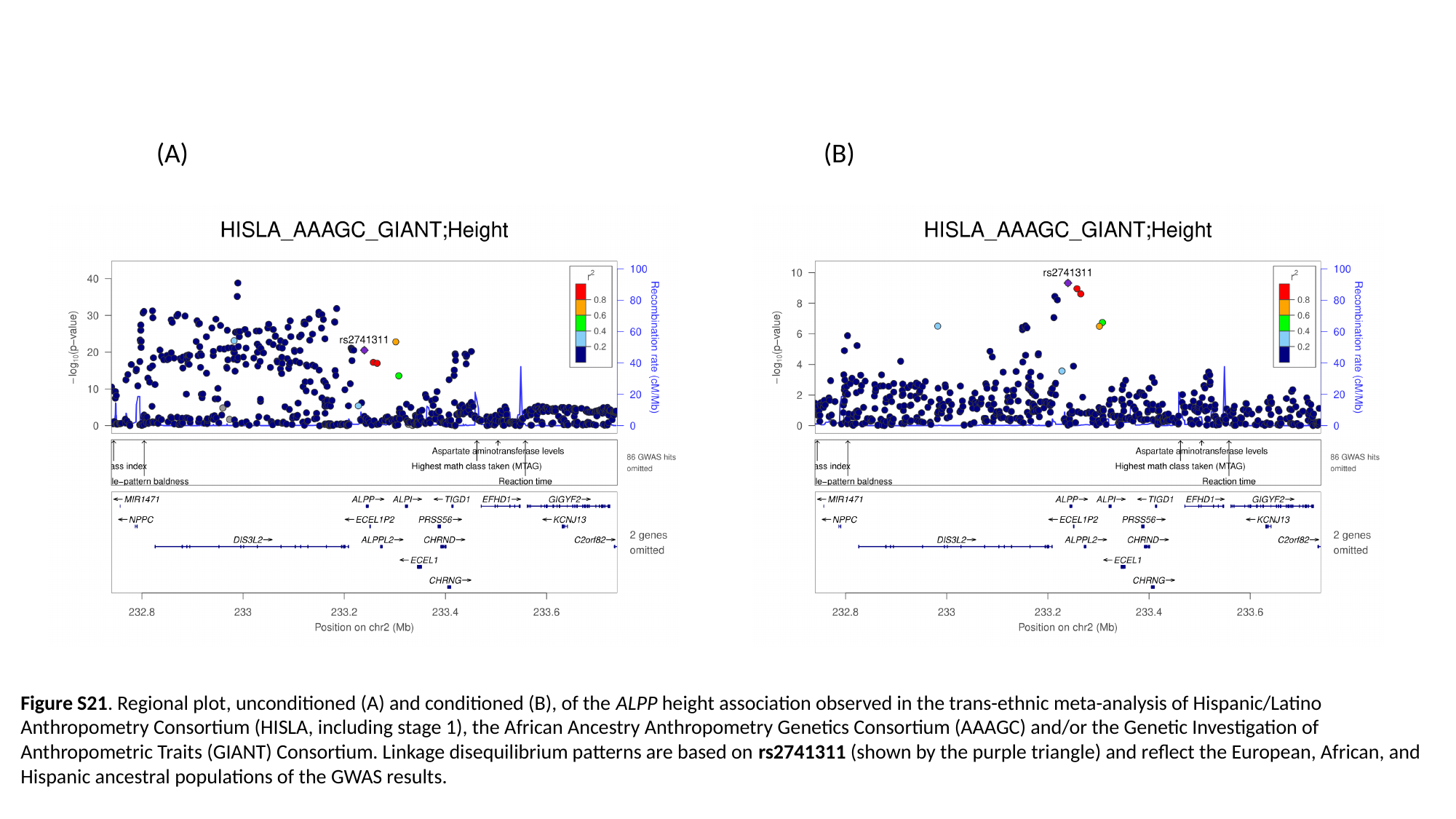

(A) (B)
Figure S21. Regional plot, unconditioned (A) and conditioned (B), of the ALPP height association observed in the trans-ethnic meta-analysis of Hispanic/Latino Anthropometry Consortium (HISLA, including stage 1), the African Ancestry Anthropometry Genetics Consortium (AAAGC) and/or the Genetic Investigation of Anthropometric Traits (GIANT) Consortium. Linkage disequilibrium patterns are based on rs2741311 (shown by the purple triangle) and reflect the European, African, and Hispanic ancestral populations of the GWAS results.

### Slide 22
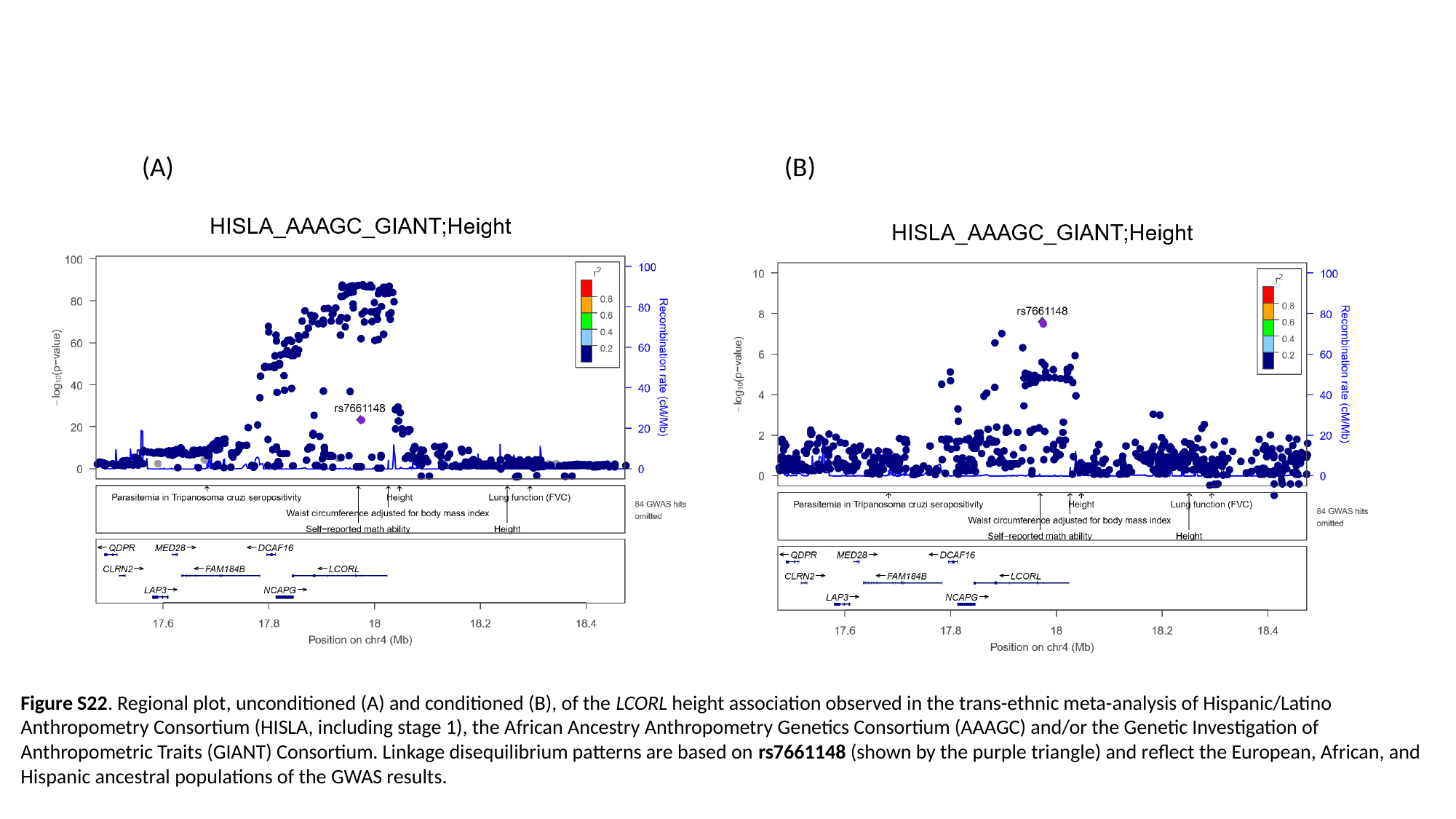

(A) (B)
Figure S22. Regional plot, unconditioned (A) and conditioned (B), of the LCORL height association observed in the trans-ethnic meta-analysis of Hispanic/Latino Anthropometry Consortium (HISLA, including stage 1), the African Ancestry Anthropometry Genetics Consortium (AAAGC) and/or the Genetic Investigation of Anthropometric Traits (GIANT) Consortium. Linkage disequilibrium patterns are based on rs7661148 (shown by the purple triangle) and reflect the European, African, and Hispanic ancestral populations of the GWAS results.

### Slide 23
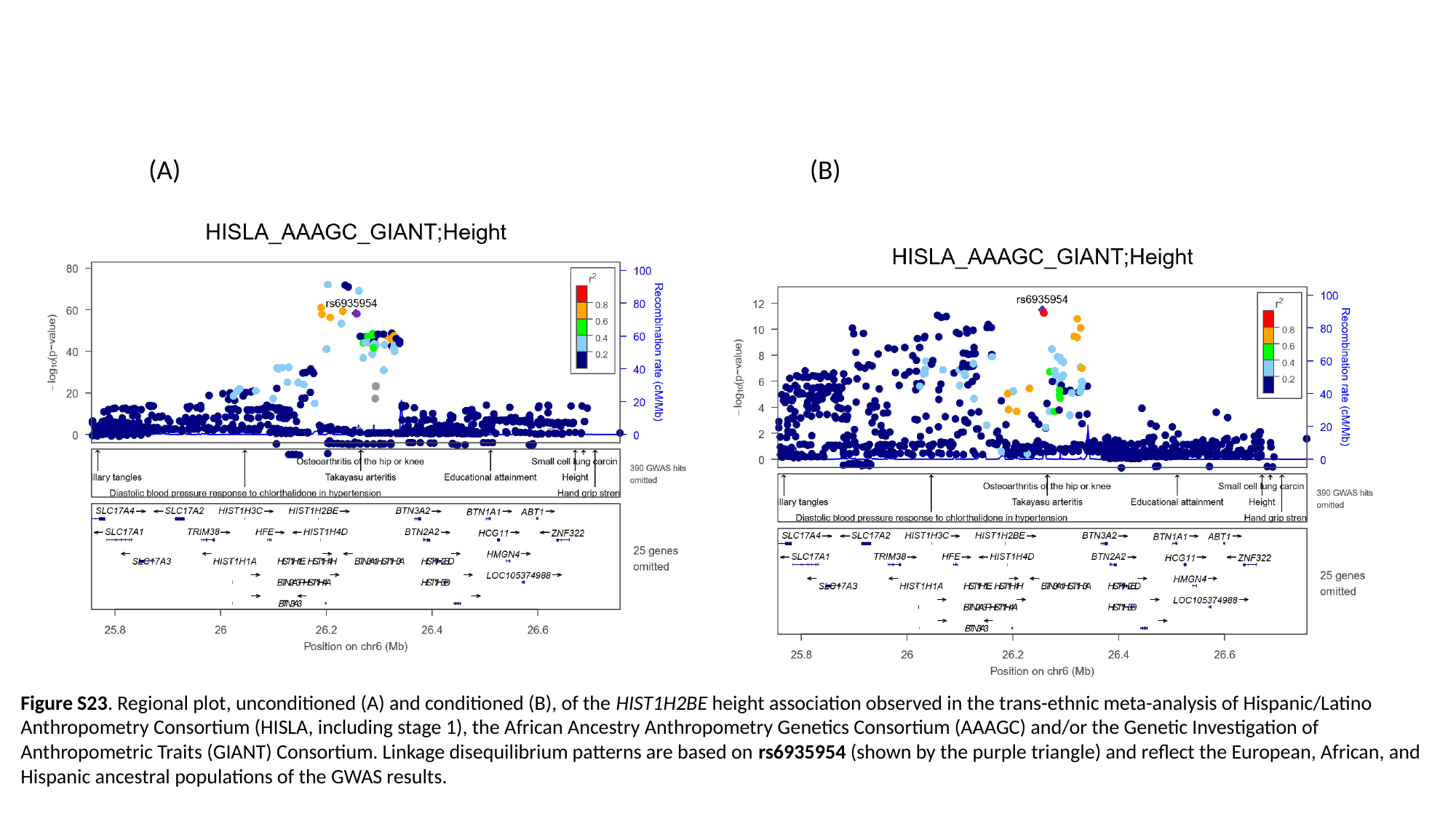

(A) (B)
Figure S23. Regional plot, unconditioned (A) and conditioned (B), of the HIST1H2BE height association observed in the trans-ethnic meta-analysis of Hispanic/Latino Anthropometry Consortium (HISLA, including stage 1), the African Ancestry Anthropometry Genetics Consortium (AAAGC) and/or the Genetic Investigation of Anthropometric Traits (GIANT) Consortium. Linkage disequilibrium patterns are based on rs6935954 (shown by the purple triangle) and reflect the European, African, and Hispanic ancestral populations of the GWAS results.

### Slide 24
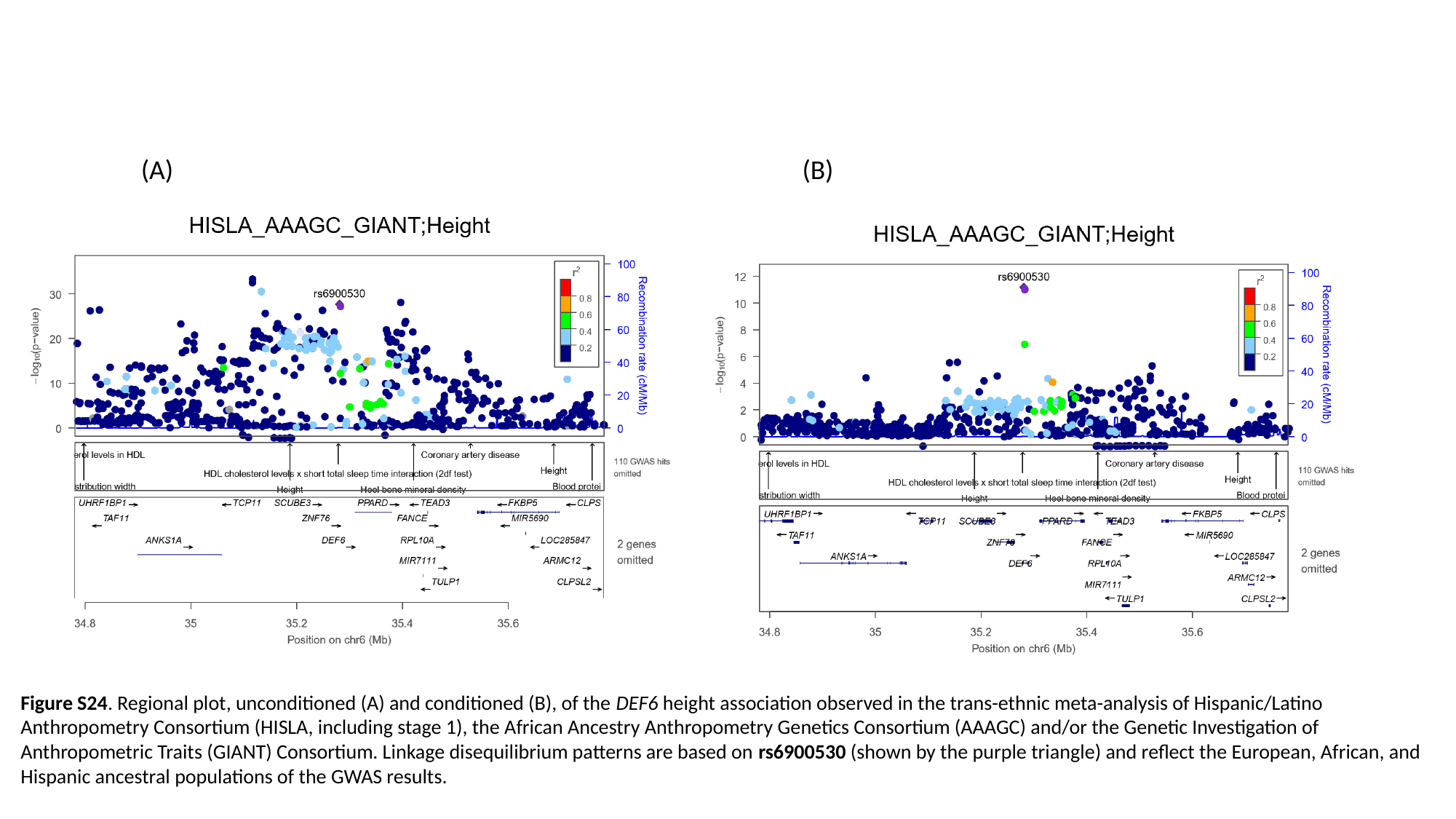

(A) (B)
Figure S24. Regional plot, unconditioned (A) and conditioned (B), of the DEF6 height association observed in the trans-ethnic meta-analysis of Hispanic/Latino Anthropometry Consortium (HISLA, including stage 1), the African Ancestry Anthropometry Genetics Consortium (AAAGC) and/or the Genetic Investigation of Anthropometric Traits (GIANT) Consortium. Linkage disequilibrium patterns are based on rs6900530 (shown by the purple triangle) and reflect the European, African, and Hispanic ancestral populations of the GWAS results.

### Slide 25
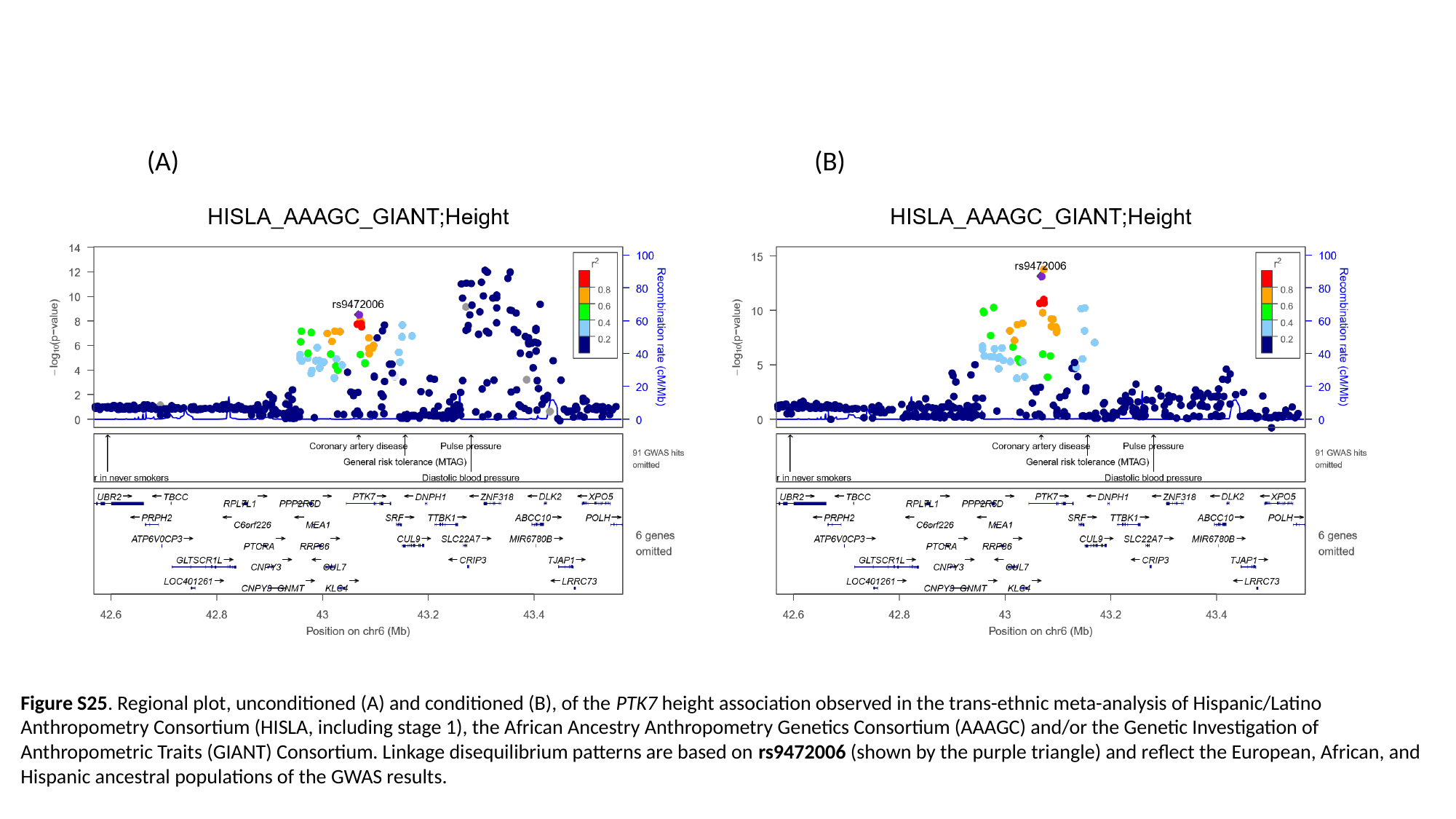

(A) (B)
Figure S25. Regional plot, unconditioned (A) and conditioned (B), of the PTK7 height association observed in the trans-ethnic meta-analysis of Hispanic/Latino Anthropometry Consortium (HISLA, including stage 1), the African Ancestry Anthropometry Genetics Consortium (AAAGC) and/or the Genetic Investigation of Anthropometric Traits (GIANT) Consortium. Linkage disequilibrium patterns are based on rs9472006 (shown by the purple triangle) and reflect the European, African, and Hispanic ancestral populations of the GWAS results.

### Slide 26
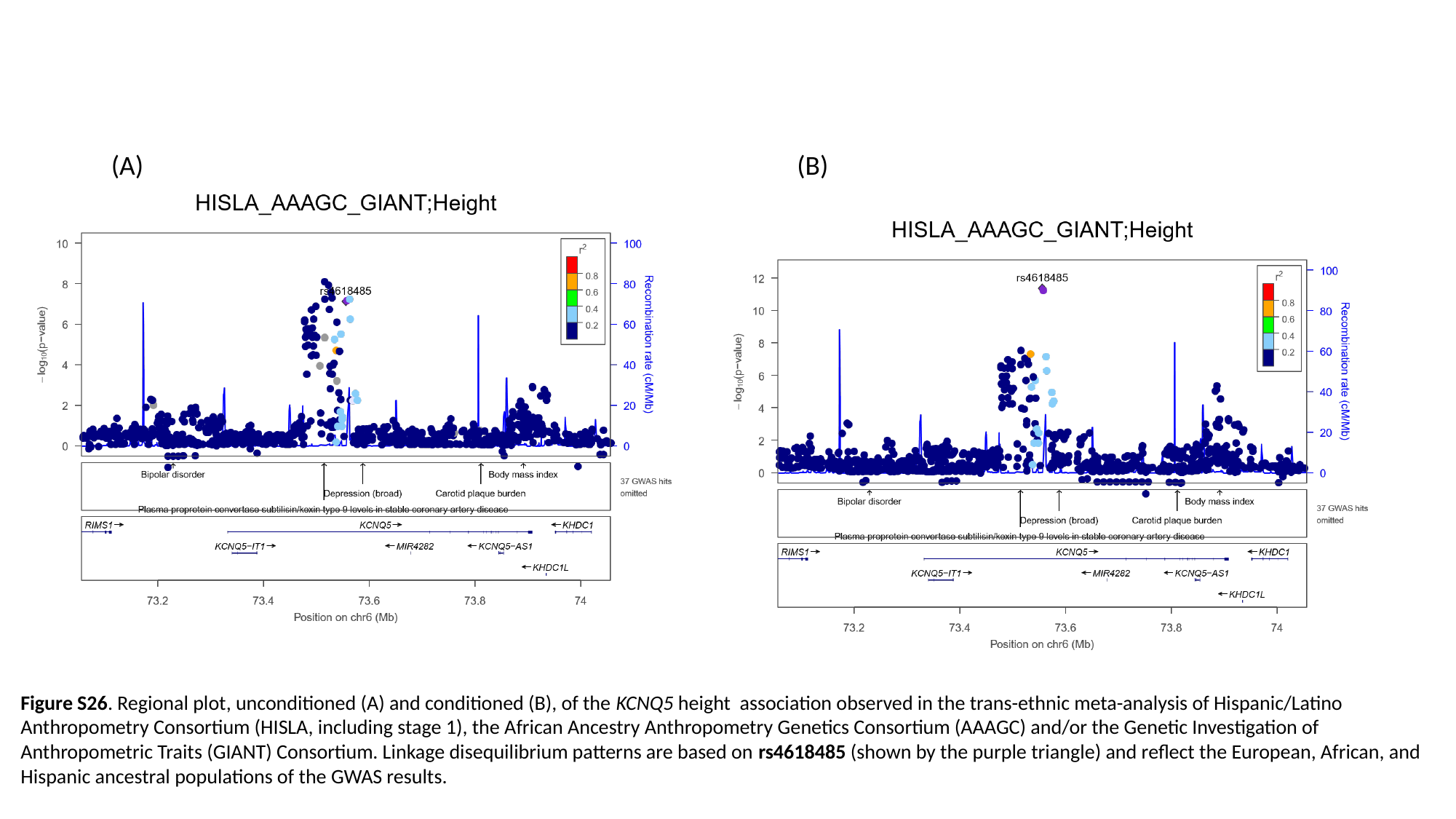

(A) (B)
Figure S26. Regional plot, unconditioned (A) and conditioned (B), of the KCNQ5 height association observed in the trans-ethnic meta-analysis of Hispanic/Latino Anthropometry Consortium (HISLA, including stage 1), the African Ancestry Anthropometry Genetics Consortium (AAAGC) and/or the Genetic Investigation of Anthropometric Traits (GIANT) Consortium. Linkage disequilibrium patterns are based on rs4618485 (shown by the purple triangle) and reflect the European, African, and Hispanic ancestral populations of the GWAS results.

### Slide 27
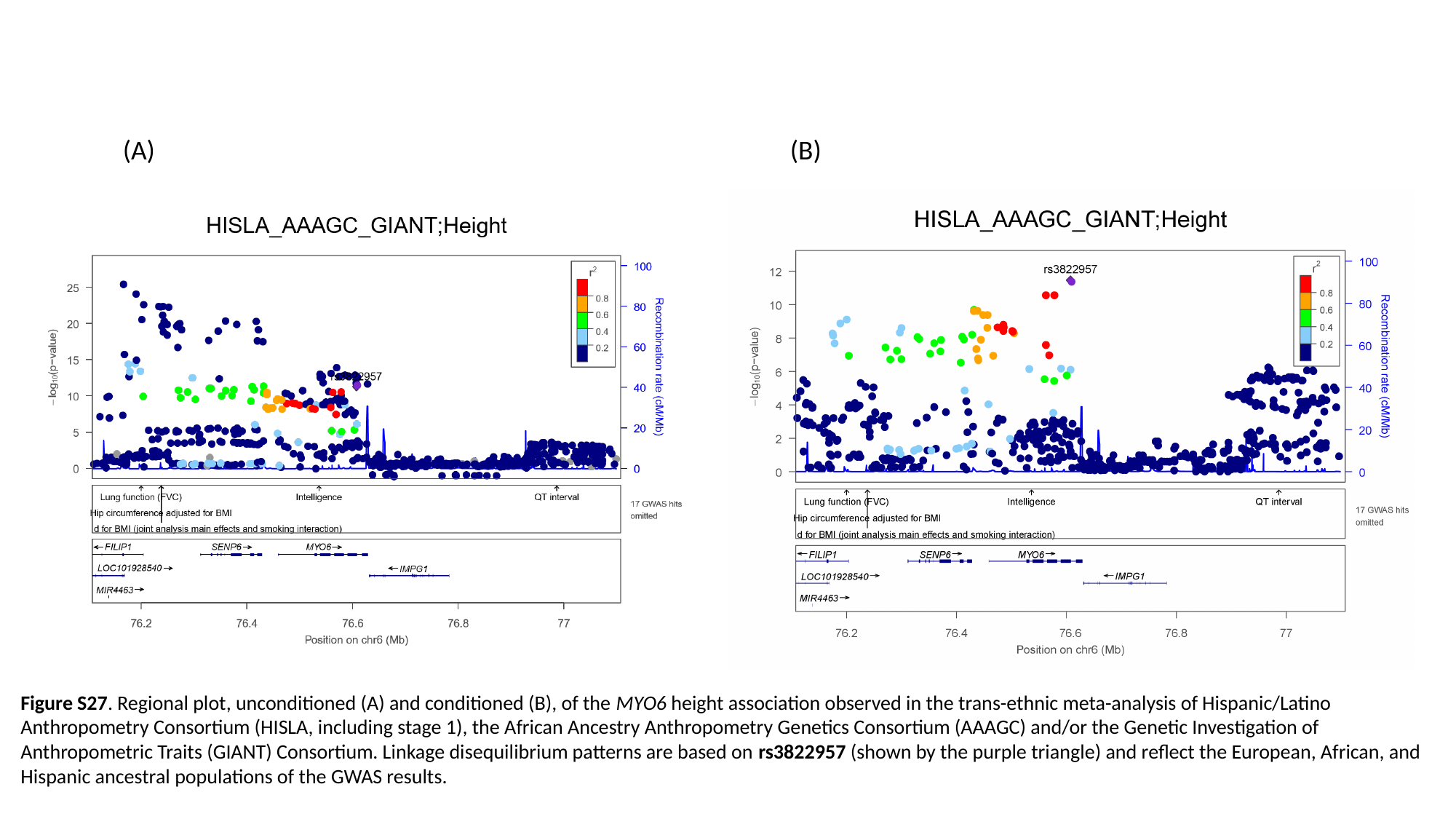

(A) (B)
Figure S27. Regional plot, unconditioned (A) and conditioned (B), of the MYO6 height association observed in the trans-ethnic meta-analysis of Hispanic/Latino Anthropometry Consortium (HISLA, including stage 1), the African Ancestry Anthropometry Genetics Consortium (AAAGC) and/or the Genetic Investigation of Anthropometric Traits (GIANT) Consortium. Linkage disequilibrium patterns are based on rs3822957 (shown by the purple triangle) and reflect the European, African, and Hispanic ancestral populations of the GWAS results.

### Slide 28
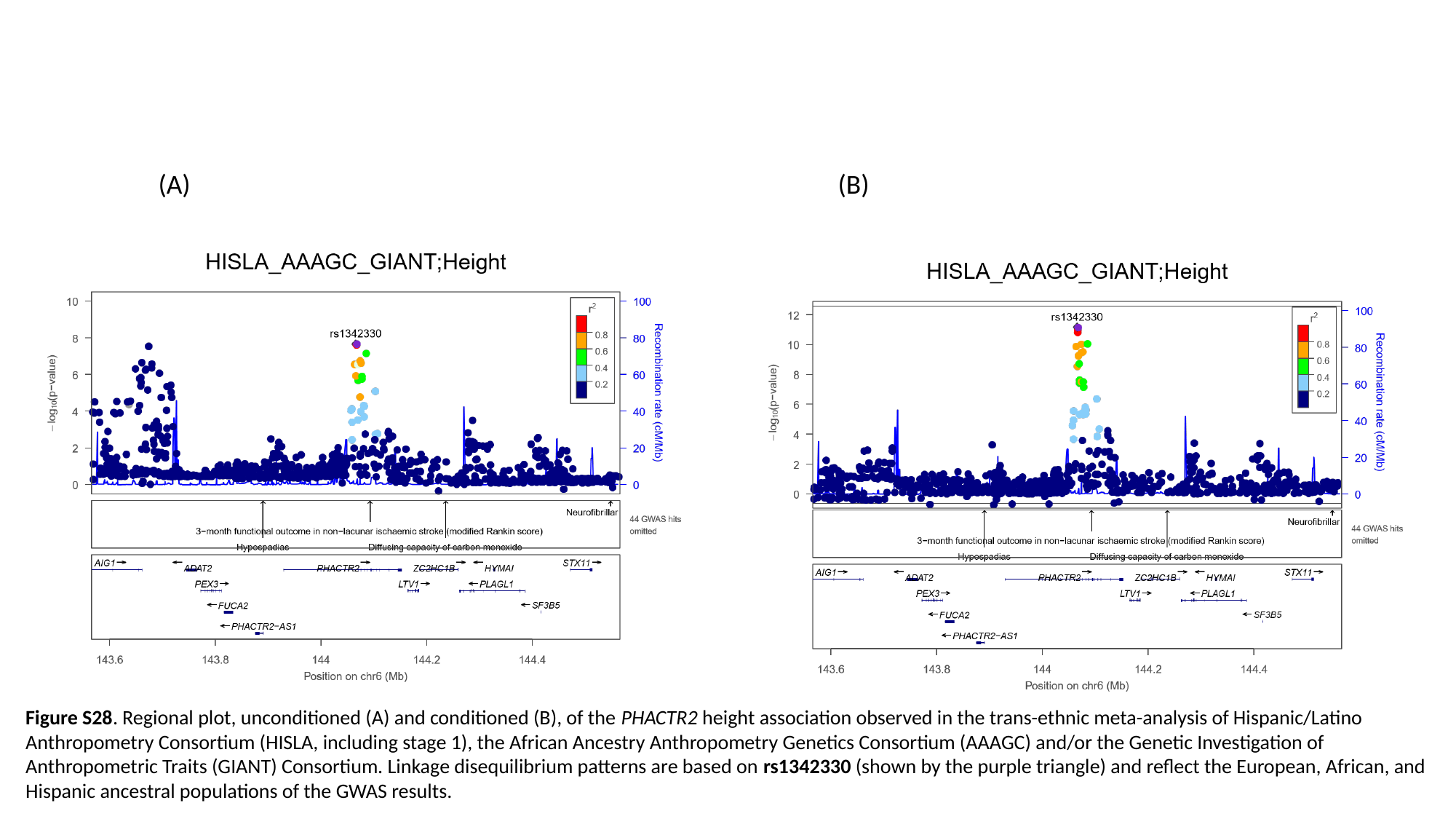

(A) (B)
Figure S28. Regional plot, unconditioned (A) and conditioned (B), of the PHACTR2 height association observed in the trans-ethnic meta-analysis of Hispanic/Latino Anthropometry Consortium (HISLA, including stage 1), the African Ancestry Anthropometry Genetics Consortium (AAAGC) and/or the Genetic Investigation of Anthropometric Traits (GIANT) Consortium. Linkage disequilibrium patterns are based on rs1342330 (shown by the purple triangle) and reflect the European, African, and Hispanic ancestral populations of the GWAS results.

### Slide 29
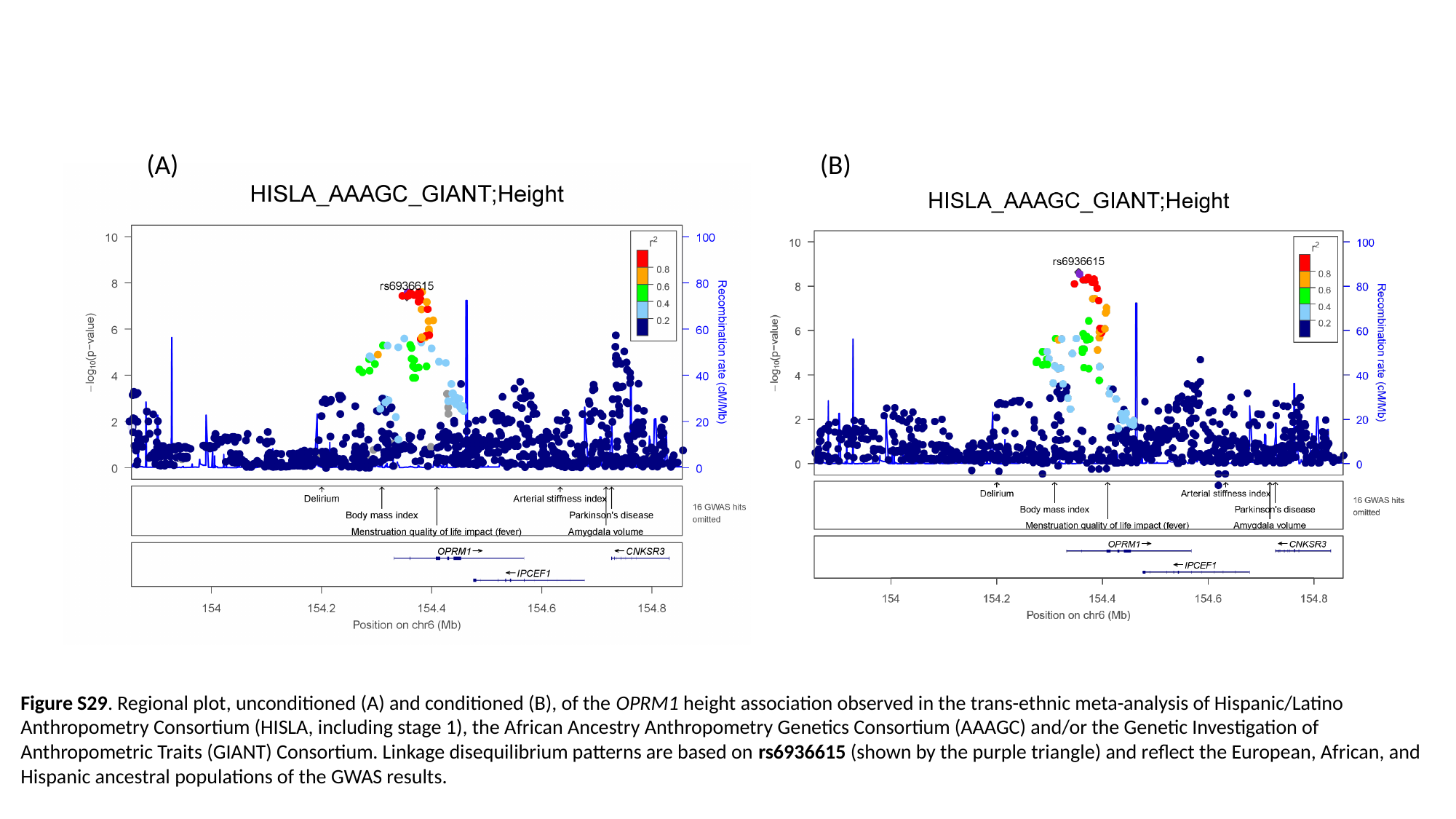

(A) (B)
Figure S29. Regional plot, unconditioned (A) and conditioned (B), of the OPRM1 height association observed in the trans-ethnic meta-analysis of Hispanic/Latino Anthropometry Consortium (HISLA, including stage 1), the African Ancestry Anthropometry Genetics Consortium (AAAGC) and/or the Genetic Investigation of Anthropometric Traits (GIANT) Consortium. Linkage disequilibrium patterns are based on rs6936615 (shown by the purple triangle) and reflect the European, African, and Hispanic ancestral populations of the GWAS results.

### Slide 30
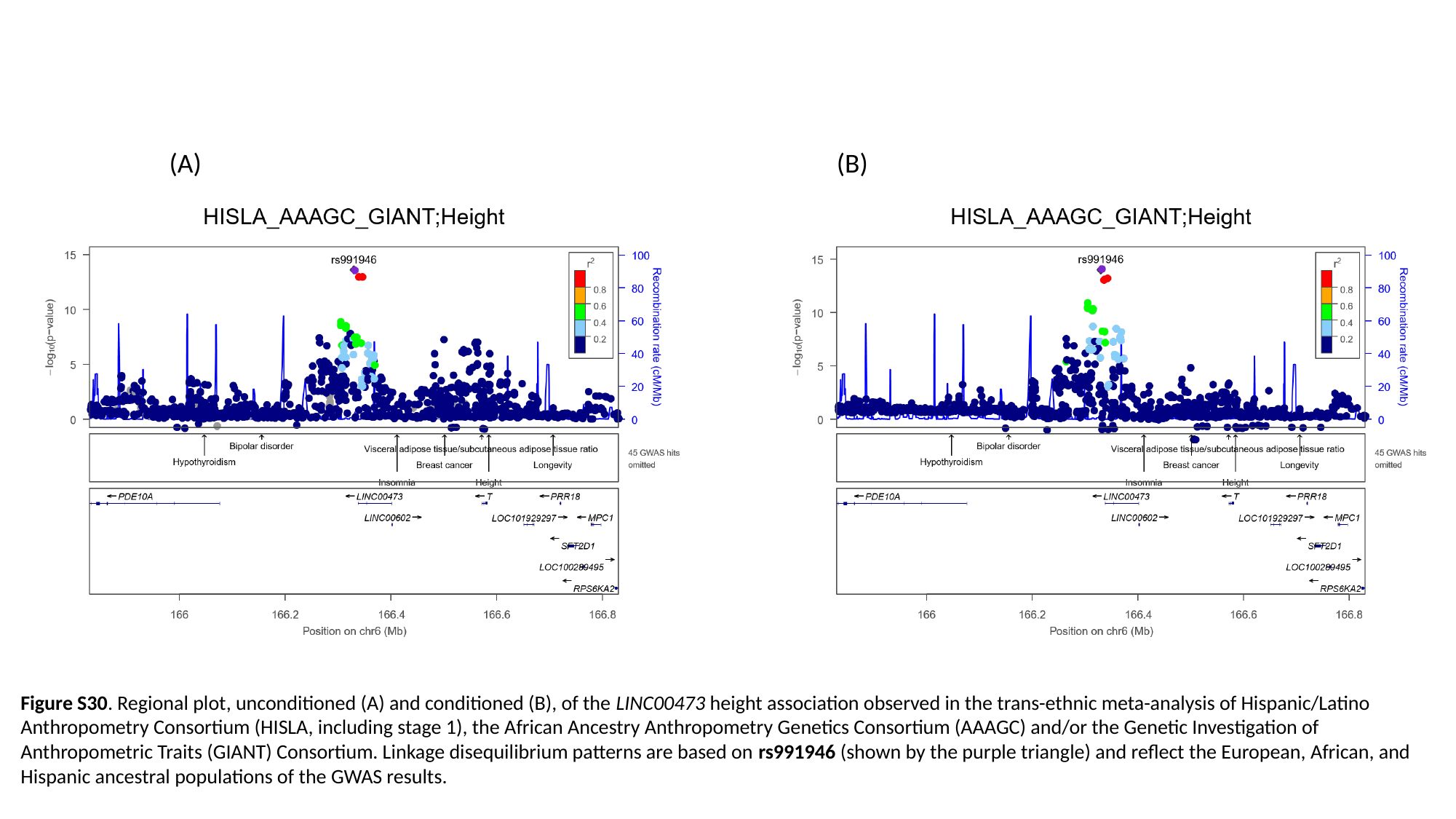

(A) (B)
Figure S30. Regional plot, unconditioned (A) and conditioned (B), of the LINC00473 height association observed in the trans-ethnic meta-analysis of Hispanic/Latino Anthropometry Consortium (HISLA, including stage 1), the African Ancestry Anthropometry Genetics Consortium (AAAGC) and/or the Genetic Investigation of Anthropometric Traits (GIANT) Consortium. Linkage disequilibrium patterns are based on rs991946 (shown by the purple triangle) and reflect the European, African, and Hispanic ancestral populations of the GWAS results.

### Slide 31
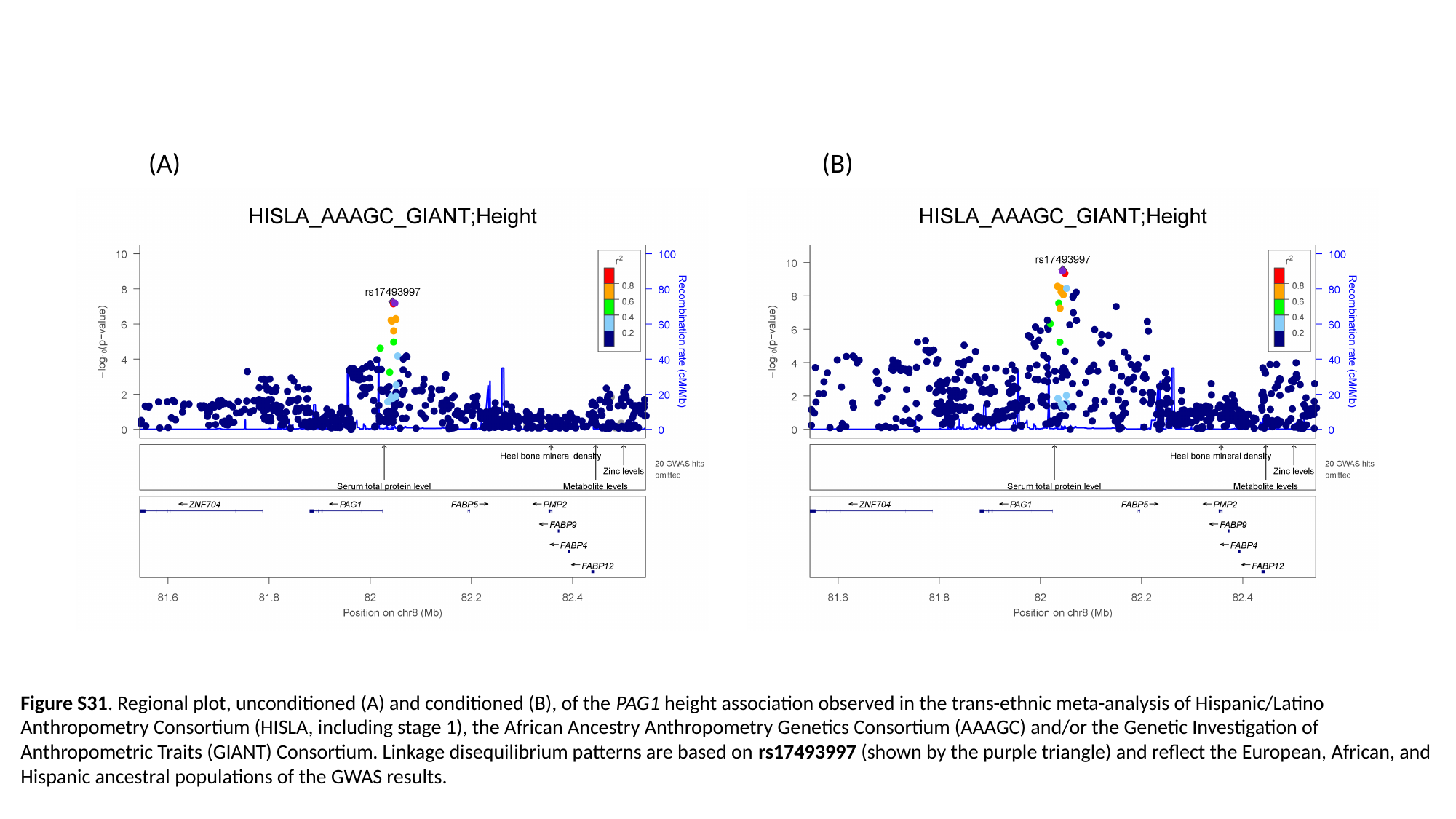

(A) (B)
Figure S31. Regional plot, unconditioned (A) and conditioned (B), of the PAG1 height association observed in the trans-ethnic meta-analysis of Hispanic/Latino Anthropometry Consortium (HISLA, including stage 1), the African Ancestry Anthropometry Genetics Consortium (AAAGC) and/or the Genetic Investigation of Anthropometric Traits (GIANT) Consortium. Linkage disequilibrium patterns are based on rs17493997 (shown by the purple triangle) and reflect the European, African, and Hispanic ancestral populations of the GWAS results.

### Slide 32
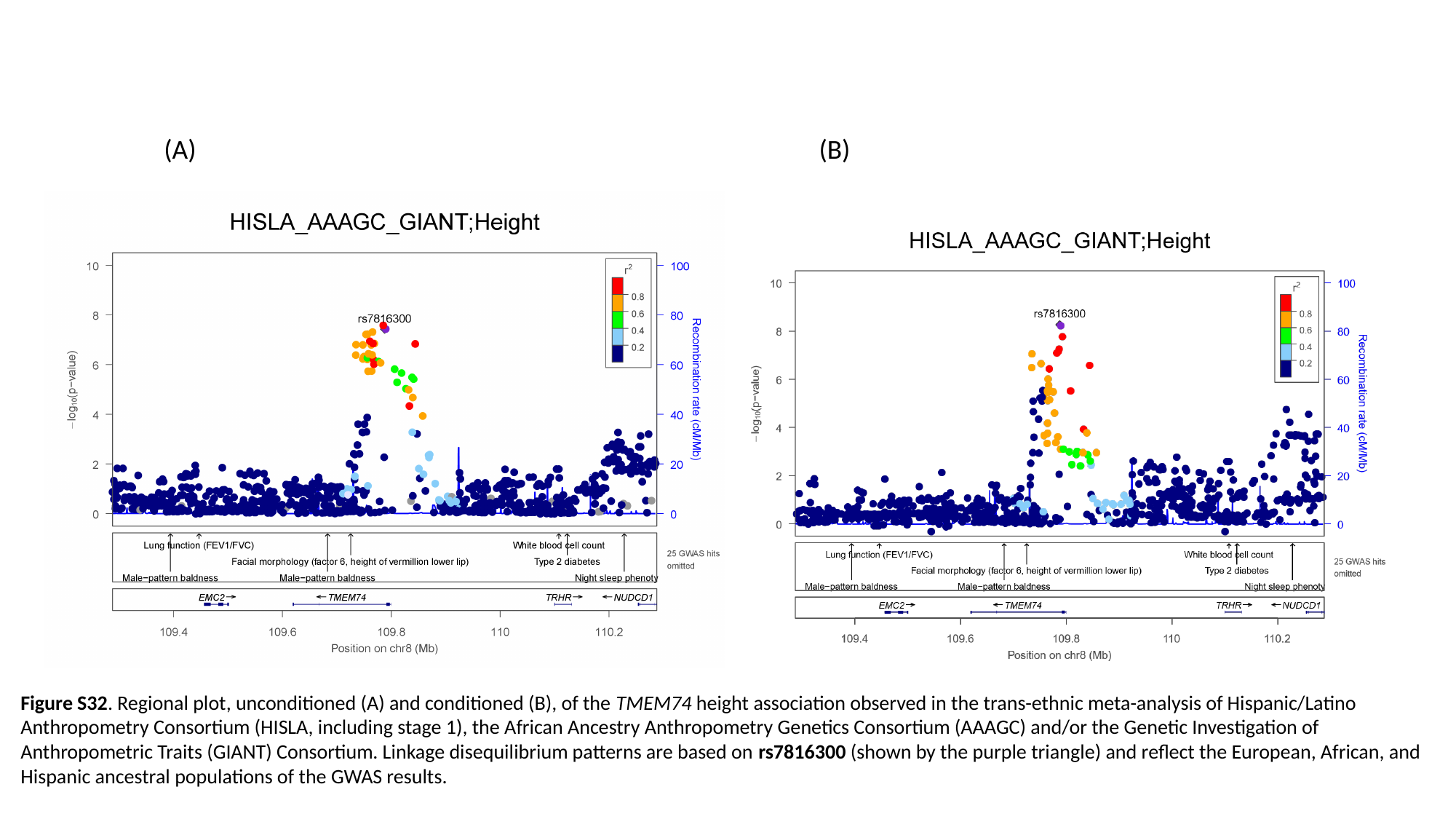

(A) (B)
Figure S32. Regional plot, unconditioned (A) and conditioned (B), of the TMEM74 height association observed in the trans-ethnic meta-analysis of Hispanic/Latino Anthropometry Consortium (HISLA, including stage 1), the African Ancestry Anthropometry Genetics Consortium (AAAGC) and/or the Genetic Investigation of Anthropometric Traits (GIANT) Consortium. Linkage disequilibrium patterns are based on rs7816300 (shown by the purple triangle) and reflect the European, African, and Hispanic ancestral populations of the GWAS results.

### Slide 33
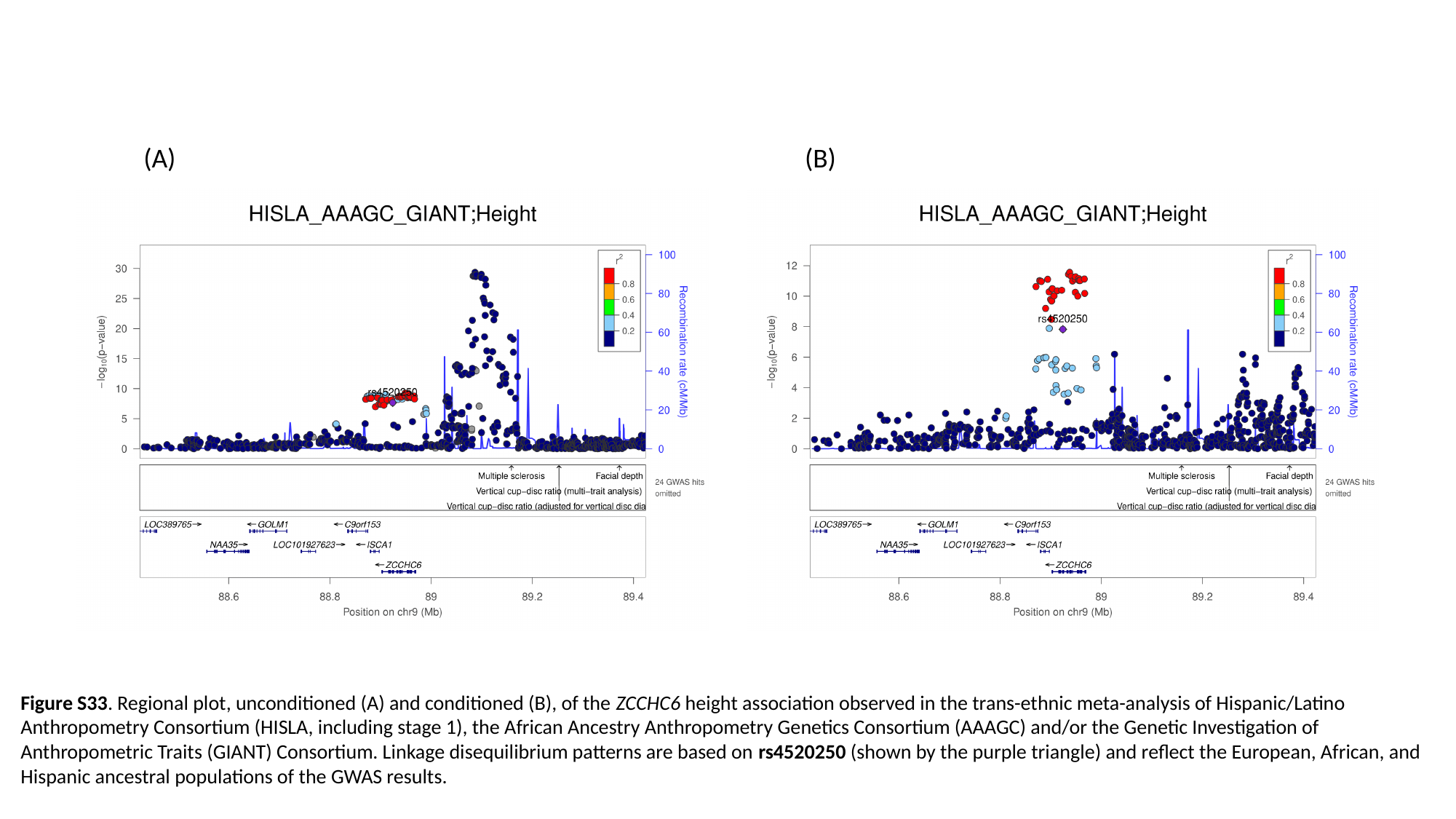

(A) (B)
Figure S33. Regional plot, unconditioned (A) and conditioned (B), of the ZCCHC6 height association observed in the trans-ethnic meta-analysis of Hispanic/Latino Anthropometry Consortium (HISLA, including stage 1), the African Ancestry Anthropometry Genetics Consortium (AAAGC) and/or the Genetic Investigation of Anthropometric Traits (GIANT) Consortium. Linkage disequilibrium patterns are based on rs4520250 (shown by the purple triangle) and reflect the European, African, and Hispanic ancestral populations of the GWAS results.

### Slide 34
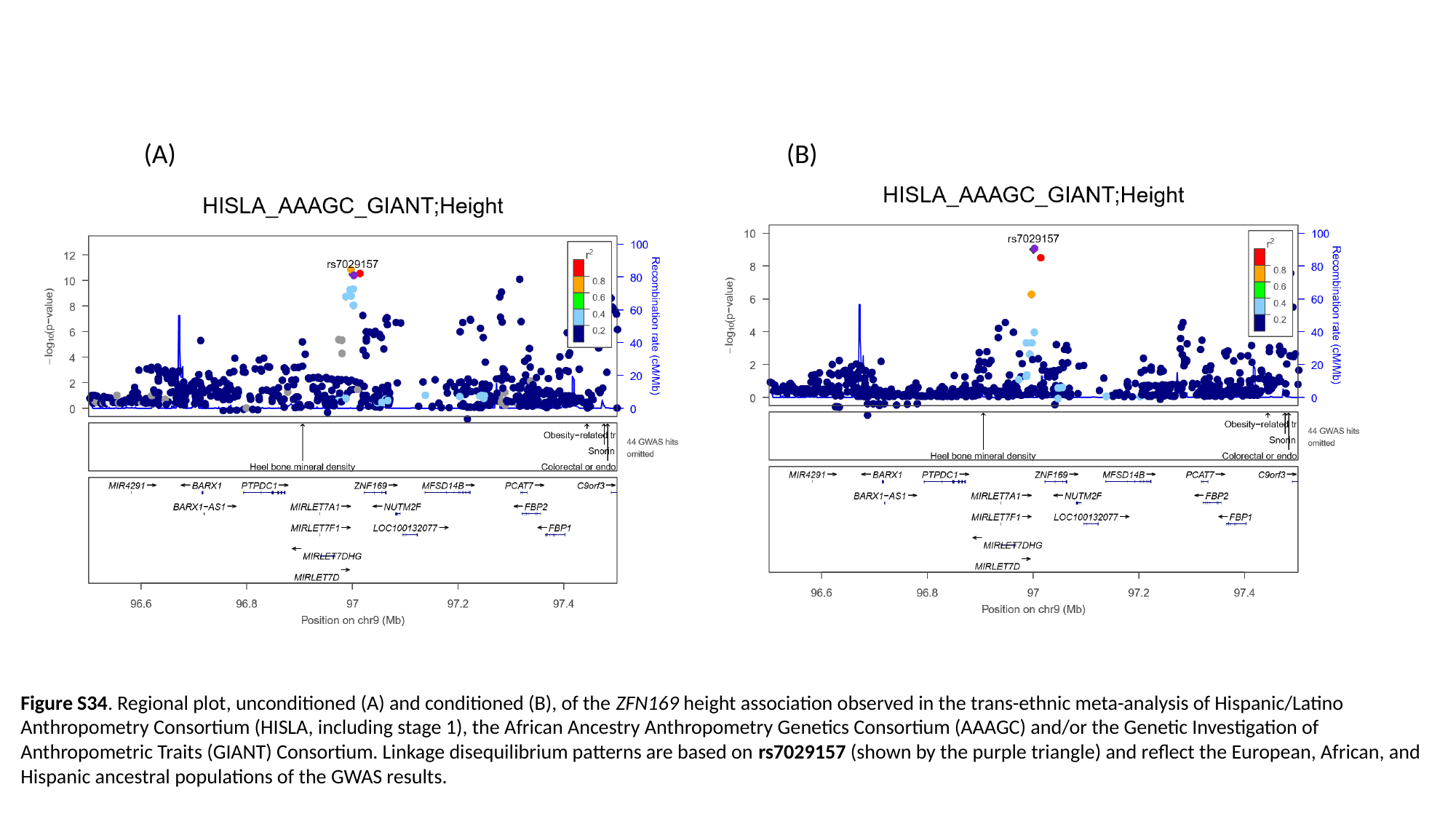

(A) (B)
Figure S34. Regional plot, unconditioned (A) and conditioned (B), of the ZFN169 height association observed in the trans-ethnic meta-analysis of Hispanic/Latino Anthropometry Consortium (HISLA, including stage 1), the African Ancestry Anthropometry Genetics Consortium (AAAGC) and/or the Genetic Investigation of Anthropometric Traits (GIANT) Consortium. Linkage disequilibrium patterns are based on rs7029157 (shown by the purple triangle) and reflect the European, African, and Hispanic ancestral populations of the GWAS results.

### Slide 35
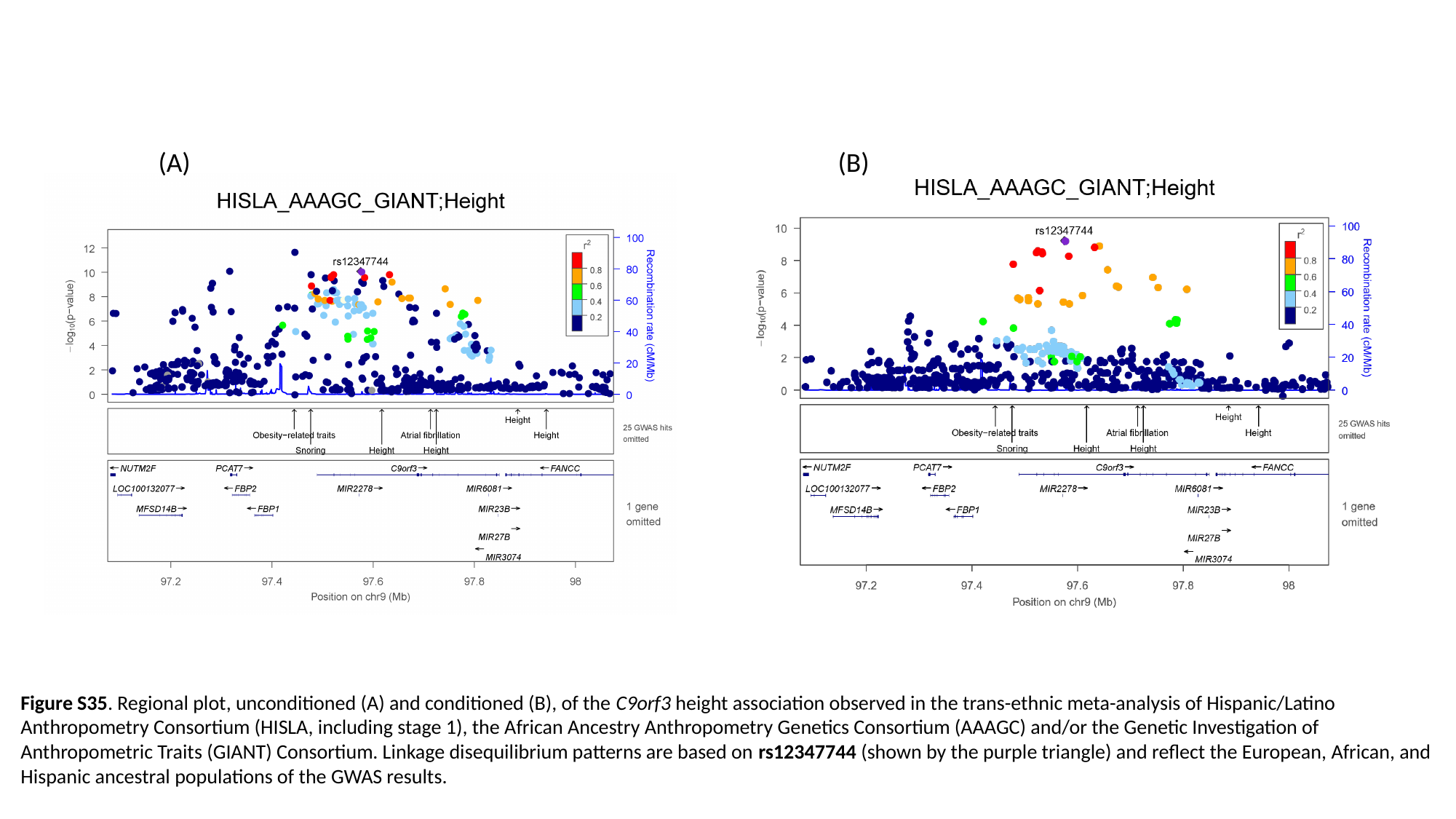

(A) (B)
Figure S35. Regional plot, unconditioned (A) and conditioned (B), of the C9orf3 height association observed in the trans-ethnic meta-analysis of Hispanic/Latino Anthropometry Consortium (HISLA, including stage 1), the African Ancestry Anthropometry Genetics Consortium (AAAGC) and/or the Genetic Investigation of Anthropometric Traits (GIANT) Consortium. Linkage disequilibrium patterns are based on rs12347744 (shown by the purple triangle) and reflect the European, African, and Hispanic ancestral populations of the GWAS results.

### Slide 36
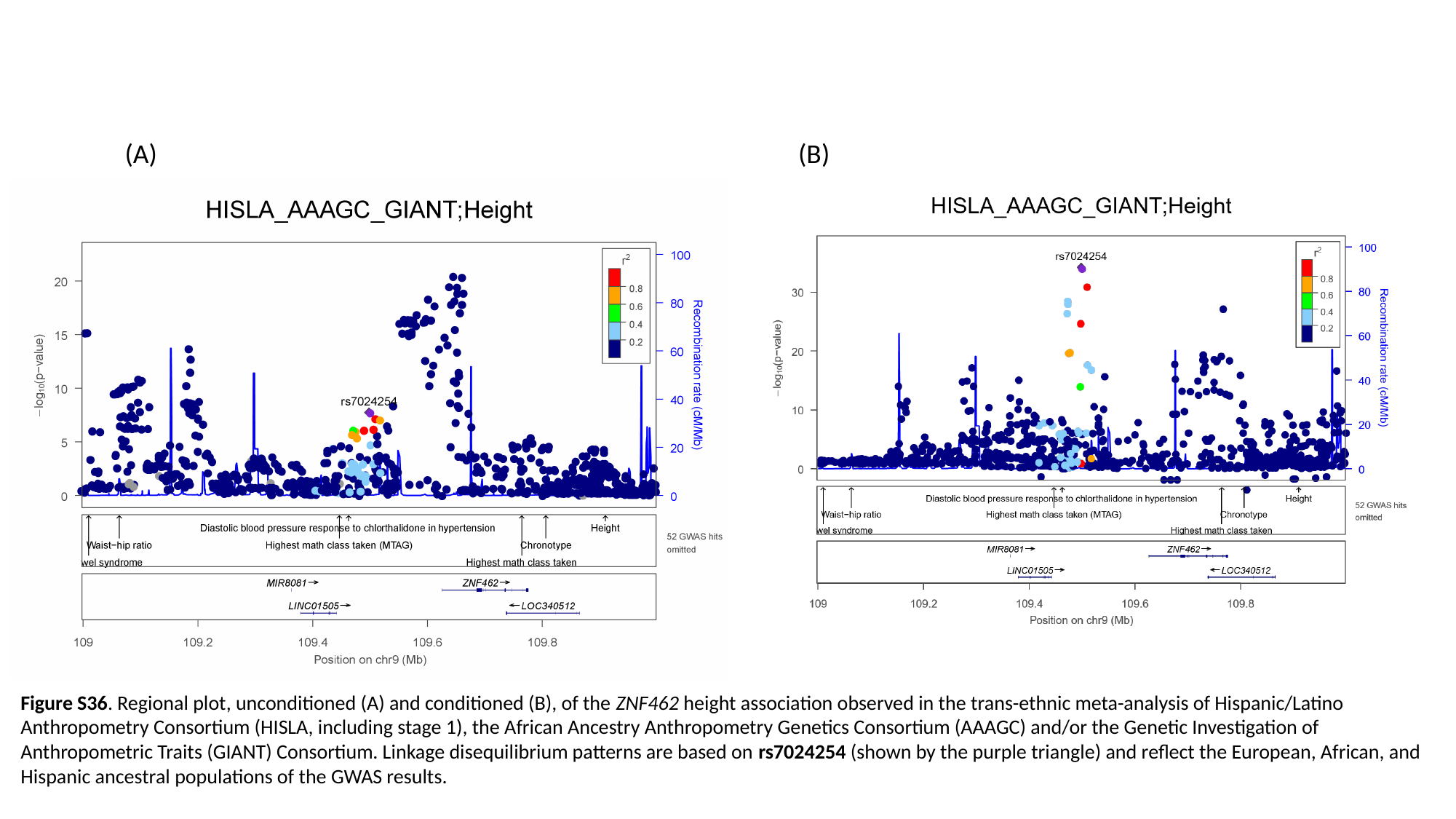

(A) (B)
Figure S36. Regional plot, unconditioned (A) and conditioned (B), of the ZNF462 height association observed in the trans-ethnic meta-analysis of Hispanic/Latino Anthropometry Consortium (HISLA, including stage 1), the African Ancestry Anthropometry Genetics Consortium (AAAGC) and/or the Genetic Investigation of Anthropometric Traits (GIANT) Consortium. Linkage disequilibrium patterns are based on rs7024254 (shown by the purple triangle) and reflect the European, African, and Hispanic ancestral populations of the GWAS results.

### Slide 37
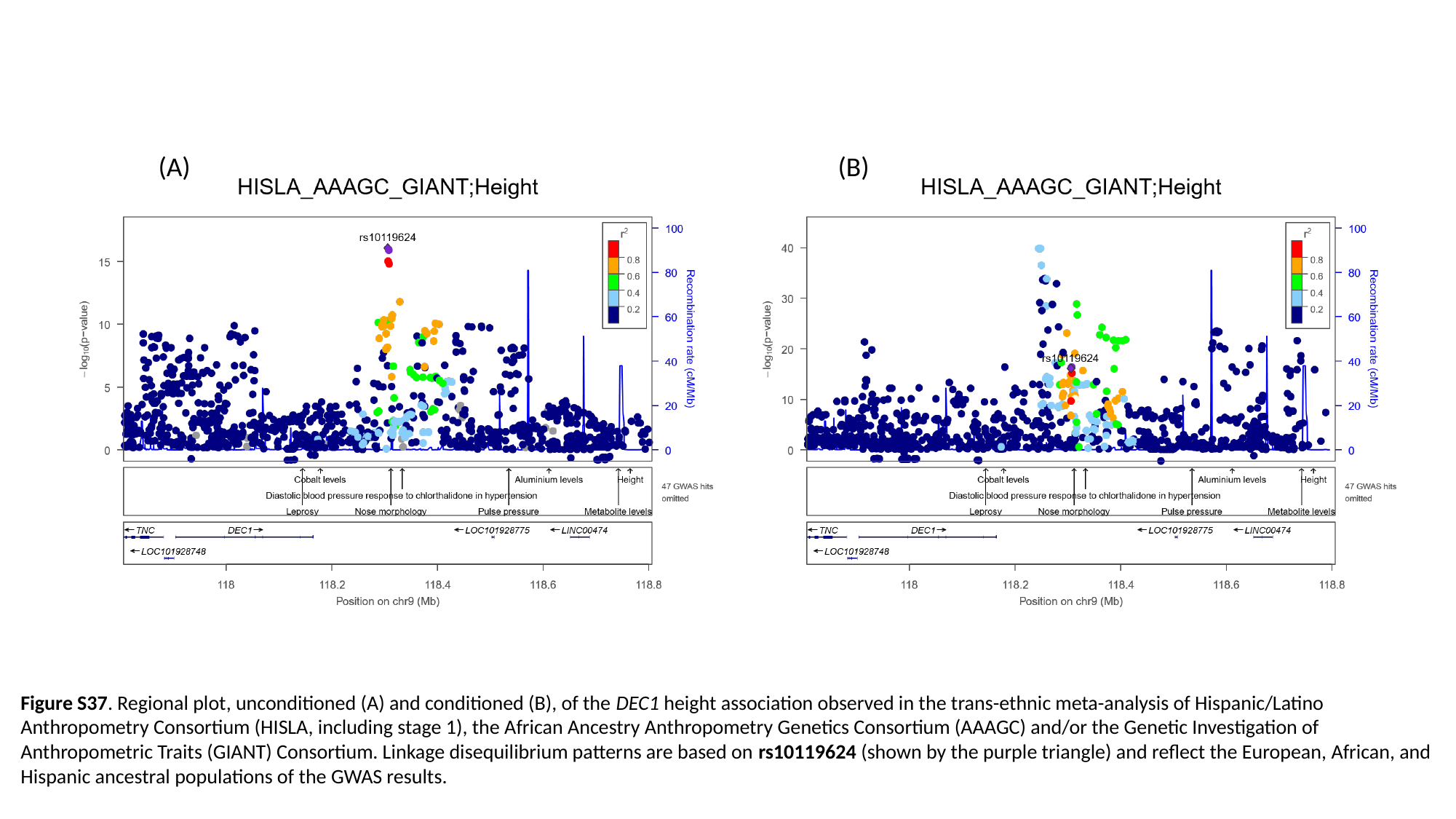

(A) (B)
Figure S37. Regional plot, unconditioned (A) and conditioned (B), of the DEC1 height association observed in the trans-ethnic meta-analysis of Hispanic/Latino Anthropometry Consortium (HISLA, including stage 1), the African Ancestry Anthropometry Genetics Consortium (AAAGC) and/or the Genetic Investigation of Anthropometric Traits (GIANT) Consortium. Linkage disequilibrium patterns are based on rs10119624 (shown by the purple triangle) and reflect the European, African, and Hispanic ancestral populations of the GWAS results.

### Slide 38
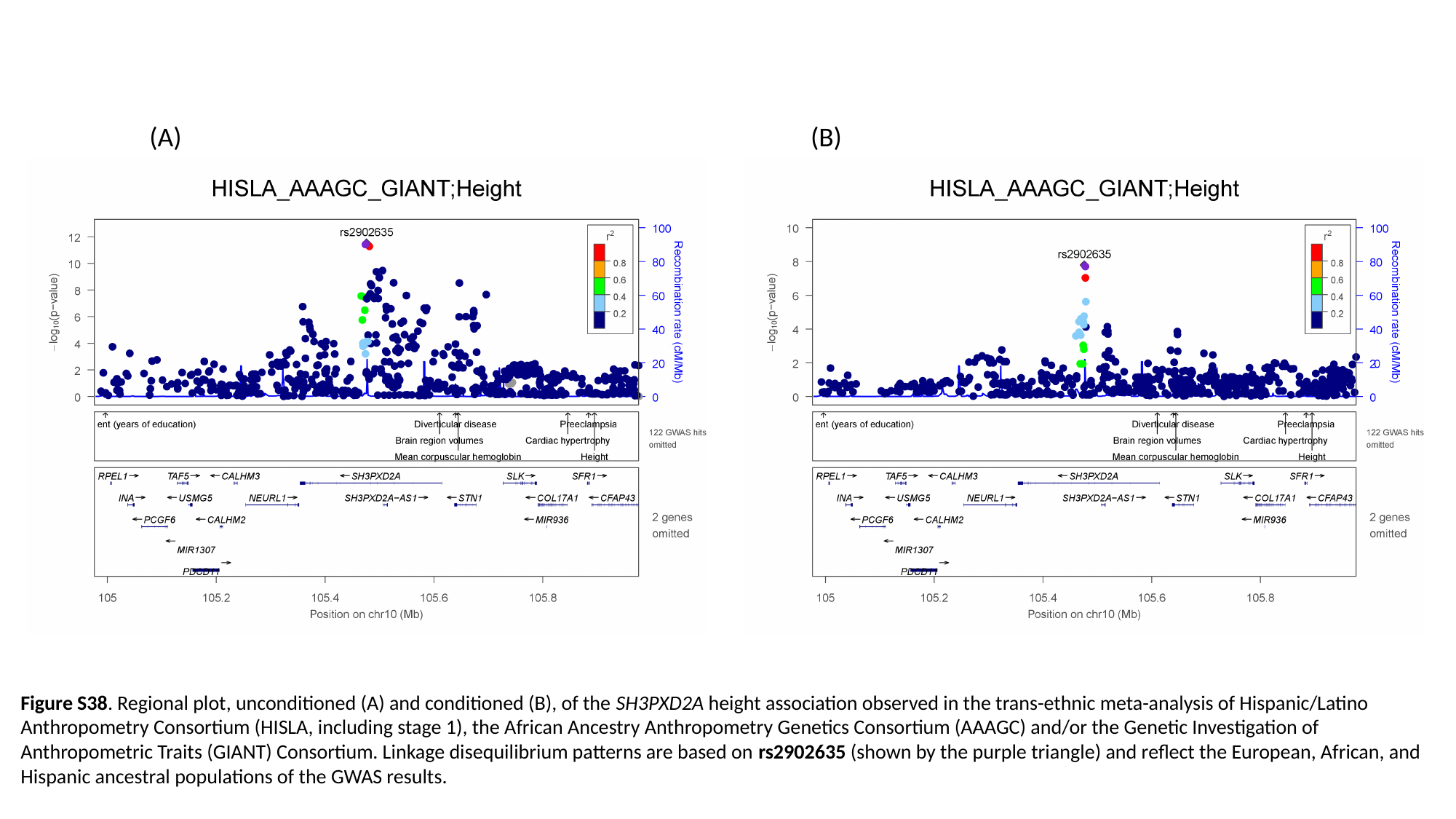

(A) (B)
Figure S38. Regional plot, unconditioned (A) and conditioned (B), of the SH3PXD2A height association observed in the trans-ethnic meta-analysis of Hispanic/Latino Anthropometry Consortium (HISLA, including stage 1), the African Ancestry Anthropometry Genetics Consortium (AAAGC) and/or the Genetic Investigation of Anthropometric Traits (GIANT) Consortium. Linkage disequilibrium patterns are based on rs2902635 (shown by the purple triangle) and reflect the European, African, and Hispanic ancestral populations of the GWAS results.

### Slide 39
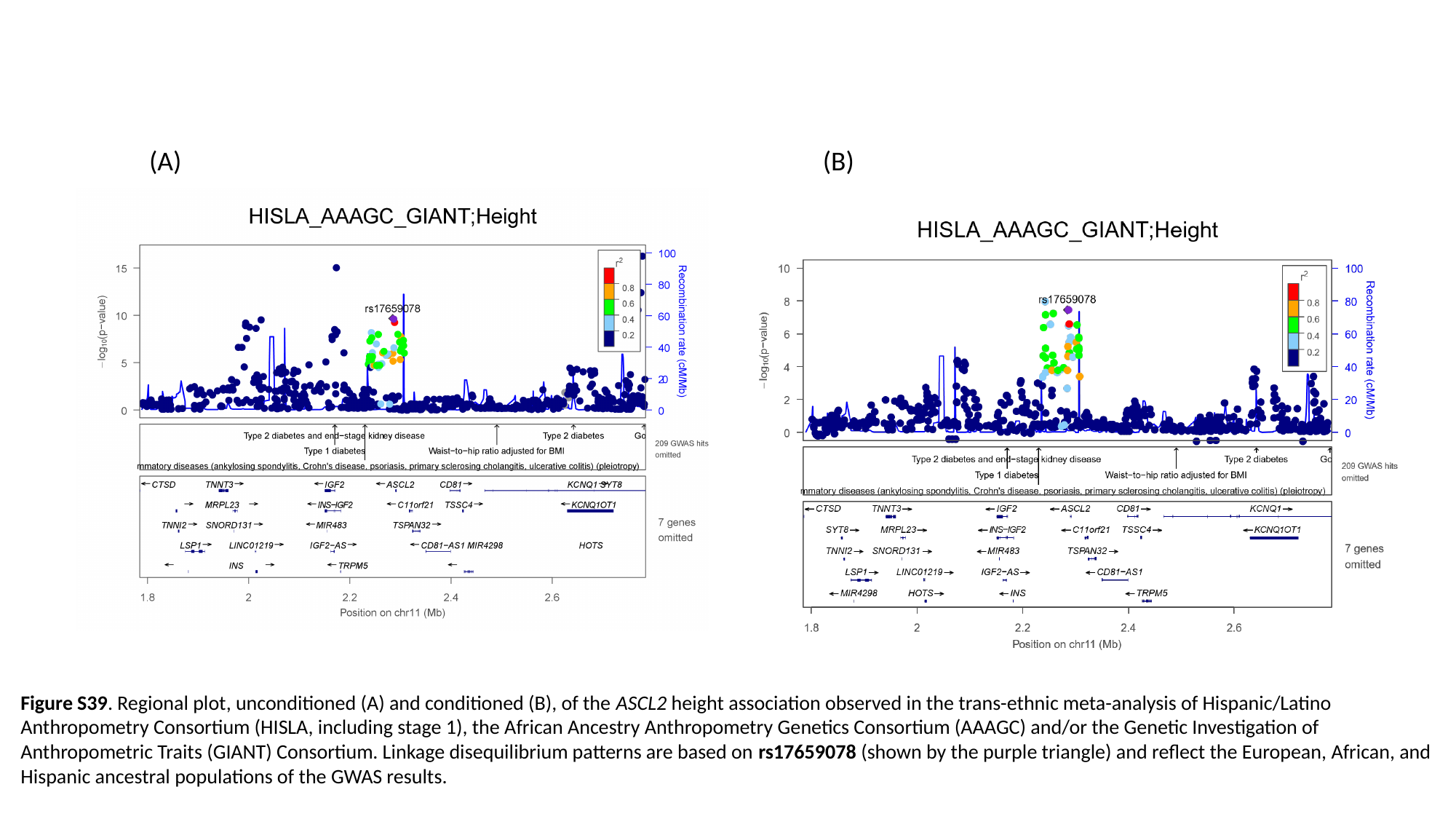

(A) (B)
Figure S39. Regional plot, unconditioned (A) and conditioned (B), of the ASCL2 height association observed in the trans-ethnic meta-analysis of Hispanic/Latino Anthropometry Consortium (HISLA, including stage 1), the African Ancestry Anthropometry Genetics Consortium (AAAGC) and/or the Genetic Investigation of Anthropometric Traits (GIANT) Consortium. Linkage disequilibrium patterns are based on rs17659078 (shown by the purple triangle) and reflect the European, African, and Hispanic ancestral populations of the GWAS results.

### Slide 40
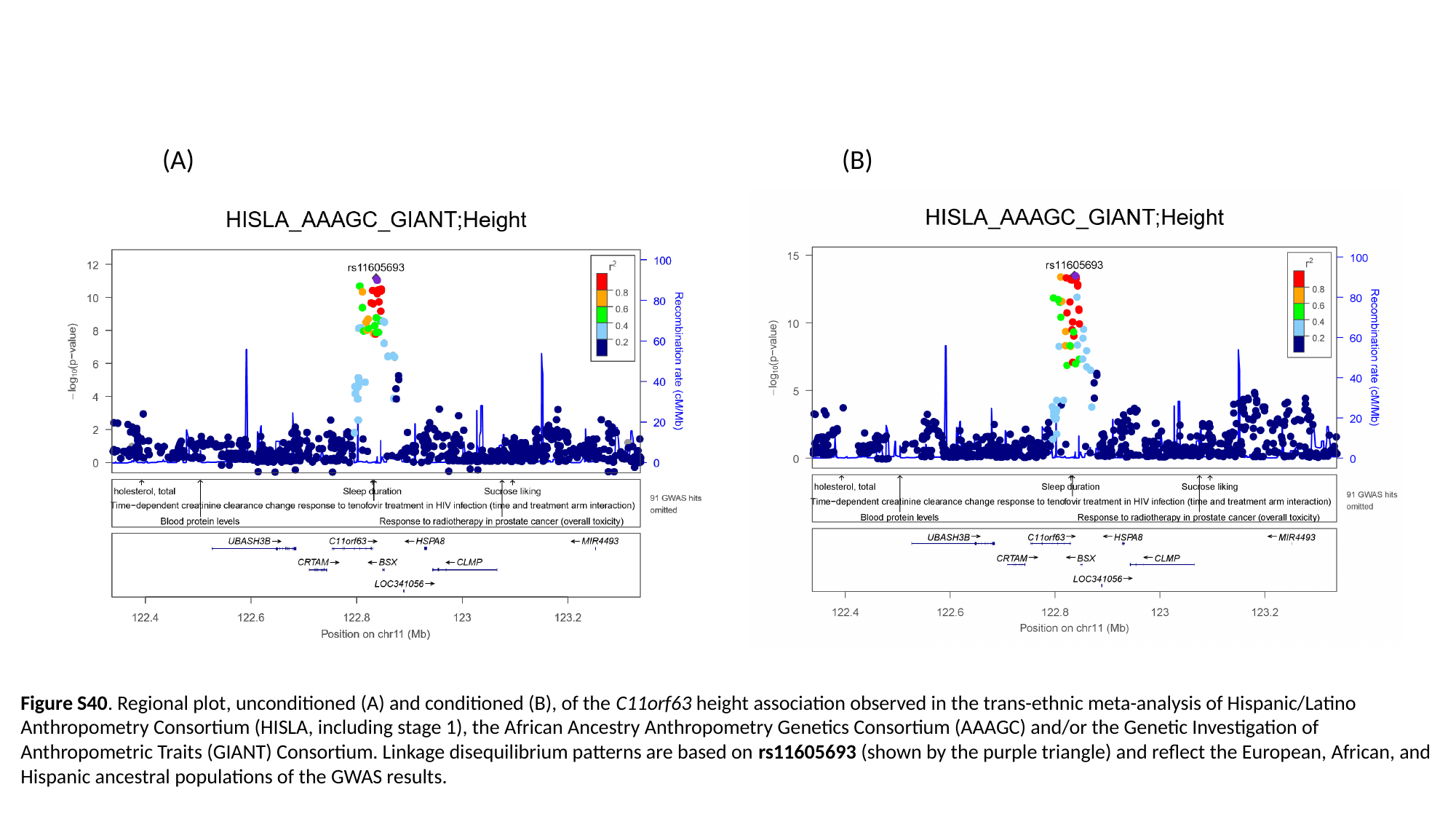

(A) (B)
Figure S40. Regional plot, unconditioned (A) and conditioned (B), of the C11orf63 height association observed in the trans-ethnic meta-analysis of Hispanic/Latino Anthropometry Consortium (HISLA, including stage 1), the African Ancestry Anthropometry Genetics Consortium (AAAGC) and/or the Genetic Investigation of Anthropometric Traits (GIANT) Consortium. Linkage disequilibrium patterns are based on rs11605693 (shown by the purple triangle) and reflect the European, African, and Hispanic ancestral populations of the GWAS results.

### Slide 41
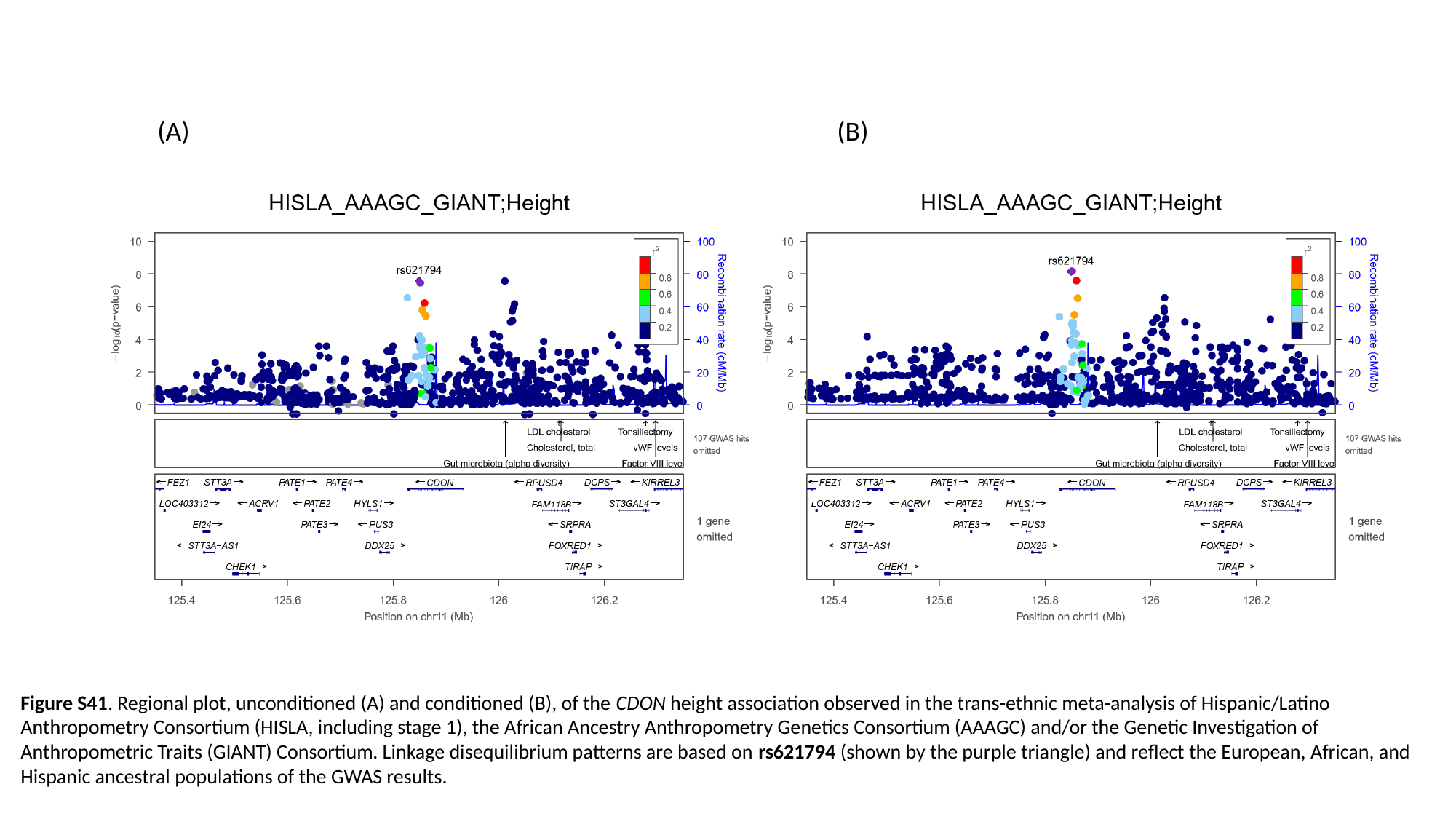

(A) (B)
Figure S41. Regional plot, unconditioned (A) and conditioned (B), of the CDON height association observed in the trans-ethnic meta-analysis of Hispanic/Latino Anthropometry Consortium (HISLA, including stage 1), the African Ancestry Anthropometry Genetics Consortium (AAAGC) and/or the Genetic Investigation of Anthropometric Traits (GIANT) Consortium. Linkage disequilibrium patterns are based on rs621794 (shown by the purple triangle) and reflect the European, African, and Hispanic ancestral populations of the GWAS results.

### Slide 42
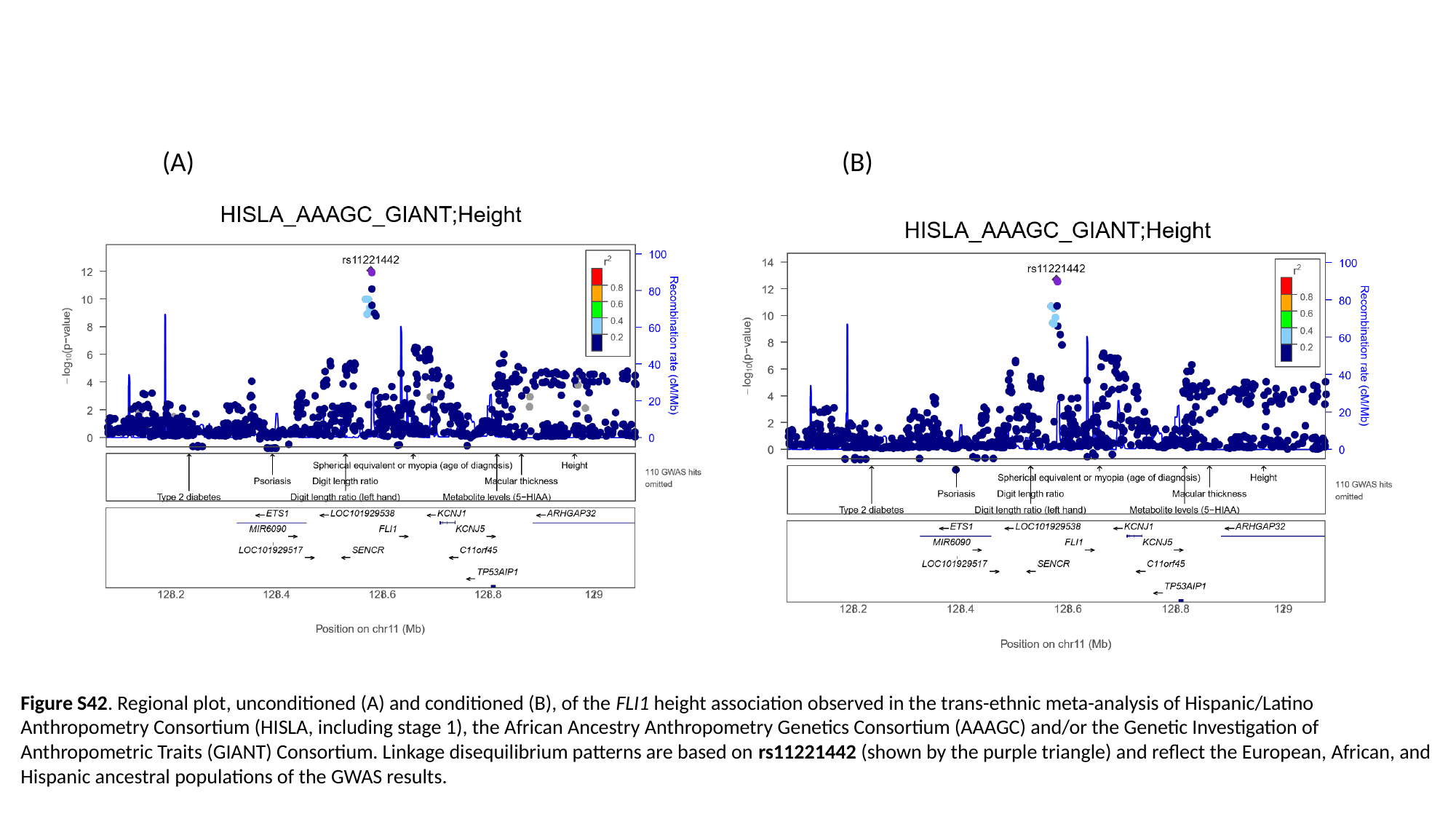

(A) (B)
Figure S42. Regional plot, unconditioned (A) and conditioned (B), of the FLI1 height association observed in the trans-ethnic meta-analysis of Hispanic/Latino Anthropometry Consortium (HISLA, including stage 1), the African Ancestry Anthropometry Genetics Consortium (AAAGC) and/or the Genetic Investigation of Anthropometric Traits (GIANT) Consortium. Linkage disequilibrium patterns are based on rs11221442 (shown by the purple triangle) and reflect the European, African, and Hispanic ancestral populations of the GWAS results.

### Slide 43
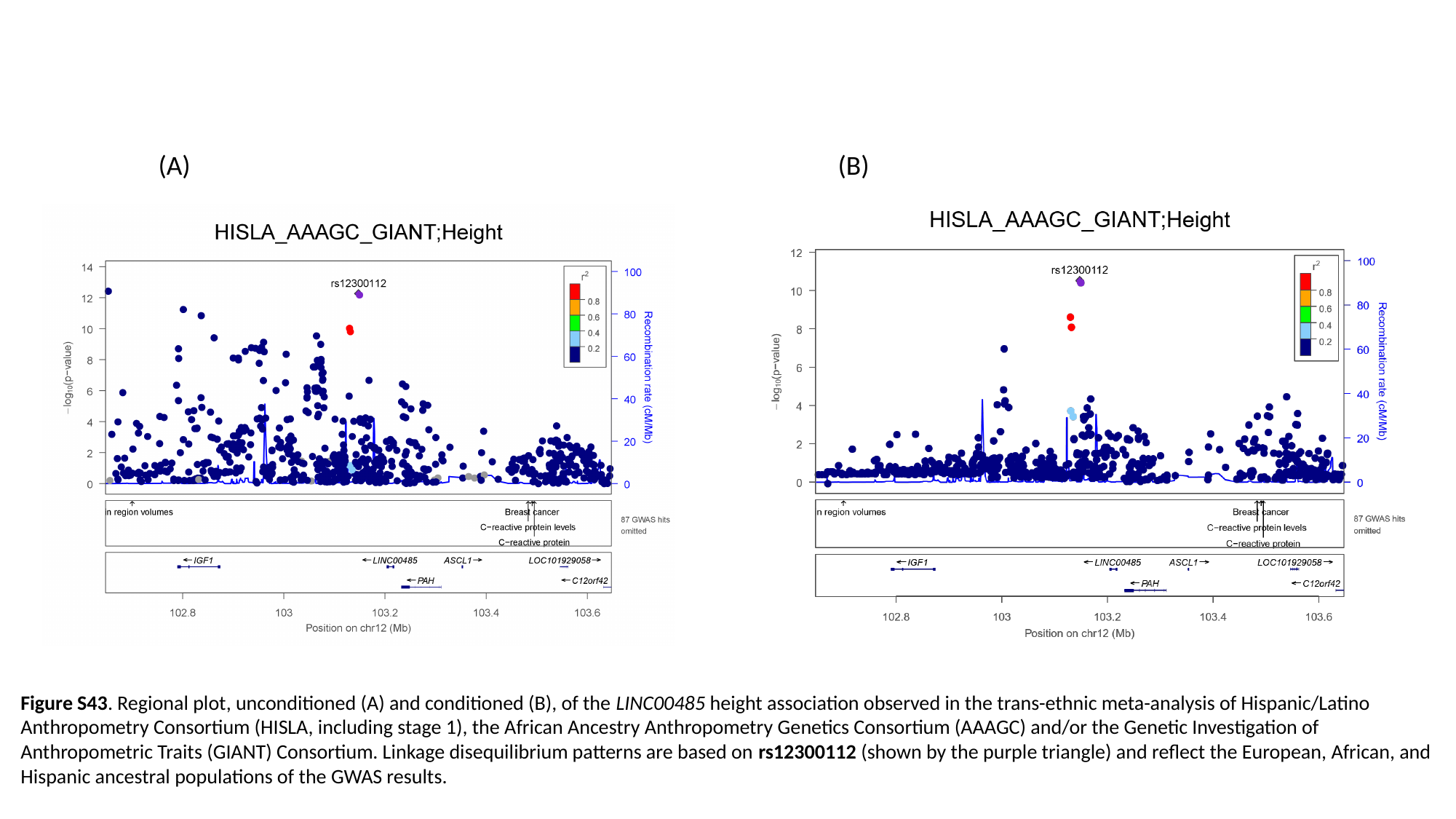

(A) (B)
Figure S43. Regional plot, unconditioned (A) and conditioned (B), of the LINC00485 height association observed in the trans-ethnic meta-analysis of Hispanic/Latino Anthropometry Consortium (HISLA, including stage 1), the African Ancestry Anthropometry Genetics Consortium (AAAGC) and/or the Genetic Investigation of Anthropometric Traits (GIANT) Consortium. Linkage disequilibrium patterns are based on rs12300112 (shown by the purple triangle) and reflect the European, African, and Hispanic ancestral populations of the GWAS results.

### Slide 44
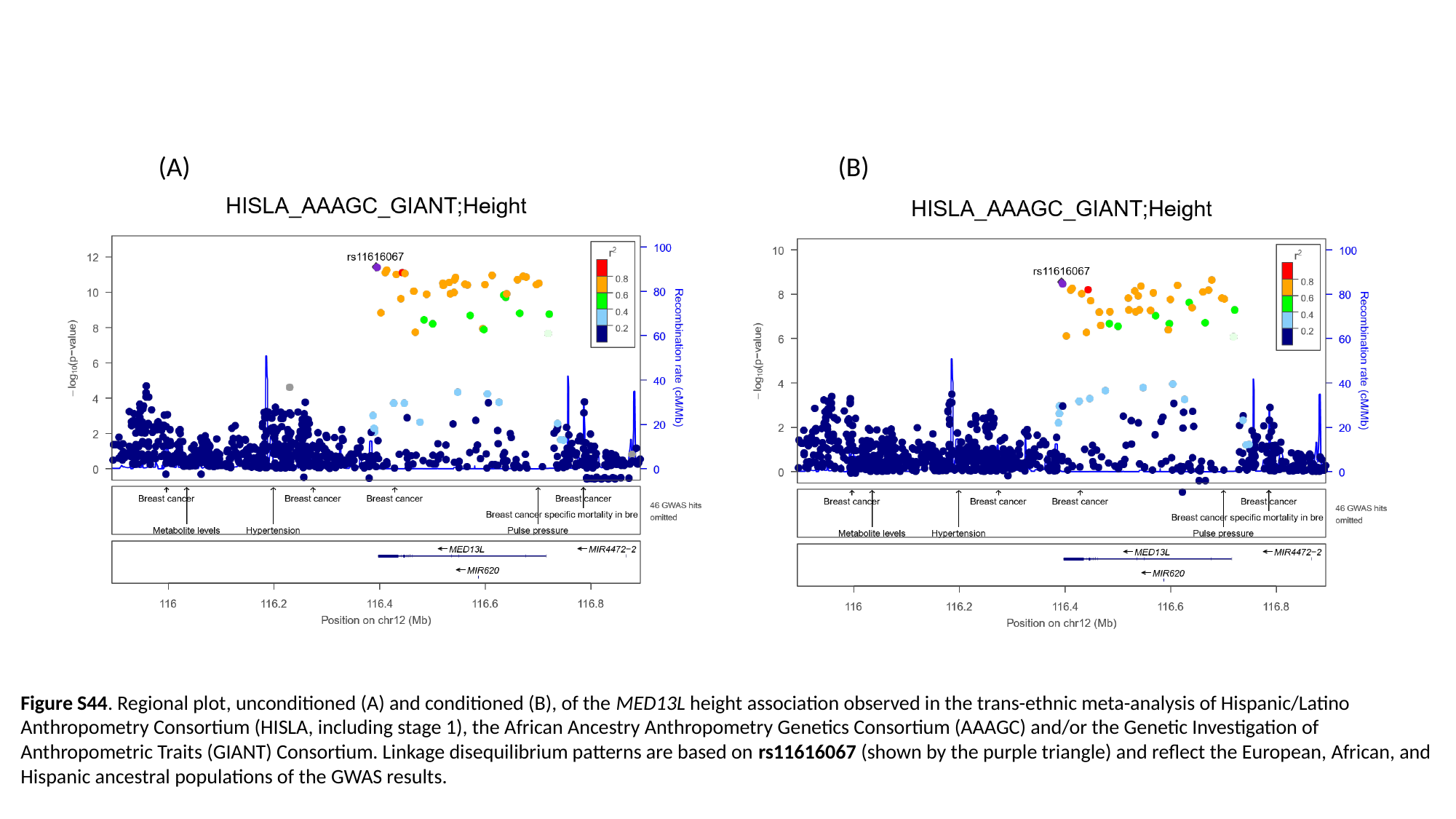

(A) (B)
Figure S44. Regional plot, unconditioned (A) and conditioned (B), of the MED13L height association observed in the trans-ethnic meta-analysis of Hispanic/Latino Anthropometry Consortium (HISLA, including stage 1), the African Ancestry Anthropometry Genetics Consortium (AAAGC) and/or the Genetic Investigation of Anthropometric Traits (GIANT) Consortium. Linkage disequilibrium patterns are based on rs11616067 (shown by the purple triangle) and reflect the European, African, and Hispanic ancestral populations of the GWAS results.

### Slide 45
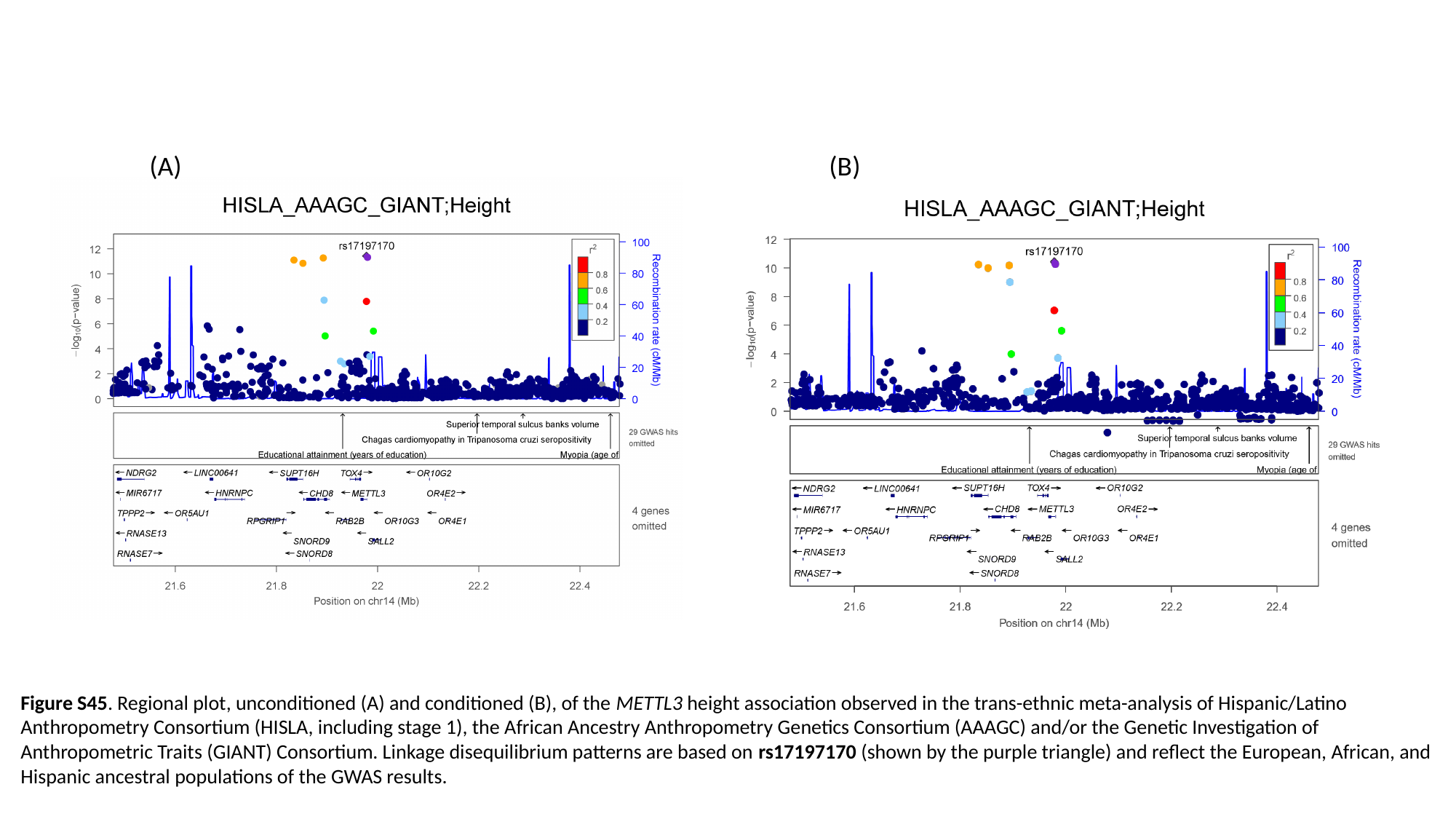

(A) (B)
Figure S45. Regional plot, unconditioned (A) and conditioned (B), of the METTL3 height association observed in the trans-ethnic meta-analysis of Hispanic/Latino Anthropometry Consortium (HISLA, including stage 1), the African Ancestry Anthropometry Genetics Consortium (AAAGC) and/or the Genetic Investigation of Anthropometric Traits (GIANT) Consortium. Linkage disequilibrium patterns are based on rs17197170 (shown by the purple triangle) and reflect the European, African, and Hispanic ancestral populations of the GWAS results.

### Slide 46
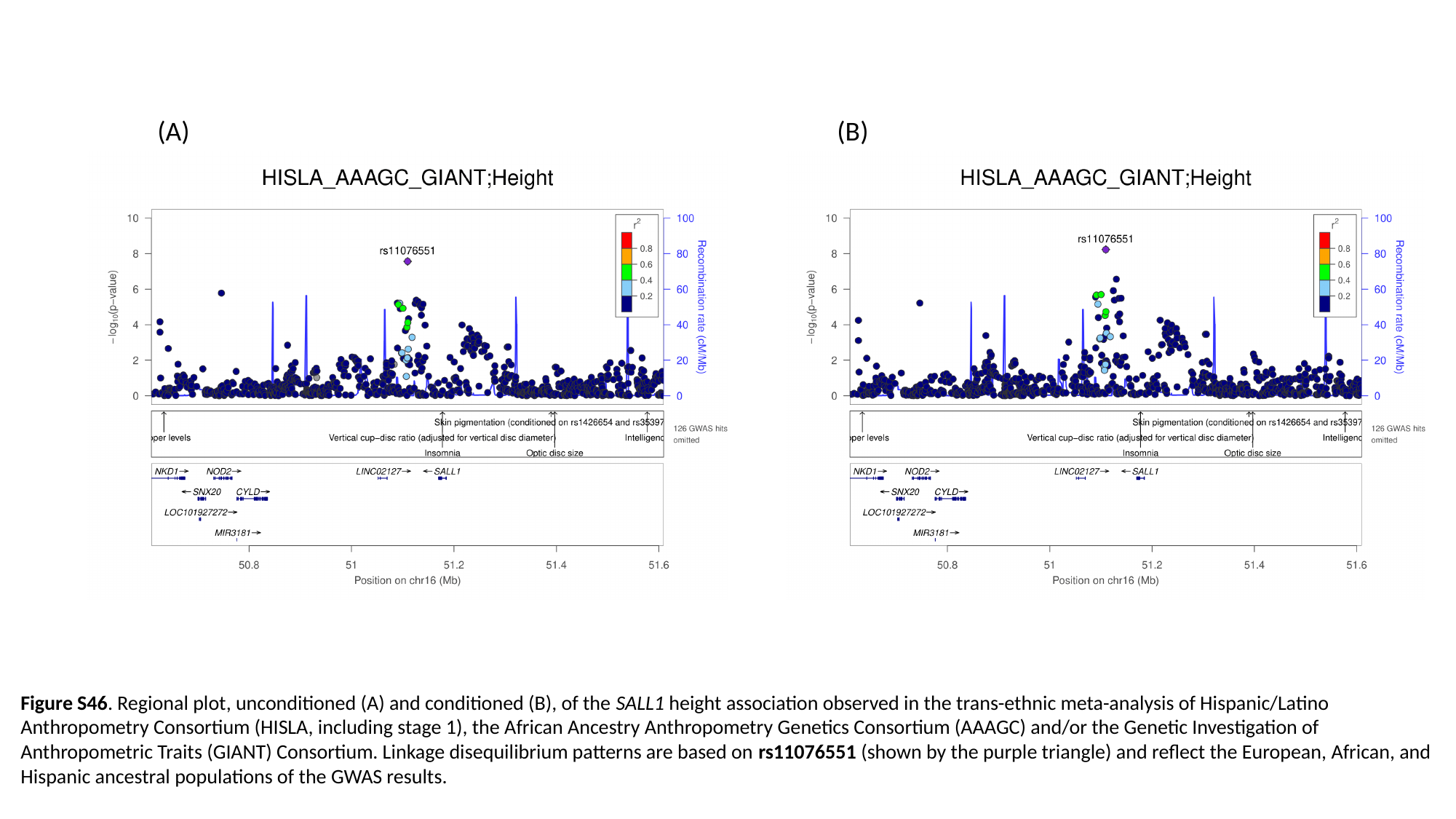

(A) (B)
Figure S46. Regional plot, unconditioned (A) and conditioned (B), of the SALL1 height association observed in the trans-ethnic meta-analysis of Hispanic/Latino Anthropometry Consortium (HISLA, including stage 1), the African Ancestry Anthropometry Genetics Consortium (AAAGC) and/or the Genetic Investigation of Anthropometric Traits (GIANT) Consortium. Linkage disequilibrium patterns are based on rs11076551 (shown by the purple triangle) and reflect the European, African, and Hispanic ancestral populations of the GWAS results.

### Slide 47
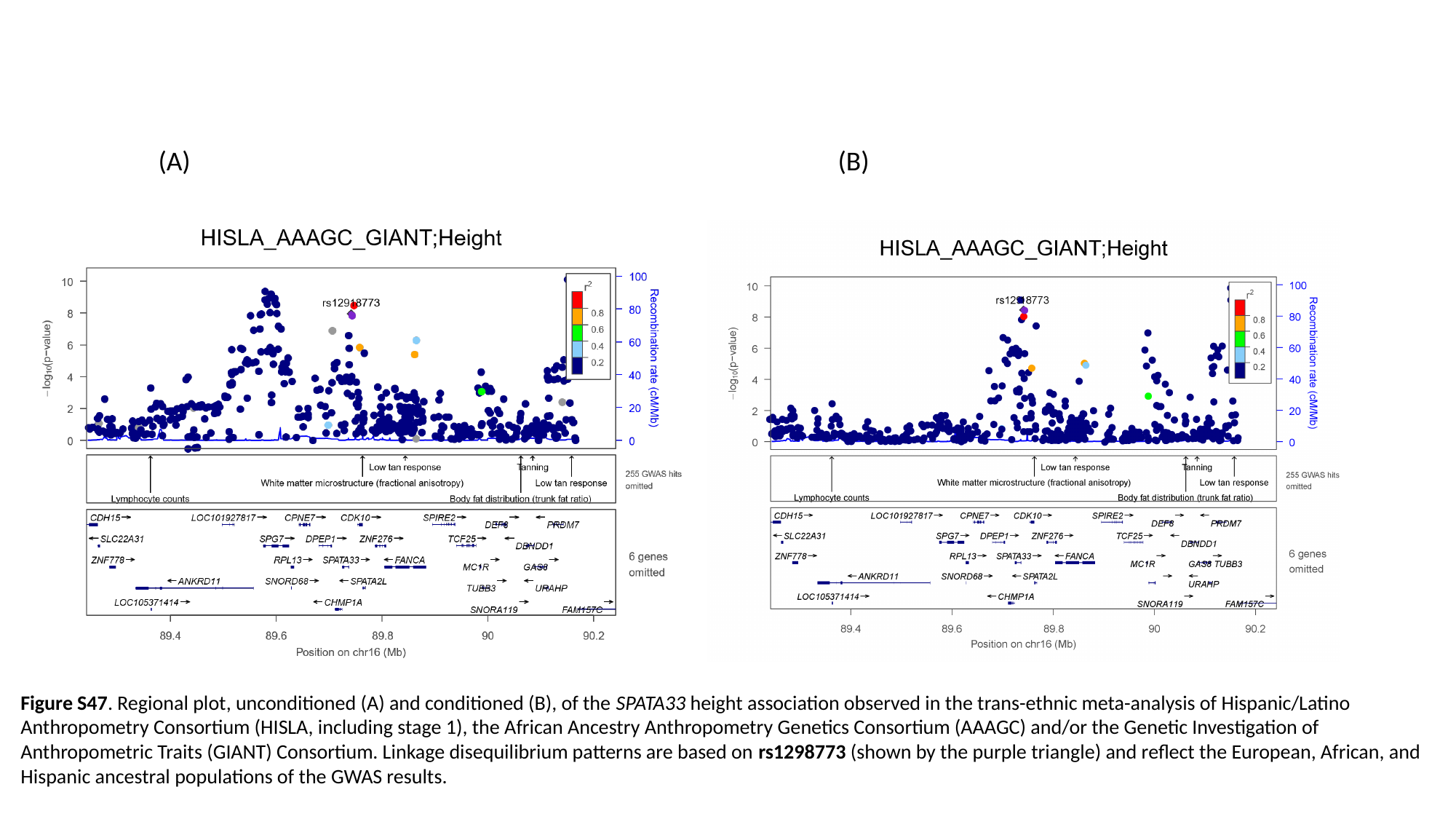

(A) (B)
Figure S47. Regional plot, unconditioned (A) and conditioned (B), of the SPATA33 height association observed in the trans-ethnic meta-analysis of Hispanic/Latino Anthropometry Consortium (HISLA, including stage 1), the African Ancestry Anthropometry Genetics Consortium (AAAGC) and/or the Genetic Investigation of Anthropometric Traits (GIANT) Consortium. Linkage disequilibrium patterns are based on rs1298773 (shown by the purple triangle) and reflect the European, African, and Hispanic ancestral populations of the GWAS results.

### Slide 48
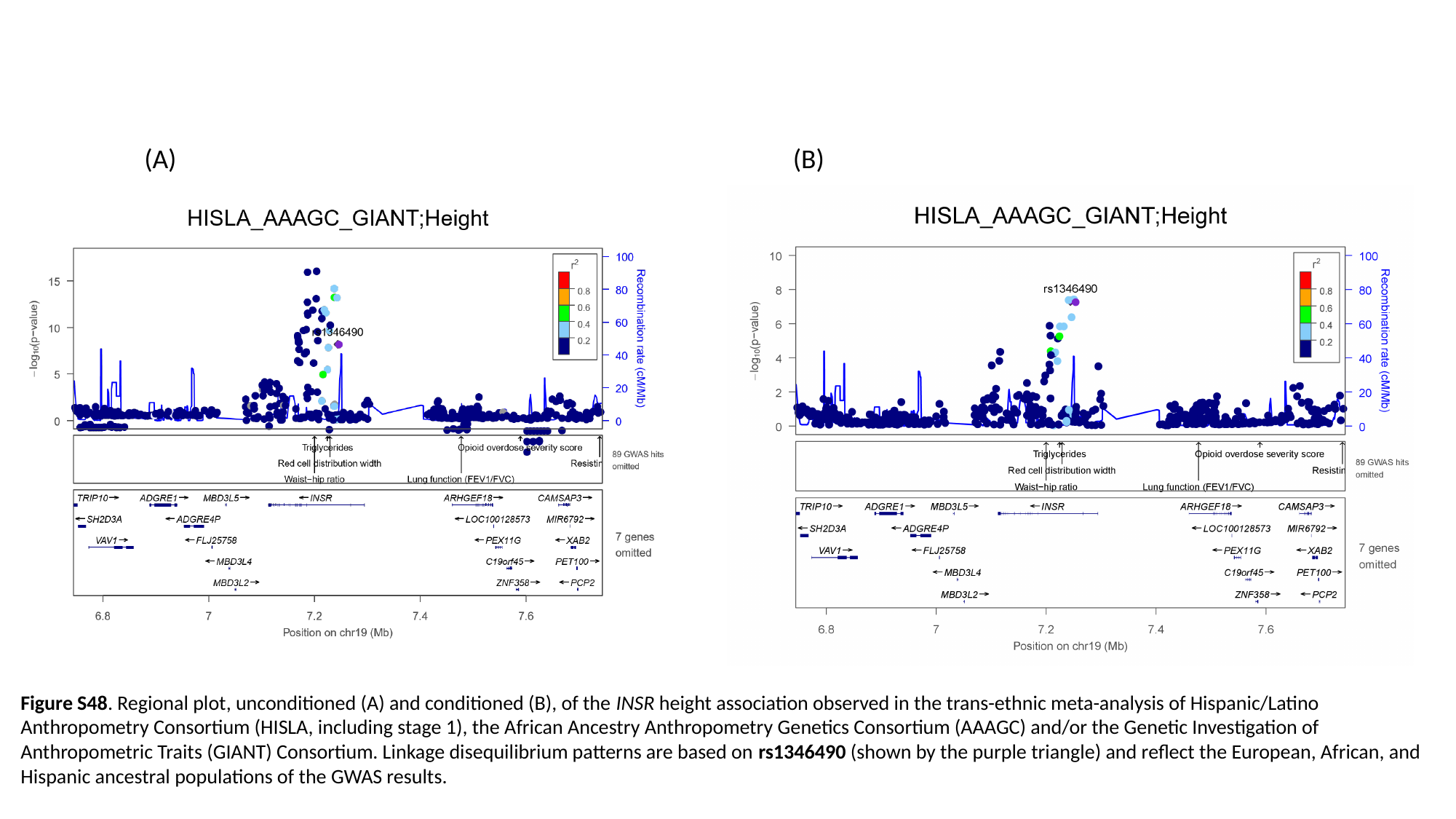

(A) (B)
Figure S48. Regional plot, unconditioned (A) and conditioned (B), of the INSR height association observed in the trans-ethnic meta-analysis of Hispanic/Latino Anthropometry Consortium (HISLA, including stage 1), the African Ancestry Anthropometry Genetics Consortium (AAAGC) and/or the Genetic Investigation of Anthropometric Traits (GIANT) Consortium. Linkage disequilibrium patterns are based on rs1346490 (shown by the purple triangle) and reflect the European, African, and Hispanic ancestral populations of the GWAS results.

### Slide 49
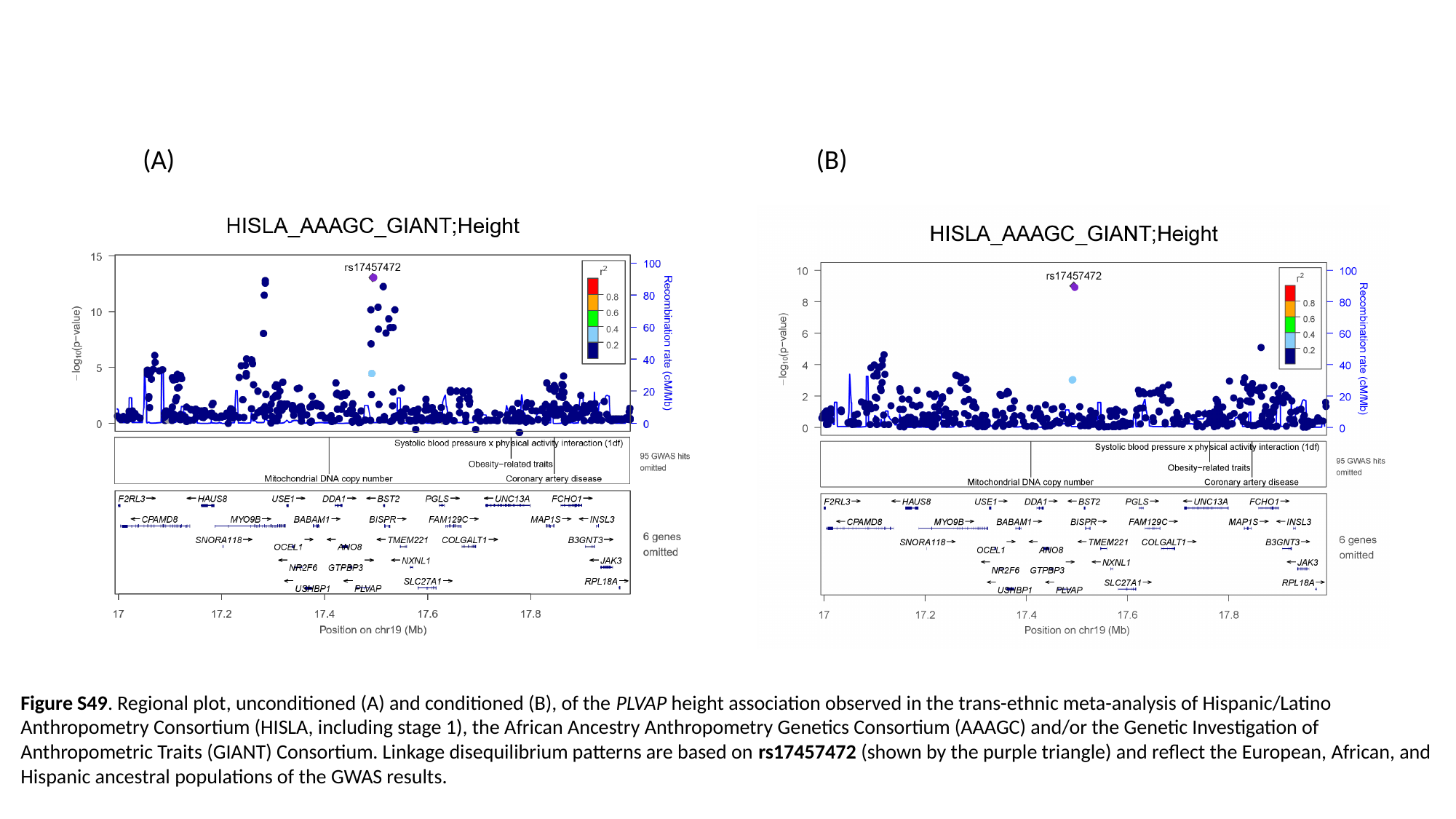

(A) (B)
Figure S49. Regional plot, unconditioned (A) and conditioned (B), of the PLVAP height association observed in the trans-ethnic meta-analysis of Hispanic/Latino Anthropometry Consortium (HISLA, including stage 1), the African Ancestry Anthropometry Genetics Consortium (AAAGC) and/or the Genetic Investigation of Anthropometric Traits (GIANT) Consortium. Linkage disequilibrium patterns are based on rs17457472 (shown by the purple triangle) and reflect the European, African, and Hispanic ancestral populations of the GWAS results.

### Slide 50
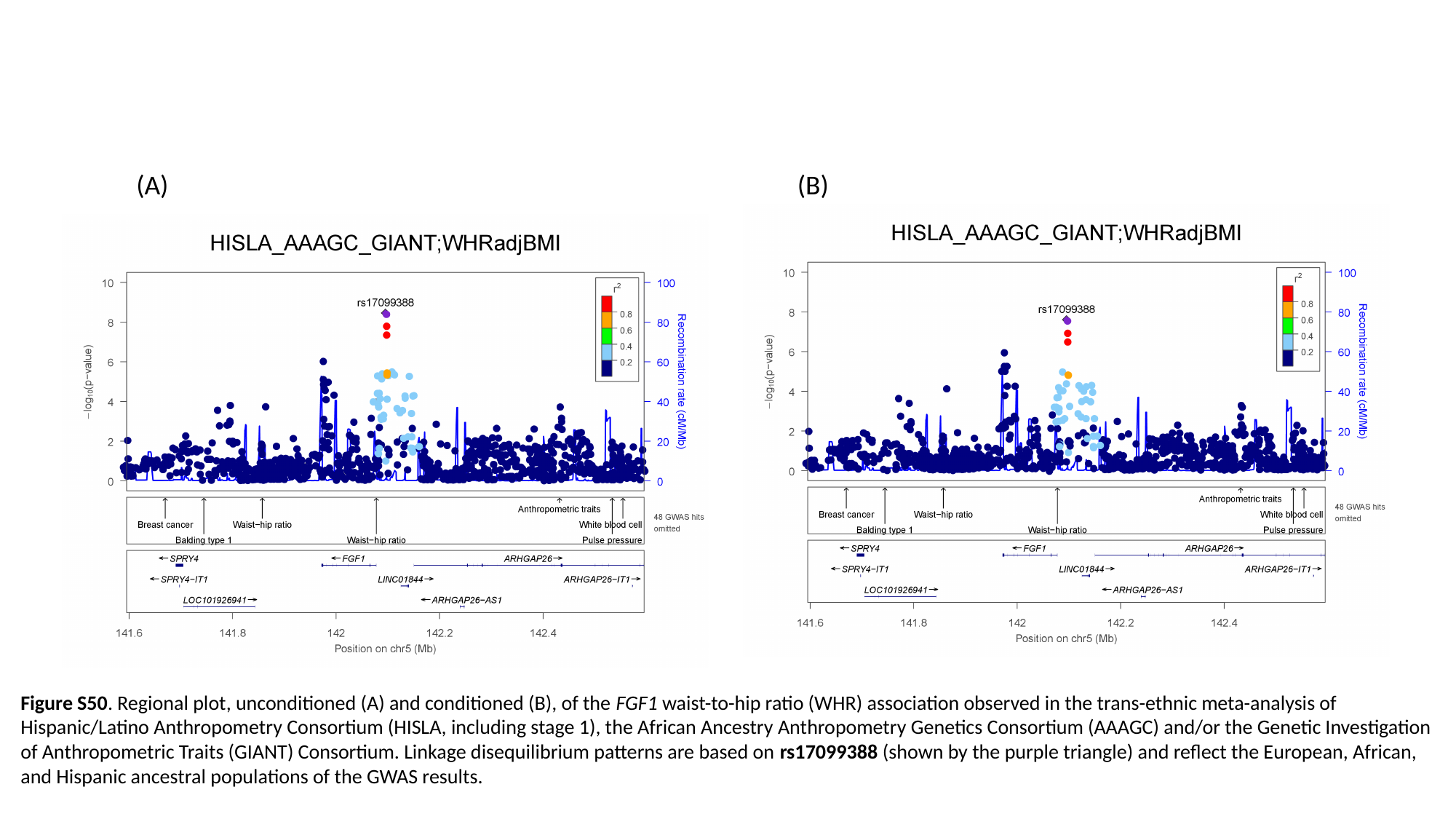

(A) (B)
Figure S50. Regional plot, unconditioned (A) and conditioned (B), of the FGF1 waist-to-hip ratio (WHR) association observed in the trans-ethnic meta-analysis of Hispanic/Latino Anthropometry Consortium (HISLA, including stage 1), the African Ancestry Anthropometry Genetics Consortium (AAAGC) and/or the Genetic Investigation of Anthropometric Traits (GIANT) Consortium. Linkage disequilibrium patterns are based on rs17099388 (shown by the purple triangle) and reflect the European, African, and Hispanic ancestral populations of the GWAS results.

### Slide 51
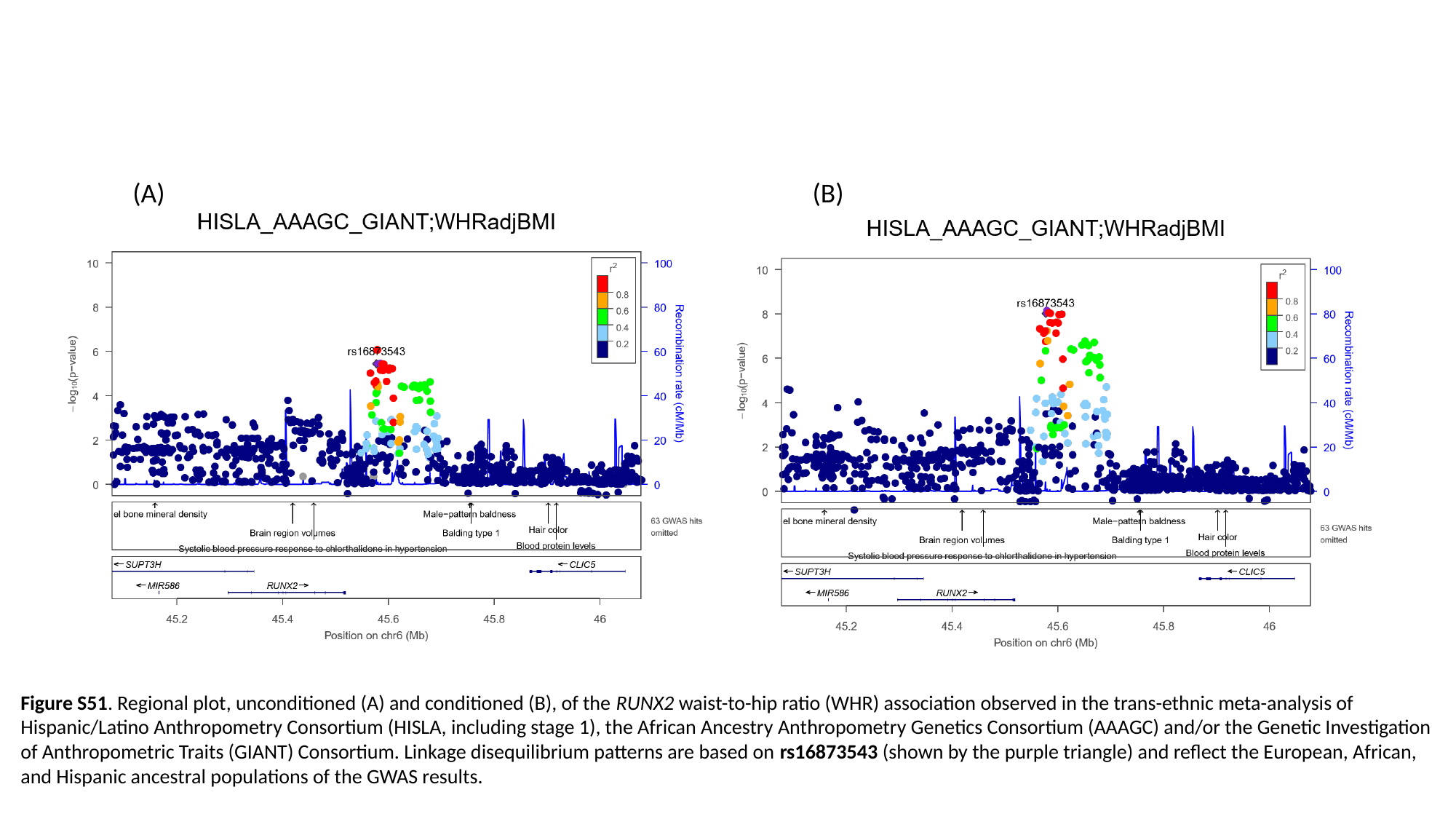

(A) (B)
Figure S51. Regional plot, unconditioned (A) and conditioned (B), of the RUNX2 waist-to-hip ratio (WHR) association observed in the trans-ethnic meta-analysis of Hispanic/Latino Anthropometry Consortium (HISLA, including stage 1), the African Ancestry Anthropometry Genetics Consortium (AAAGC) and/or the Genetic Investigation of Anthropometric Traits (GIANT) Consortium. Linkage disequilibrium patterns are based on rs16873543 (shown by the purple triangle) and reflect the European, African, and Hispanic ancestral populations of the GWAS results.

### Slide 52
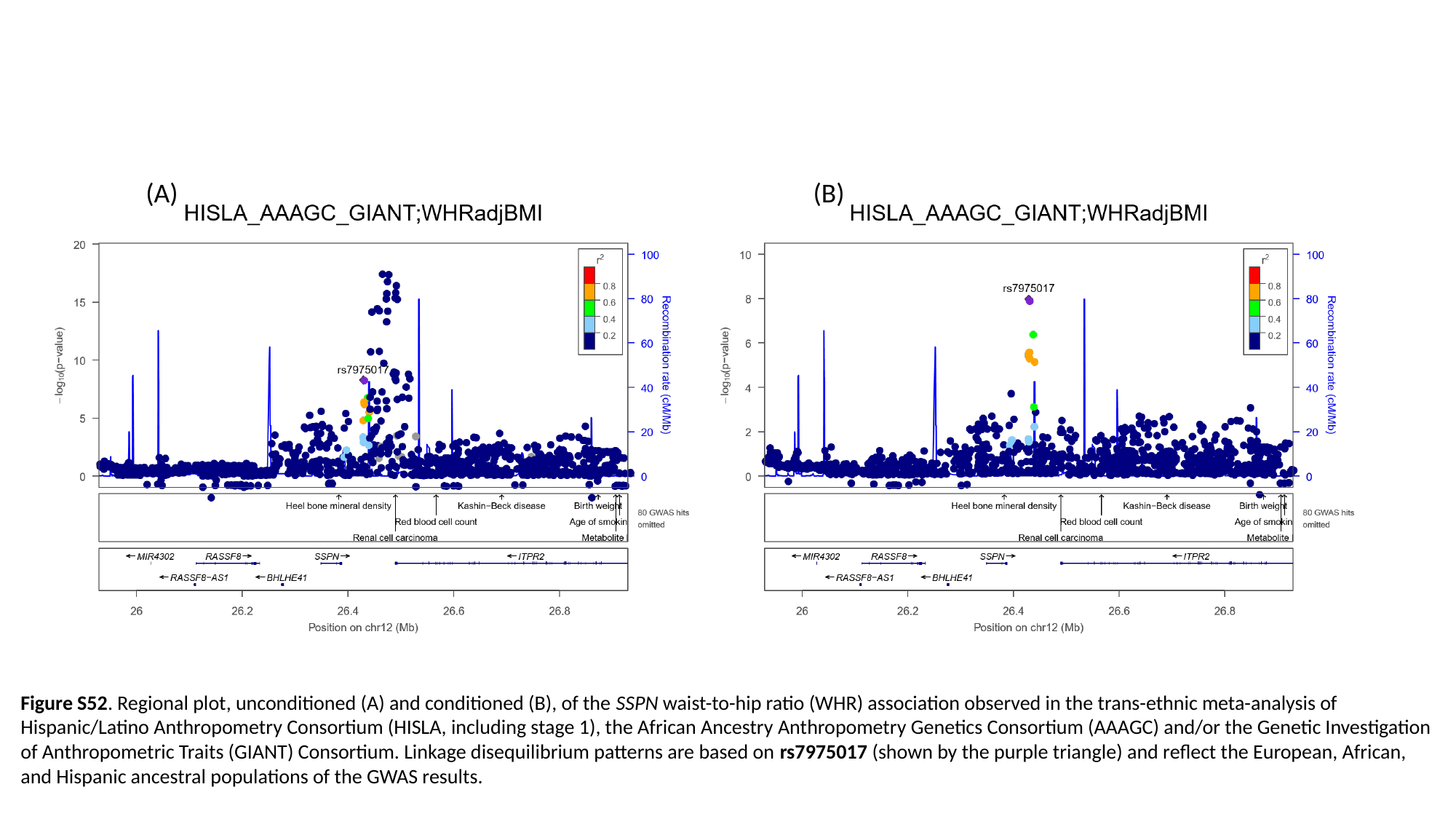

(A) (B)
Figure S52. Regional plot, unconditioned (A) and conditioned (B), of the SSPN waist-to-hip ratio (WHR) association observed in the trans-ethnic meta-analysis of Hispanic/Latino Anthropometry Consortium (HISLA, including stage 1), the African Ancestry Anthropometry Genetics Consortium (AAAGC) and/or the Genetic Investigation of Anthropometric Traits (GIANT) Consortium. Linkage disequilibrium patterns are based on rs7975017 (shown by the purple triangle) and reflect the European, African, and Hispanic ancestral populations of the GWAS results.

### Slide 53
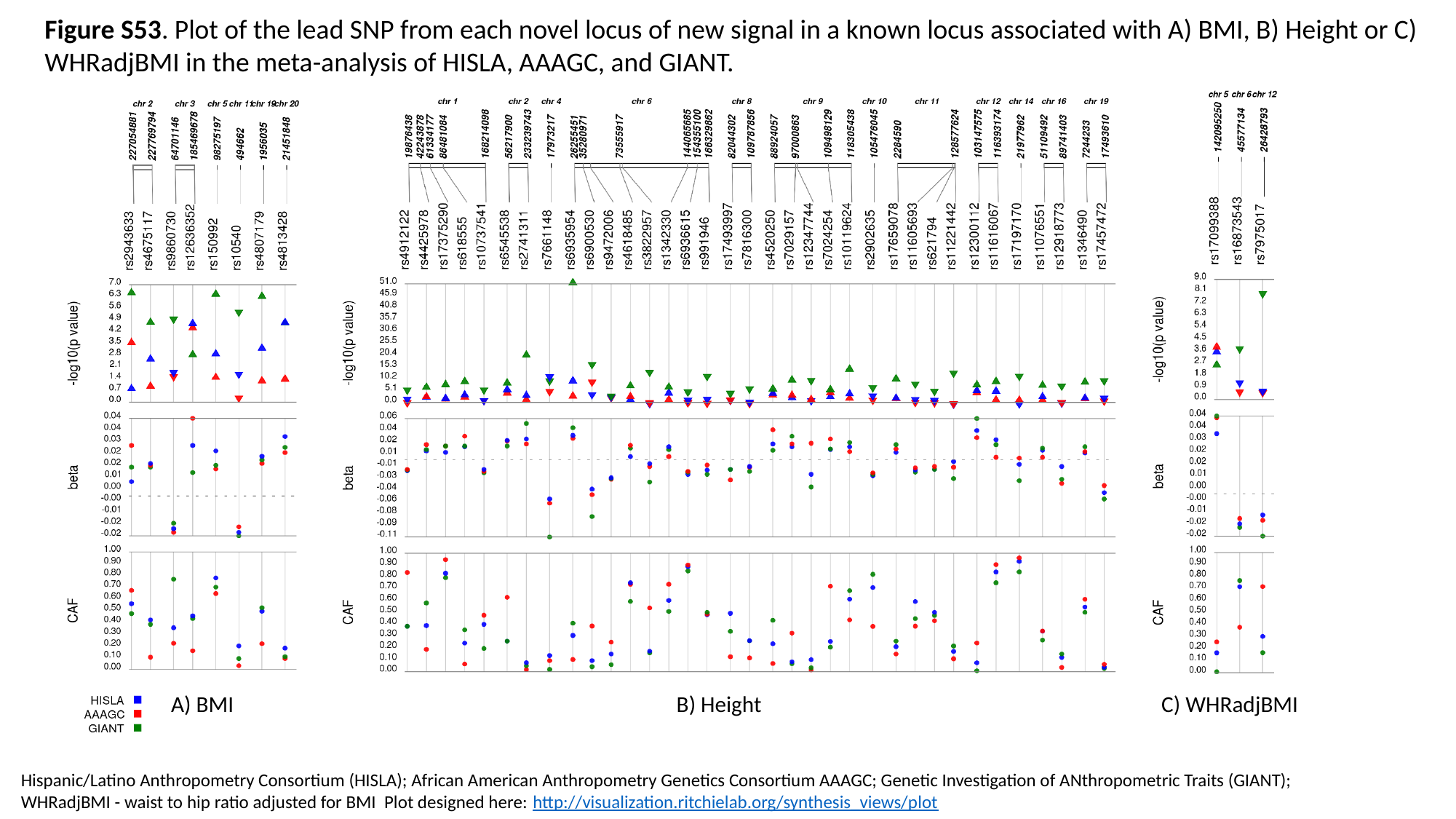

Figure S53. Plot of the lead SNP from each novel locus of new signal in a known locus associated with A) BMI, B) Height or C) WHRadjBMI in the meta-analysis of HISLA, AAAGC, and GIANT.
A) BMI			 B) Height	 C) WHRadjBMI
Hispanic/Latino Anthropometry Consortium (HISLA); African American Anthropometry Genetics Consortium AAAGC; Genetic Investigation of ANthropometric Traits (GIANT);
WHRadjBMI - waist to hip ratio adjusted for BMI Plot designed here: http://visualization.ritchielab.org/synthesis_views/plot

### Slide 54
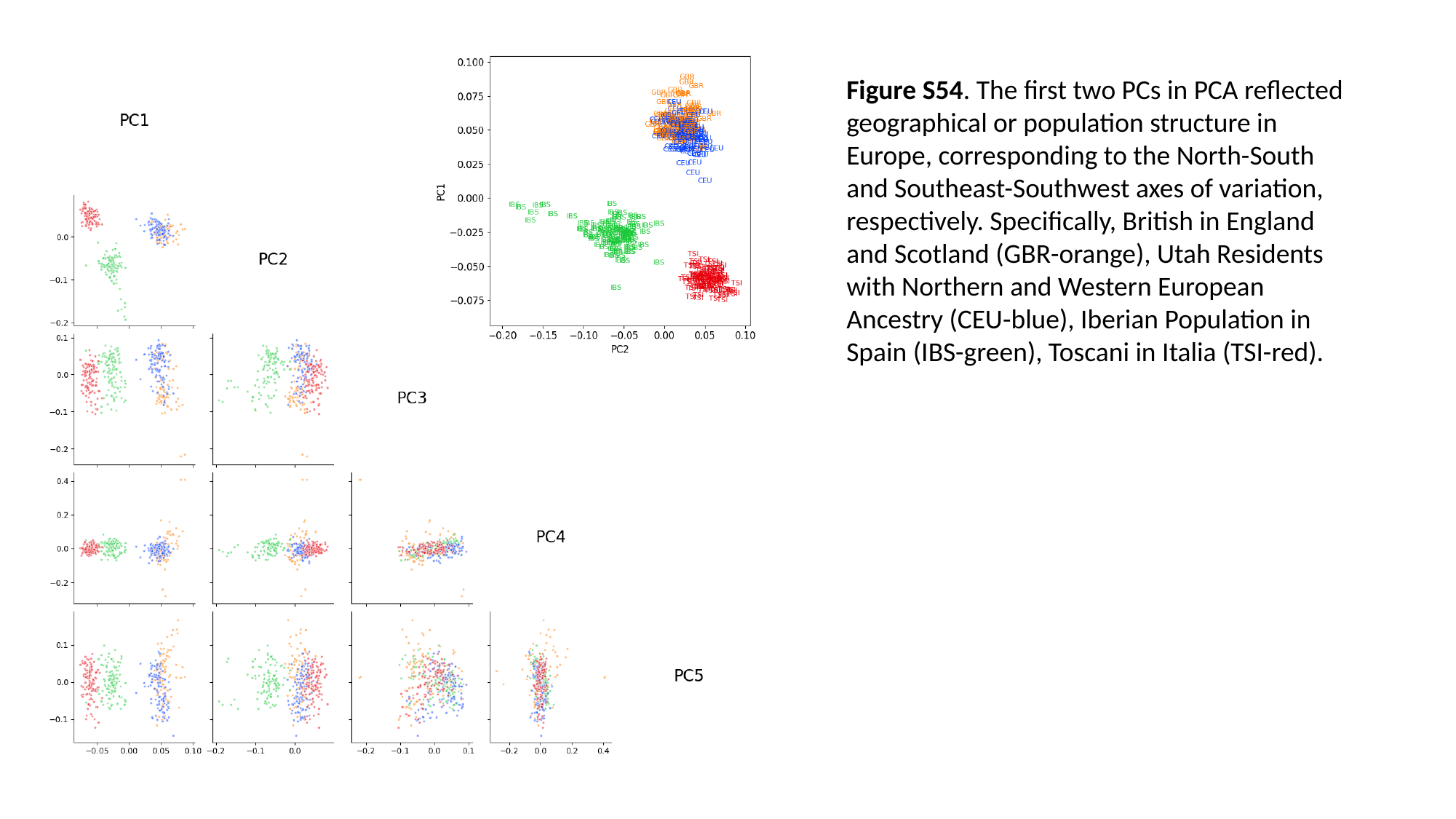

Figure S54. The first two PCs in PCA reflected geographical or population structure in Europe, corresponding to the North-South and Southeast-Southwest axes of variation, respectively. Specifically, British in England and Scotland (GBR-orange), Utah Residents with Northern and Western European Ancestry (CEU-blue), Iberian Population in Spain (IBS-green), Toscani in Italia (TSI-red).
